# Neighbouring modifications interfere with the detection of phosphorylated alpha-synuclein at Serine 129: Revisiting the specificity of pS129 antibodies

**DOI:** 10.1101/2022.03.30.486322

**Authors:** Hilal A. Lashuel, Anne-Laure Mahul-Mellier, Salvatore Novello, Ramanath Narayana Hegde, Yllza Jasiqi, Melek Firat Altay, Sonia Donzelli, Sean M. DeGuire, Ritwik Burai, Pedro Magalhães, Anass Chiki, Jonathan Ricci, Manel Boussouf, Ahmed Sadek, Erik Stoops, Christian Iseli, Nicolas Guex

**Affiliations:** Laboratory of Molecular and Chemical Biology of Neurodegeneration, School of Life Sciences, Brain Mind Institute, Ecole Polytechnique Fédérale de Lausanne (EPFL), CH-1015 Lausanne, Switzerland; Bioinformatics Competence Center, Ecole Polytechnique Fédérale de Lausanne, CH-1015 Lausanne, Switzerland; Bioinformatics Competence Center, University of Lausanne, CH-1015 Lausanne, Switzerland; ADx NeuroSciences, Technologiepark 94, Ghent, Belgium

**Keywords:** Synuclein, phosphorylation, antibodies, aggregation, post-translational modifications

## Abstract

Alpha-synuclein (aSyn) within Lewy bodies, Lewy neurites, and other pathological hallmarks of Parkinson’s disease and synucleinopathies have consistently been shown to accumulate in aggregated and phosphorylated forms of the protein, predominantly at Serine 129 (S129). Antibodies against phosphorylated S129 (pS129) have emerged as the primary tools to investigate, monitor, and quantify aSyn pathology in the brain and peripheral tissues. However, most of the antibodies and immunoassays aimed at detecting pS129-aSyn were developed based on the assumption that neighbouring post-translational modifications (PTMs) either do not co-occur with pS129 or do not influence its detection. Herein, we demonstrate that the co-occurrence of multiple pathology-associated C-terminal PTMs (e.g., phosphorylation at Tyrosine 125 or truncation at residue 133 or 135) differentially influences the detection of pS129-aSyn species by pS129-aSyn antibodies. These observations prompted us to systematically reassess the specificity of the most commonly used pS129 antibodies against monomeric and aggregated forms of pS129-aSyn in mouse brain slices, primary neurons, mammalian cells and seeding models of aSyn pathology formation. We identified two antibodies that are insensitive to pS129 neighbouring PTMs. However, consistent with previous reports, most pS129 antibodies showed cross-reactivity towards other proteins and often detected low and high molecular weight bands in aSyn knock-out samples that could be easily mistaken for monomeric or High Molecular Weight aggregates of aSyn. Our observations suggest that the pS129 antibodies do not capture the biochemical and morphological diversity of aSyn pathology. They also underscore the need for more specific pS129 antibodies, more thorough characterization and validation of existing antibodies, and the use of the appropriate protein standards and controls in future studies.

## Introduction

Phosphorylation of alpha-synuclein (aSyn) at serine 129 (pS129) has become the most commonly used marker of aSyn pathology formation and propagation in Parkinson’s disease (PD) and other synucleinopathies. Initial studies suggested that the majority (>90%) of aSyn in PD and dementia with Lewy bodies (DLB) brains is phosphorylated at S129^1^. Subsequent studies on the biochemical composition of Lewy bodies (LB) revealed that pS129 is the dominant aSyn post-translational modification (PTM) in LB^2,3^. Furthermore, an increase in pS129 levels in both cellular models and brains of animal models of synucleinopathies is usually associated with the appearance of aSyn aggregates, and the majority of pS129-aSyn species are typically found in the insoluble fractions of brains and cell extracts^4^. Finally, pS129 immunoreactivity has been observed in peripheral tissues and organs associated with non-motor symptoms of PD. These observations, combined with the development of a large number of antibodies against pS129-aSyn, have led to an increased reliance on pS129 antibodies as the primary tools for monitoring and quantifying aSyn pathology formation and spreading in human brains, peripheral tissues, and in cellular and animal models of synucleinopathies^5^.

This has prompted several studies to assess the sensitivity and specificity of pS129-aSyn antibodies. However, most of these studies have focused primarily on the cross-reactivity of the pS129-aSyn antibodies with other proteins^4,6,7^. Rutherford et al. reported that some of their in-house generated monoclonal pS129 antibodies cross-reacted with neurofilament subunits (NFL) phosphorylated at Serine 473 while others cross-reacted with other proteins^8^. Several pS129 antibodies have also been shown to cross-react with cellular nuclei under conditions where aSyn is either not expressed or could not be detected in nuclear fractions by Western blotting (WB)^8,9^. Delic et al. compared the specificity of four of the most commonly used pS129 monoclonal antibodies (clones EP1536Y, MJF-R13, 81A and pSyn#64) in brain slices and protein lysates from mice and rats^4^. Three of the four clones (MJF-R13, 81A, and pSyn#64) showed non-specific staining in tissues from aSyn knock-out (KO) mice and all four antibodies cross-reacted with other proteins. Consistent with previous studies by Rutherford et al., EP1536Y showed the highest sensitivity and specificity for detecting pS129-aSyn^4,8^. However, in a recent study Arlinghaus et al. evaluated the specificity of MJF-R13, pSyn#64, and EP1536Y, and reported that all three showed cross-reactivity and lack of specificity towards endogenous aSyn in wildtype (WT) or *SNCA* KO brain slices or primary cultures [by immunohistochemistry (IHC) or immunocytochemistry (ICC)]^10^. Recently, Fayyad et al. described the generation of three novel pS129-aSyn antibodies^5^. However, the specificity and affinity of these antibodies were assessed only against unmodified WT recombinant aSyn and unpurified *in vitro* Polo-Like Kinase 2 (PLK2) phosphorylated pS129-aSyn standards^5^.

Interestingly, most of the antibodies reported in the study by Fayyad et al. seem to detect primarily aSyn monomeric bands and did not detect the classical pattern of high molecular weight (HMW) aSyn bands usually detected in the insoluble fractions from PD brain homogenates^5^. This increasing reliance on pS129 antibodies and continuous generation of more antibodies necessitates the development of robust standardized pipelines for the characterization and validation of these antibodies before their use in PD and neurodegenerative disease research.

Increasing evidence from the biochemical characterization of aSyn pathology isolated from brain, peripheral tissues, and biological fluids by mass spectrometry show that aSyn is subjected to several modifications in the vicinity of serine 129, including phosphorylation at Tyrosine 125 (Y125)^11^, nitration at Y125, Y133, and Y136^12^ and truncations at residues 119, 122, 133 and 136^1,3,13,14^. aSyn species phosphorylated at S129 and Y125 have been shown to co-exist in pathological aggregates (Figure 1A). For example, several immunohistochemical studies examining the distribution of aSyn species in pathological aggregates have demonstrated that they are enriched in phosphorylated and C-terminally truncated aSyn species^3,13,15-19^. Recent mass spectrometry studies on LBs and insoluble brain-derived aSyn aggregates have identified several truncated aSyn species, including aSyn proteins that are truncated at both N- and C-terminal regions (Figure 1C)^1,13,14,20^. In the appendix, aSyn was predominantly a mixture of different species truncated at the N-terminus, C-terminus, or both termini. Thus far, all the studies on mapping aSyn PTMs have focused on identifying or characterizing one PTM at a time and global mapping of the distribution of aSyn species in biological samples rather than the distribution of PTMs that co-occur at the single-molecule level.

**Figure 1.**
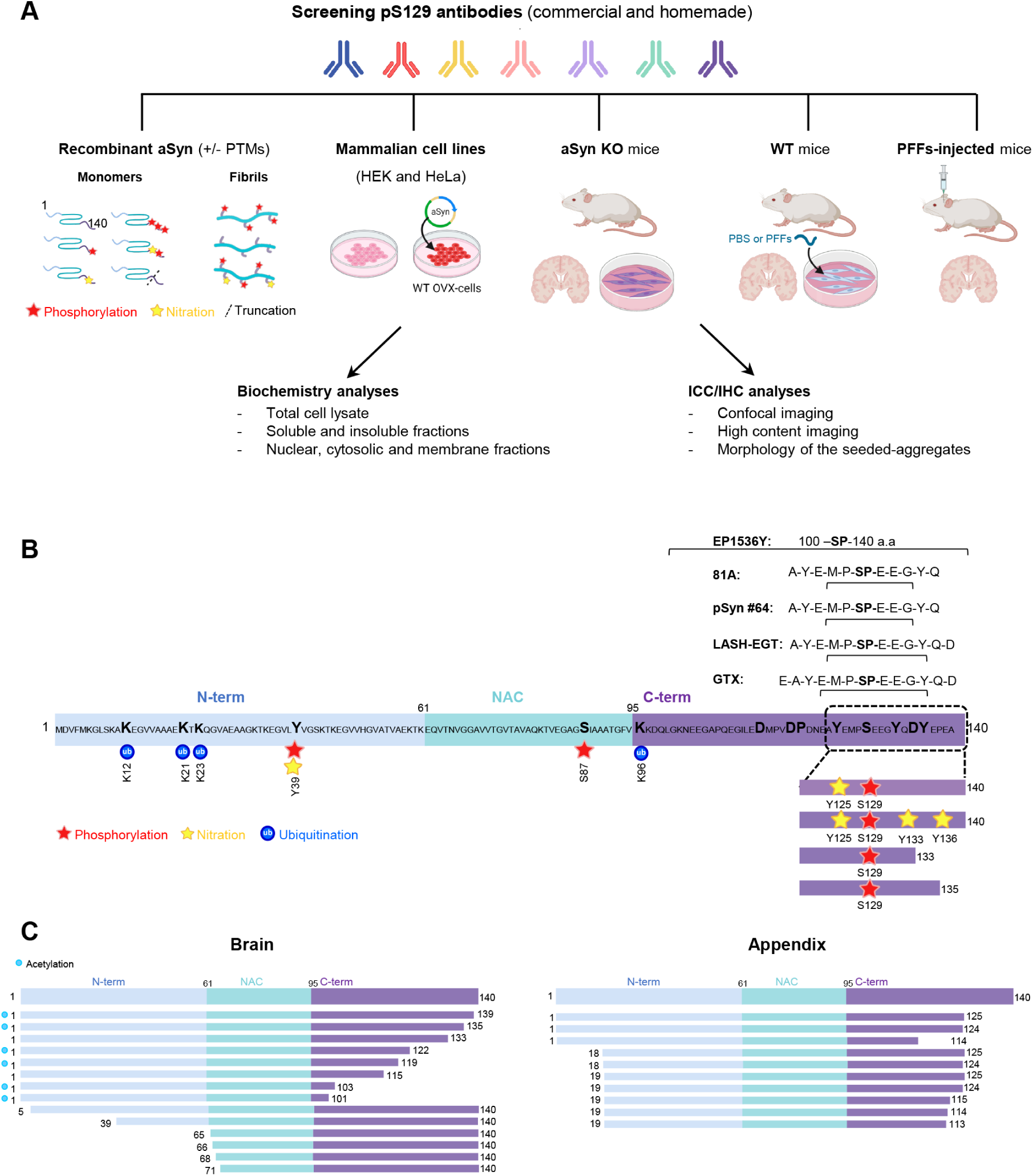
Assessment of the background and specificity of commercial and homemade pS129 antibodies *in vitro* and in cellular and animal models. **A**. The diagram depicts the workflow of our study to assess the background and specificity of commercial and homemade pS129 antibodies using recombinant aSyn monomers and fibrils (WT or carrying specific PTMS such as nitration, phosphorylation and/or truncation), mammalian cell lines overexpressing aSyn, aSyn KO primary neurons, WT neurons treated with aSyn PFFs or WT mice injected with aSyn PFFs. The background and specificity of the selected pS129 antibodies were evaluated by Western blot analyses or ICC or HTS combined to confocal imaging. Panel A was created using BioRender.com. **B**. The diagram depicts aSyn human sequence, the epitopes of each of the pS129 antibodies (whenever available), and the post-translational modifications that can occur concomitantly in the C-terminal region of aSyn. **C**. The diagram depicts the truncation that occurs in the different regions of aSyn in human brains and the appendix.

We hypothesized that the presence of multiple PTMs in the C-terminus could significantly influence or mask the detection of pS129-aSyn^1,3,21-24^. To test this hypothesis, we systematically assessed, for the first time, the effect of the co-occurrence of pathology-associated C-terminal PTMs on the detection of pS129-aSyn. This was achieved by leveraging our protein synthesis and recombinant expression platforms, which enabled the generation of a library of highly pure and site-specifically modified recombinant and semisynthetic full-length aSyn proteins phosphorylated at S129 and bearing different combinations of other disease-associated PTMs, including tyrosine 125 phosphorylation/nitration and C-terminal truncations at 133 and 135 (Figure 1B). However, given the large number of available pS129 antibodies, it was not possible to assess them all. Therefore, for this study, we conducted a thorough review of the aSyn literature and selected the five most commonly used commercially available antibodies across different methods. These include two mouse monoclonal antibodies (pSyn#64 and 81A) and two monoclonal rabbit antibodies: EP1536Y, MJF-R13 and the polyclonal rabbit GTX) (Figure S1). In addition, we included a homemade rabbit antibody (LASH-EGT) (Figure 1B). While working with these antibodies, we observed, as reported previously^4,8,25^, that they cross-react with other proteins and that some of the non-specific signals could be easily mistaken for aSyn signals. This prompted us to expand the pipeline for assessing the specificity of pS129 antibodies in mouse brain tissues, in primary neurons and mammalian cells lacking or expressing aSyn. Furthermore, we investigated the differential ability of these antibodies to capture the diversity of aSyn aggregates in neuronal and *in vivo* models of aSyn seeding and pathology spreading. Our results show that several C-terminal PTMs found in aSyn pathological aggregates eliminate or mask the detection of pathological aggregates by pS129 antibodies. These observations suggest that pS129 antibodies cannot capture the biochemical diversity of pathological aSyn species, and using these antibodies might result in underestimating aSyn pathology or selective preference for pathological aggregates composed of the full-length aSyn protein. Furthermore, we demonstrate that these antibodies exhibit differential cross-reactivity towards a large number of proteins of various sizes and subcellular localization. Finally, we searched the human and mouse proteome and generated a comprehensive list of candidate proteins that could cross-react with pS129 antibodies. Taken together, our findings underscore the critical importance of developing more specific pS129 antibodies, exercising caution in interpreting results from experiments relying mainly on the use of pS129 antibodies, and including the appropriate controls in experiments using these antibodies to investigate aSyn biology or mechanisms of pathology formation and spreading. Specific recommendations for developing, validating, and characterize more specific aSyn antibodies are presented and discussed.

## Results

### All pS129 antibodies tested robustly detected phosphorylated aSyn (pS129), but not the unmodified protein

As the first step in our study, we assessed the specificity of the pS129 antibodies against highly pure aSyn protein that is site-specifically phosphorylated at S129 only and unmodified WT aSyn (Figure S2). None of the six antibodies (pSyn#64, EP1536Y, MJF-R13, 81A, GTX, and LASH-EGT) tested (Figure S3B) detected the unmodified protein and showed robust signals for pS129-aSyn (Figure S4A). Next, we assessed the specificity of three pS129 antibodies (pSyn#64, MJF-R13 and LASH-EGT) against aSyn proteins that are site-specifically phosphorylated at residue Y125 (Figures 2A and S2A and B) or other residues such as Y39 or S87 (Figure S4B and C). None of the pS129 antibodies tested could detect any phosphorylated aSyn where S129 was not phosphorylated. These findings demonstrate that the selected pS129 antibodies are highly specific for pS129.

**Figure 2.**
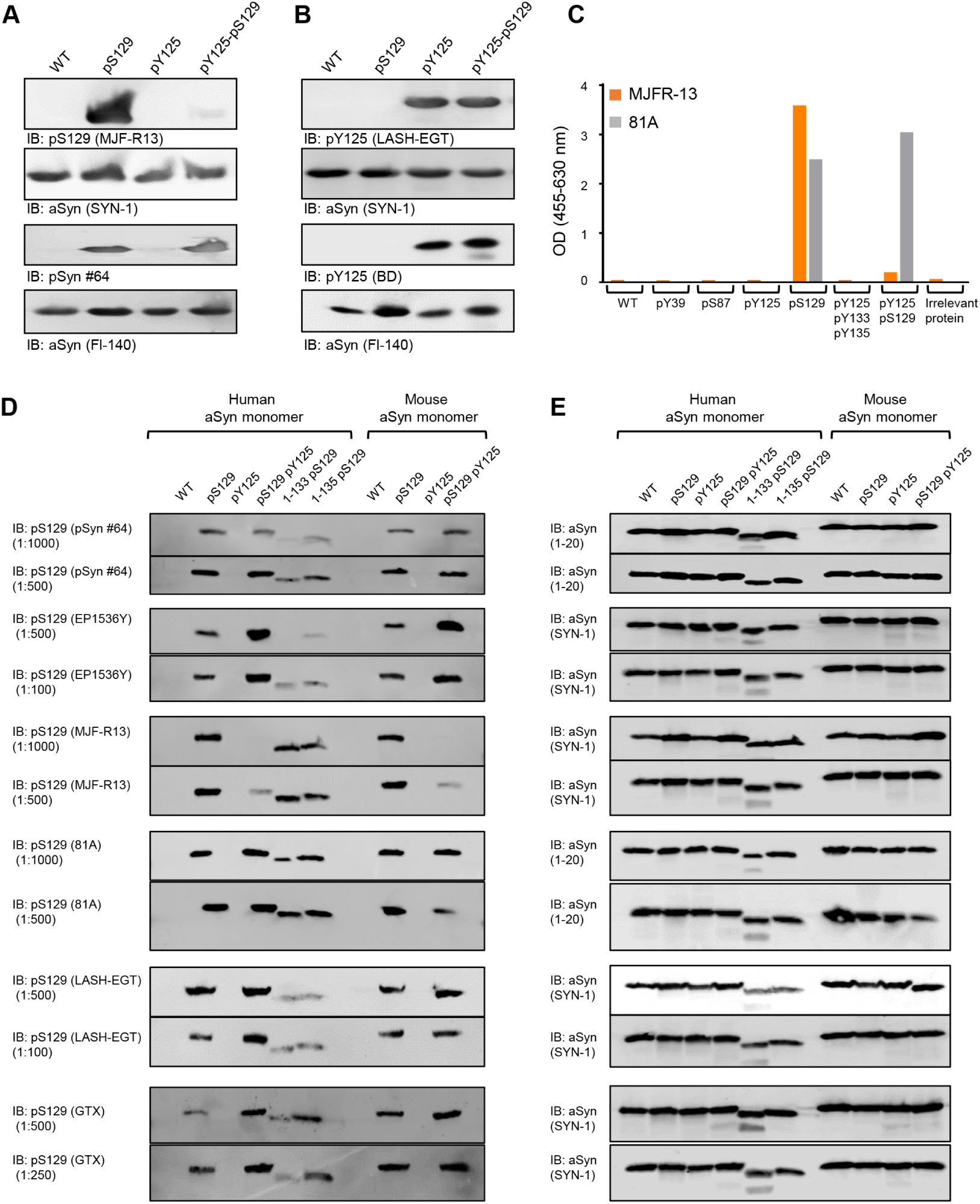
PTMs nearby S129 residue interferes with the capacity of some pS129 antibodies to detect pS129-aSyn. The capacity of human or mouse recombinant monomeric aSyn (WT or modified by specific PTMs) to be detected by pS129 commercial antibodies (pSyn#64, MJF-R13, 81A, EP1536Y and GTX) or the LASH-EGT homemade pS129 antibody was assessed by WB (**A-B, D**) or ELISA (**C**). **A-B**. Forty nanograms of human recombinant monomeric aSyn unphosphorylated (WT) or phosphorylated at the S129 residue (pS129) or di-phosphorylated (pY125/pS129) were detected by WB using total aSyn antibodies (SYN-1 or Fl-140) combined with either (**A**) commercial (MJF-R13 or pSyn#64) and homemade (LASH-EGT) pS129 antibodies or (**B**) commercial pY125 antibody (BD) or the homemade antibody LASH-EGT-pY125. **C**. ELISA confirmed that MJF-R13, but not 81A, is no longer able to detect pS129-aSyn when aSyn was di-phosphorylated (pY125/pS129). **D-E**. One-hundred nanograms of unmodified aSyn (WT) or phosphorylated aSyn at S129 and/or Y125 residues (respectively named pS129, pY125 or pS129 pY125) full length or truncated after residues 133 (1-133 pS129) or 135 (1-133 pS129) were detected by WB using pS129 antibodies (**D**) or total aSyn antibodies (SYN-1 or LASH-EGT 1-20) (**E**).

### The presence of additional PTMs close to the S129 interfere with the detection of pS129

The majority of pS129 antibodies were developed based on the assumption that neighbouring modifications either do not co-occur with pS129 or do not influence its detection. Therefore, most peptide antigens used to raise antibodies were comprised of 3–6 residues on each side of S129 (Figures 1B and S3B) and were modified only at S129. We hypothesized that the co-occurrence of pS129 and other PTMs in the C-terminal region spanning residues 123–135 could interfere with pS129 detection. To test this hypothesis, we first assessed whether phosphorylation of tyrosine 125 (Y125), which has been observed in pS129 immunoreactive LBs^22^, could influence the detection of pS129. Towards this goal, we generated semi-synthetic human full-length aSyn proteins site-specifically phosphorylated on residue S129 (pS129) or residue Y125 (pY125) or di-phosphorylated (pY125/pS129) (Figure S2C). The proteins were produced as described previously^26-28^.

Initially, we assessed the ability of two of the most commonly used pS129 antibodies (MJF-R13 and pSyn#64) to detect singly (pS129) and diphosphorylated aSyn pY125/pS129 by WB. To our surprise, the MJF-R13 detected only singly phosphorylated pS129, whereas the pSyn#64 antibody detected both singly pS129 and di-phosphorylated pY125/pS129-aSyn proteins (Figure 2A-B). Interestingly, the two pY125 antibodies tested detected both pY125 and pY125/pS129-aSyn (Figure 2B). These findings confirmed our hypothesis and suggested that the presence of other C-terminal PTMs, e.g., pY125, could interfere with the detection of pS129 by some antibodies raised against this PTM. However, pS129 does not seem to influence the detection of pY125. These findings were validated using an enzyme-linked immunosorbent assay (ELISA) where two pS129 antibodies (MJF-R13 and 81A) were evaluated using a library of aSyn proteins containing single (pY39, pS87, pY125, pS129) or multiple C-terminal phosphorylation sites (pY125/pS129 and pY125/pS133/pY136). As shown in Figure 2C, none of the pS129 antibodies detected aSyn proteins phosphorylated at other residues (pY39, pS87, pS129), and only the 81A antibody detected aSyn that is phosphorylated at both pY125 and pS129. These observations prompted us to conduct a more systematic investigation of the impact of other disease-associated PTMs on the detection of phosphorylated aSyn by pS129 antibodies (Figure 2C).

Several studies have confirmed that some of the aSyn found in LBs in human brain tissue is truncated at several residues, including Tyrosine 133 (Y133) or Aspartic acid 135 (D135)^1,14,29^ (Figure 1C). Recent studies have also reported that aSyn is cleaved at Y125, S129, Y133, and D135 in preformed fibrils (PFFs) seeded neuroblastoma cell lines^30^. Similarly, we and others have shown that human and mouse aSyn PFFs undergo multiple C-terminal cleavages, including E114, D115, D119, D121, S122, Y125, and D135^30,31^. Therefore, we sought to assess the ability of the pS129 antibodies to detect aSyn species that are both phosphorylated at S129 and truncated at the far C-terminus, at residues 133 or 135. To produce aSyn bearing both modifications, monomeric full-length or truncated (1–133 or 1–135) aSyn were phosphorylated specifically at S129 using the Polo-Like kinase 3 (PLK3) *in vitro* as previously described^27,28^. The homogeneously phosphorylated proteins were purified by RP-HPLC, and the final purity of the proteins was confirmed by SDS-PAGE, UPLC and LC-MS analyses (Figure S2D). With all these proteins in hand, we screened by WB the five most commonly used pS129 antibodies in the PD field (pSyn#64, MJF-R13, 81A, EP1536Y, and GTX, Figure S1), in addition to our pS129 homemade antibody (LASH-EGT).

As shown in Figures 2D and E, all six antibodies evaluated in this study detected the singly phosphorylated (pS129) but not the unmodified, human, and mouse aSyn proteins. Interestingly, the aSyn proteins bearing pS129 and truncations at 133 or 135 were barely detected by three (pSyn#64, EP1536Y, and LASH-EGT) out of the six antibodies (Figure 2D and E). Detection of pS129 by MJF-R13 and 81A was not affected by truncation at 133 or 135, whereas the GTX antibody detected 1-135/pS129 but not 1-133/pS129-aSyn. This data confirmed that cleavages in the sequences flanking S129 residue (e.g., truncations at 114, 119, 115, 120, 121, 129, 133, or 135)^1,3,13,16,31^ could either abolish (e.g., truncations at 114, 119, 120 or 121) or interfere with (e.g., truncations at 129, 133 or 135) pS129 detection by all or some pS129 antibodies, respectively. Among the six antibodies tested, only MJF-R13 completely lost its ability to detect di-phosphorylated human and mouse aSyn at S129 and Y125 (pY125/pS129) while efficiently detecting the singly phosphorylated aSyn at S129 or 1-133/pS129-aSyn and 1-135/pS129-aSyn (Figure 2D). The EP1536Y antibody detected di-phosphorylated (pY125/pS129) aSyn (human and mouse) but failed to detect pS129-aSyn truncated after residues 133 or 135. Overall, only the 81A clone was insensitive to neighbouring PTMs and detected pS129-aSyn that is also phosphorylated at Y125 or truncated at residue 133 or 135 (Figure 2D). Together, these results show that the co-occurrence of pS129 and other C-terminal modifications interfere with pS129-aSyn detection but show that it is possible to develop antibodies capable of capturing the diversity pS129-aSyn species *in vitro*.

### Not all pS129 antibodies detect aSyn fibrils that are subjected to both C-terminal nitration (Y125/Y133/Y136) and phosphorylation (S129)

Given that C-terminal aSyn PTMs are more abundant in the aggregated state of aSyn, we next sought to determine if neighbouring modifications could influence the detection of aSyn fibrils phosphorylated at S129. We first generated aSyn fibrils site-specifically phosphorylated at S129 (Figure 3A). This was achieved by in vitro phosphorylation of aSyn PFFs using PLK3 (Figure S2C). Next, fibrils nitrated at all tyrosine residues (nY39/nY125/nY133/nY136) were also generated by chemical nitration of fibrils using Tetranitromethane (TNM), as described previously^32^ (Figure S5). Finally, fibrils that were both phosphorylated and nitrated (pS129/nY125/nY133/nY136) were also generated by treatment of pS129-aSyn fibril with TNM (Figure S5). In all cases, the extent of fibril modification was verified by mass spectrometry (Figure S5A).

**Figure 3.**
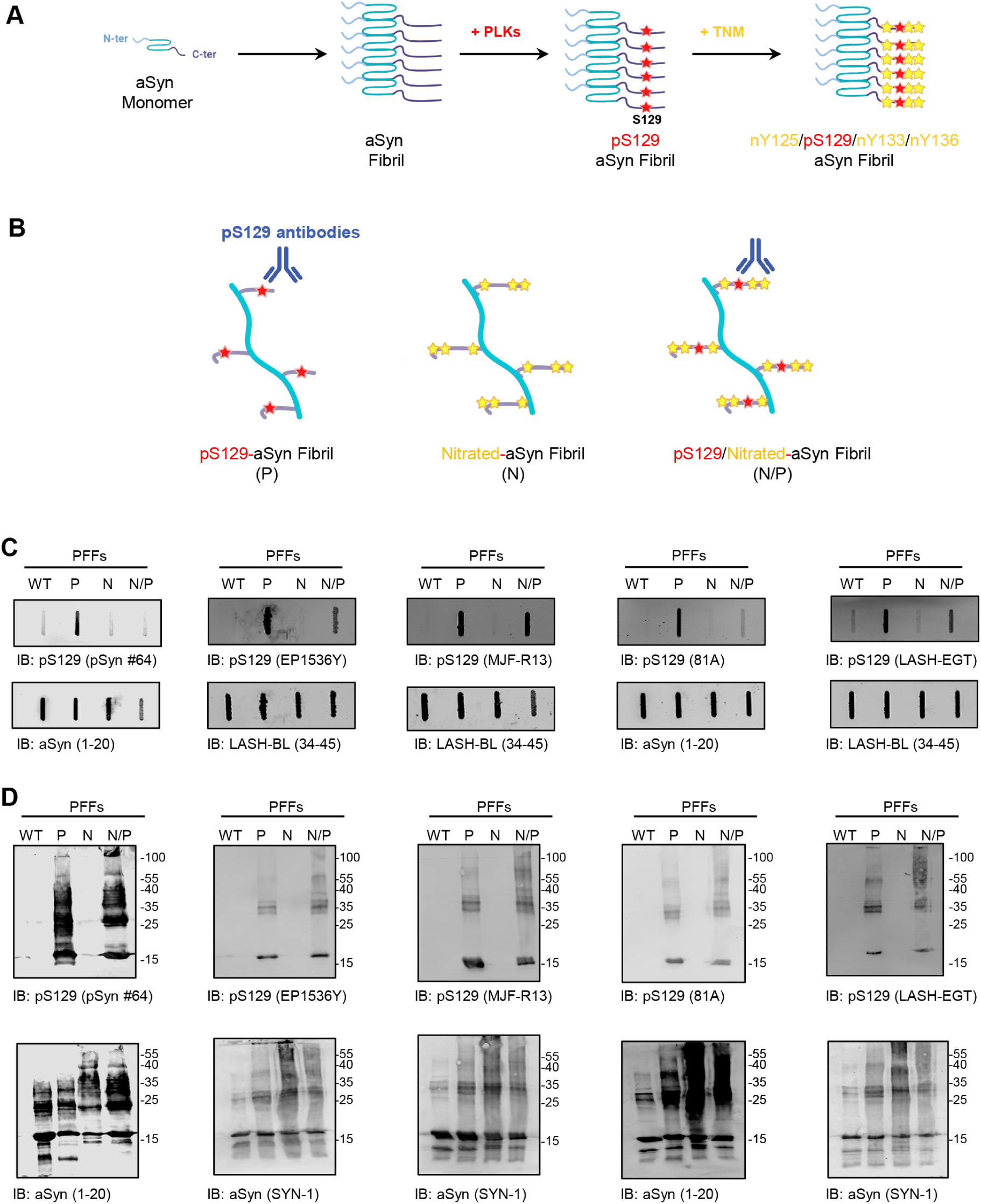
Nitration of aSyn does not impair the capacity of pS129 antibodies to detect pS129 fibrils. **A**. Preparation of aSyn fibrils carrying both phosphorylation and nitration modifications. aSyn fibrils were first site-specifically phosphorylated at S129 residue by being incubated with PLK3. Afterwards, TNM was added to induce the nitration of the phosphorylated aSyn fibrils. **B**. The capacity of aSyn fibrils site-specifically phosphorylated at S129 and/or nitrated to be detected by the pS129 commercial antibodies (pSyn#64, MJF-R13, 81A and EP1536Y) or the LASH-EGT homemade pS129 antibody was assessed by slot blot (**C**) and WB (**D**) analyses. Panels **A** and **B** were created using BioRender.com. **C-D**. Thirty-six nanograms of aSyn fibrils (PFFs) that were unmodified (WT), phosphorylated (P), nitrated (N), or nitrated/phosphorylated (N/P) were detected by slot blot (**C**) and WB (**D**) using pS129 antibodies (top panel) in combination with total aSyn antibodies (1–20 and 34–45, bottom panel).

The pS129 antibodies pSyn#64, EP1536Y, MFJ-R13, 81A, and LASH-EGT, were then screened by dot blot and WB using pS129-aSyn monomers, pS129-aSyn fibrils, and unmodified aSyn fibrils as a negative control. As shown in Figures 3B and C, all five antibodies detected pS129-aSyn fibrils, but not the unmodified aSyn fibrils. Nitration of the three C-terminal tyrosine residues abolished the ability of the pSyn#64 and 81A antibodies to detect pS129 in pS129/nY125/nY133/nY136-aSyn (Figure 3B-C). In contrast, these modifications did not significantly affect the detection using MJF-R13 and LASH-EGT. The EP1536Y antibody showed reduced signal for pS129/nY125/nY133/nY136-aSyn fibrils compared to pS129-aSyn fibrils. These findings demonstrate that some pS129 antibodies may fail to capture aSyn fibrils that are both phosphorylated and nitrated at the C-terminal domain of the protein. Interestingly, all five antibodies showed high specificity and immunoreactivity towards both monomeric and high molecular weight bands of both pS129/nY125/nY133/nY136-aSyn fibrils and pS129-aSyn fibrils in denaturing conditions (WB analyses) (Figure 3C). These results suggest that treatment with denaturing agents helps reveal the pS129 signal masked by the presence of some neighbouring PTMs, e.g., nitration. They also demonstrate that antibodies that were not affected by neighbouring phosphorylation or truncation events (e.g., 81A) are sensitive to other modifications (e.g., nitration). These observations underscore the importance of always using multiple pS129 antibodies with well-characterized specificity towards the diversity of modified forms of the protein.

### C-terminal truncations affect the detection of aSyn pS129 level in primary neurons

Next, we screened the ability of the antibodies to detect pS129-aSyn aggregates in a neuronal seeding model^33,34^ where newly formed aggregates were shown to exhibit a PTM profile^31^ resembling that of LBs in human brains^1^. Namely, phosphorylation at S129, C-terminal cleavage at multiple sites (103, 114, 119, 121, 129 and 135), and ubiquitination at multiple lysine residues^31^ (Figure 1B). We compared the ability of the six antibodies (Figure 1C) to detect pS129 in neurons treated with aSyn PFFs^34^. After 10 days of treatment, neurons were fixed, and ICC was performed. Confocal imaging showed that the six antibodies detected S129 phosphorylated aSyn inside the seeded-aggregates in MAP2 positive neurons (Figure 4A). Quantification of images acquired by a high-throughput wide-field cell imaging system (HTS) robustly demonstrated that the pSyn#64, 81A, MJF-R13, and GTX antibodies were the most efficient in detecting pS129-aSyn seeded-aggregates (Figures 4B and C and S6A). These findings suggest that the pS129 antibodies differentially detect aSyn fibrillar aggregates in neurons. Two of the four antibodies, pSyn#64 and MJFR-13, showed nuclear background signal in PBS treated neurons (*see below*).

**Figure 4.**
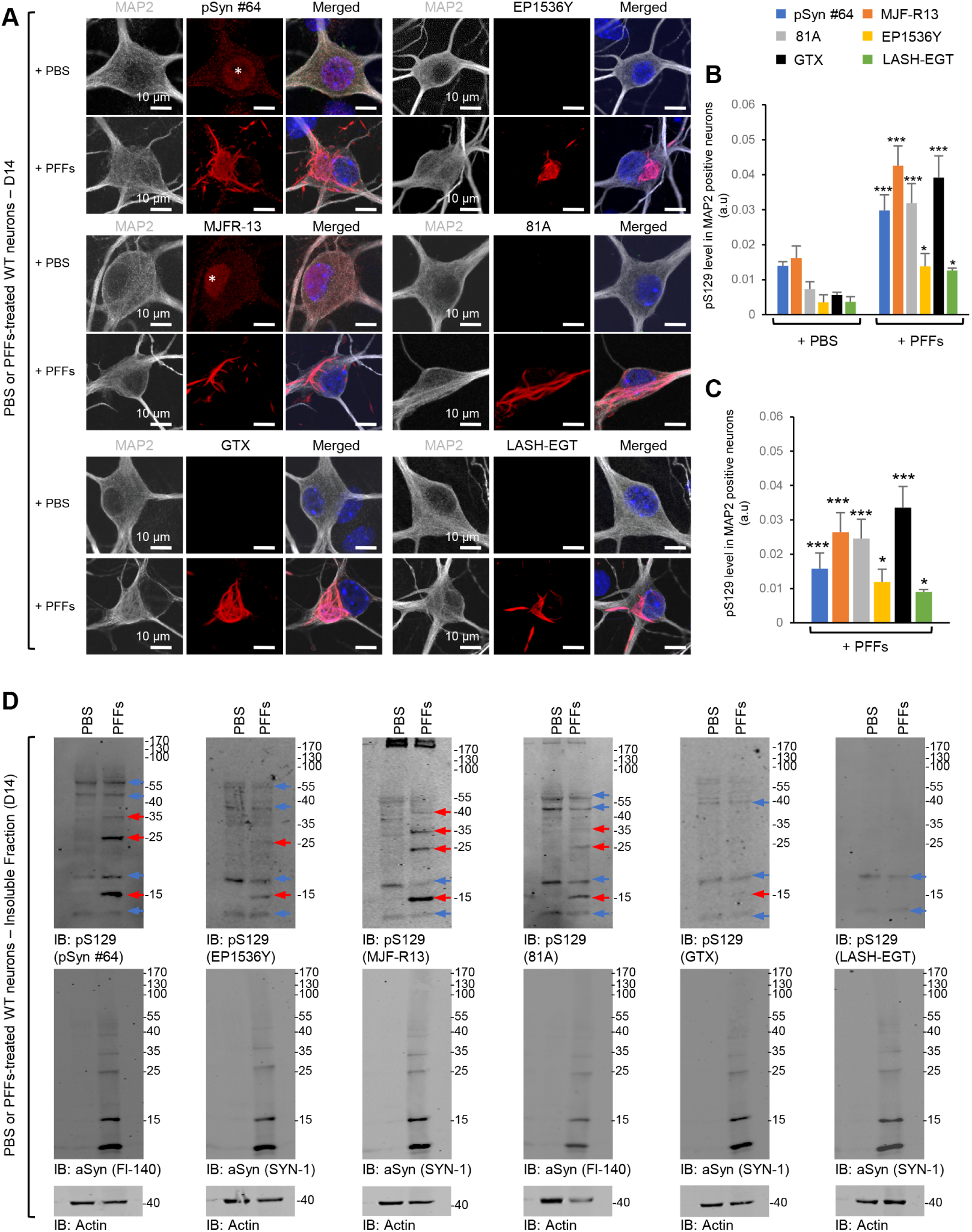
Assessment of pS129 antibodies detection in the PFFs-seeded WT primary hippocampal neurons. After 14 days of treatment with 70 nM of mouse aSyn PFFs or PBS buffer (negative control), primary hippocampal neurons were fixed or lysed, and ICC (**A-C**) or WB (**D**) analyses were performed using pS129 antibodies (pSyn#64, MJF-R13, 81A, EP1536Y and GTX or LASH-EGT). **A-C**. Newly formed fibrils were detected by confocal imaging (**A**) and quantified by a high-throughput wide-field cell imaging system as previously described^34^ (**B-C**). Neurons were counterstained with MAP2 antibody and the nucleus with DAPI staining. Scale bar = 10 μM. **B**. The left-hand side part of the histogram shows the background of each pS129 antibody in PBS-treated primary neurons, while the right-hand side part of the histogram shows the pS129 level detected by the pS129 antibodies in PFFs-treated primary neurons. **C**. pS129 level in PFFs-treated primary neurons was re-evaluated after subtracting the pS129 background level from the PBS-treated neurons. The graphs represent the mean +/- SD of three independent experiments. p<0.01=*, p<0.0001=*** (ANOVA followed by Tukey HSD post-hoc test, PBS vs. PFFs treated neurons). **D**. WB analyses of the insoluble fractions of the PBS- and PFF-treated neurons. Membranes were then counterstained by total aSyn antibodies (SYN-1 or Fl-140), and actin was used as a loading control. The red arrows indicate the pS129-aSyn positive bands. The blue arrows indicate the non-specific bands.

Next, we compared the specificity of the six antibodies towards aSyn aggregates isolated from the neuronal seeding model by WB analysis (Figures 4D and S6B). The GTX and LASH-EGT antibodies did not show any pS129 signal. The pSyn#64, EP1536Y, MJF-R13, and 81A antibodies detected several non-specific bands in the control samples from neurons treated with PBS. Two specific bands occur just above and below where aSyn typically runs, as evidenced by comparing the samples from PBS and PFF-treated neurons (Figure 4D). Furthermore, these antibodies also detect higher molecular weight bands between 35–55 kDa (pSyn#64, EP1536Y, MJF-R13, and 81A) and above 170 kDa (MJF-R13 and 81A). In the case of the samples from PFF-treated neurons, all four antibodies detect the 15 kDa band and the non-specific 17 kDa and additional higher bands that appear to be aSyn specific as they were not detected in the samples from PBS neurons (Figure 4D).

Our findings underscore the importance of including the appropriate control samples to ensure accurate assignment of the aSyn bands by WBs. The aSyn bands between 25 and 40 KDa are frequently referred to as HMW or oligomeric aSyn species. However, previous studies from our lab^31^ and others^1,2,35^ suggest that they represent ubiquitinated forms of aSyn, which are breakdown products from HMW aggregates, including fibrils.

### Not all the pS129 antibodies can capture the morphological diversity of the pS129-aSyn seeded-aggregates

We have recently shown that the nature of the PFF seeds is a strong determinant of the morphological diversity of aSyn aggregates we detect in our seeding neuronal model of LB-like inclusion formation. When hippocampal or cortical primary neurons are treated with mouse WT PFF, newly formed fibrils appear mainly as filamentous- and ribbon-like aggregates or LB-like inclusions at D21^34,36^ (Figure S7). In contrast, when hippocampal or cortical primary neurons are treated with PFFs derived from the newly identified human aSyn mutant E83Q, we observe a greater morphological diversity, as evident by the detection of up to six distinct morphologies, namely the nuclear-, the granular-, the filamentous and granular-, the filamentous- and the filamentous-like seeded aggregates as well as the dense- and the ring-like LBs^36^ (Figure 6). Therefore, we used this model to evaluate the ability of the top four pS129 antibodies (MJF-R13, GTX, 81A, or pSyn#64) to capture the morphological diversity of aSyn pathology in primary neurons. The LASH-EGT and the EP1536Y antibodies were excluded from the screen for the following reasons: 1) the LASH-EGT antibody due to its poor capacity to recognize aSyn pathology in PFF-treated neurons (Figures 4B and C); and 2) because the new EP1536Y antibody batches we recently acquired were no longer suitable for ICC, as acknowledge by Abcam by removing the ICC recommendation from their datasheet (https://www.abcam.com/alpha-synuclein-phospho-s129-antibody-ep1536y-ab51253.html).

**Figure 5.**
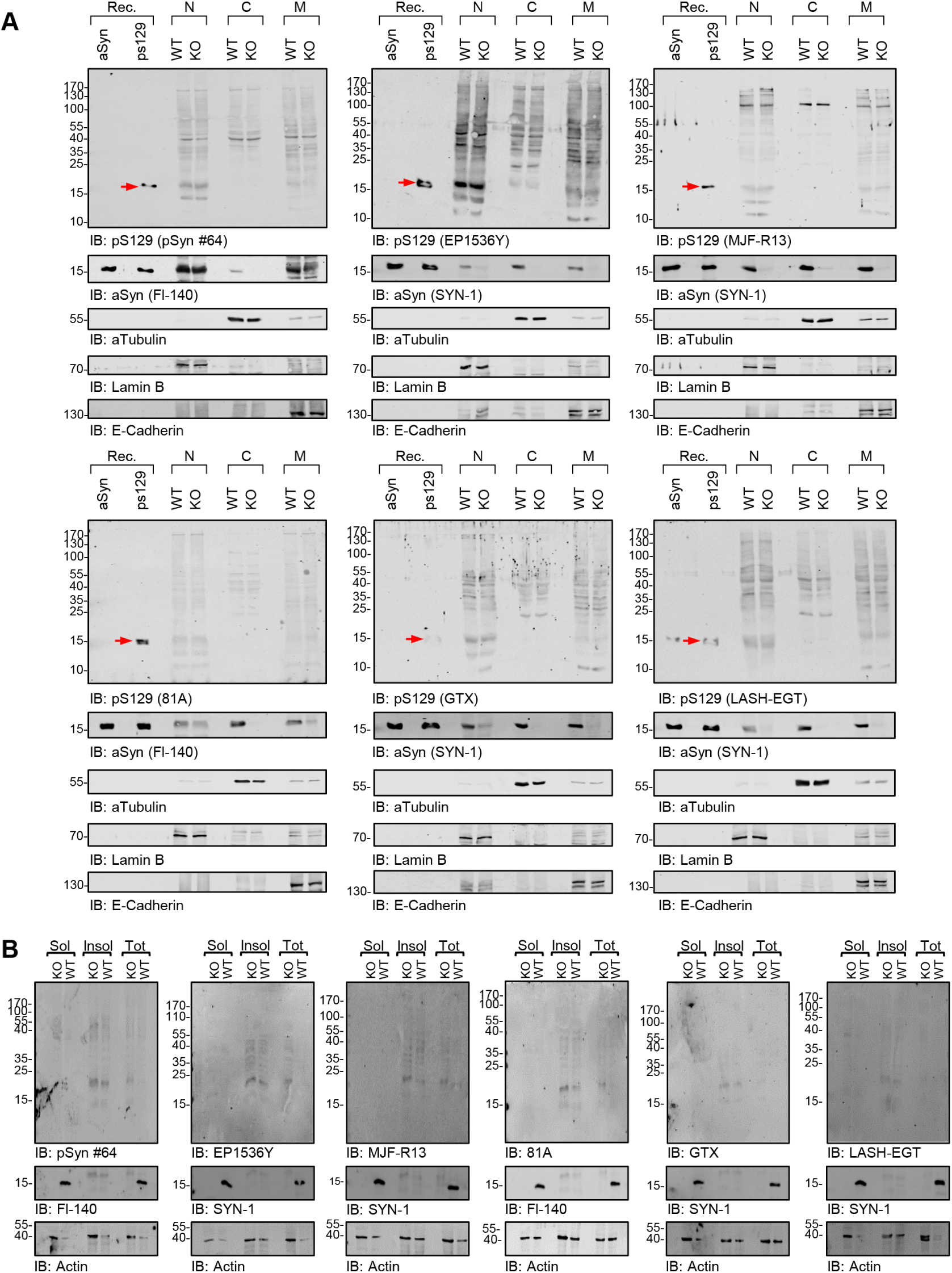
Assessment of pS129 antibodies detection in the nuclear, cytosolic, and membranous fractions of aSyn KO and WT primary hippocampal neurons by WB. **A-B**. Sequential biochemical extraction was performed on aSyn KO and WT primary hippocampal neurons cultured for 14 days (DIV14), and the pS129 level was assessed in the nuclear (N), cytosolic (C), and membrane (M) fractions (**A**) or in the soluble (sol) and insoluble (insol) fractions (**B**) by WB using the six pS129 antibodies (pSyn#64, MJF-R13, 81a, EP1536Y, GTX and LASH-EGT). Forty nanograms of recombinant aSyn WT or phosphorylated at residue S129 (pS129, indicated by the red arrow) were used as positive controls. The pS129 antibodies were used at a dilution of 1/1000. Membranes were then counterstained by total aSyn antibodies (SYN-1 or Fl-140). aTubulin, Laminin B, and E-Cadherin were used as loading control for the cytosolic, nuclear, and membrane fractions, respectively (**A**). Actin was used as a loading control for the soluble and insoluble fractions (**B**).

**Figure 6.**
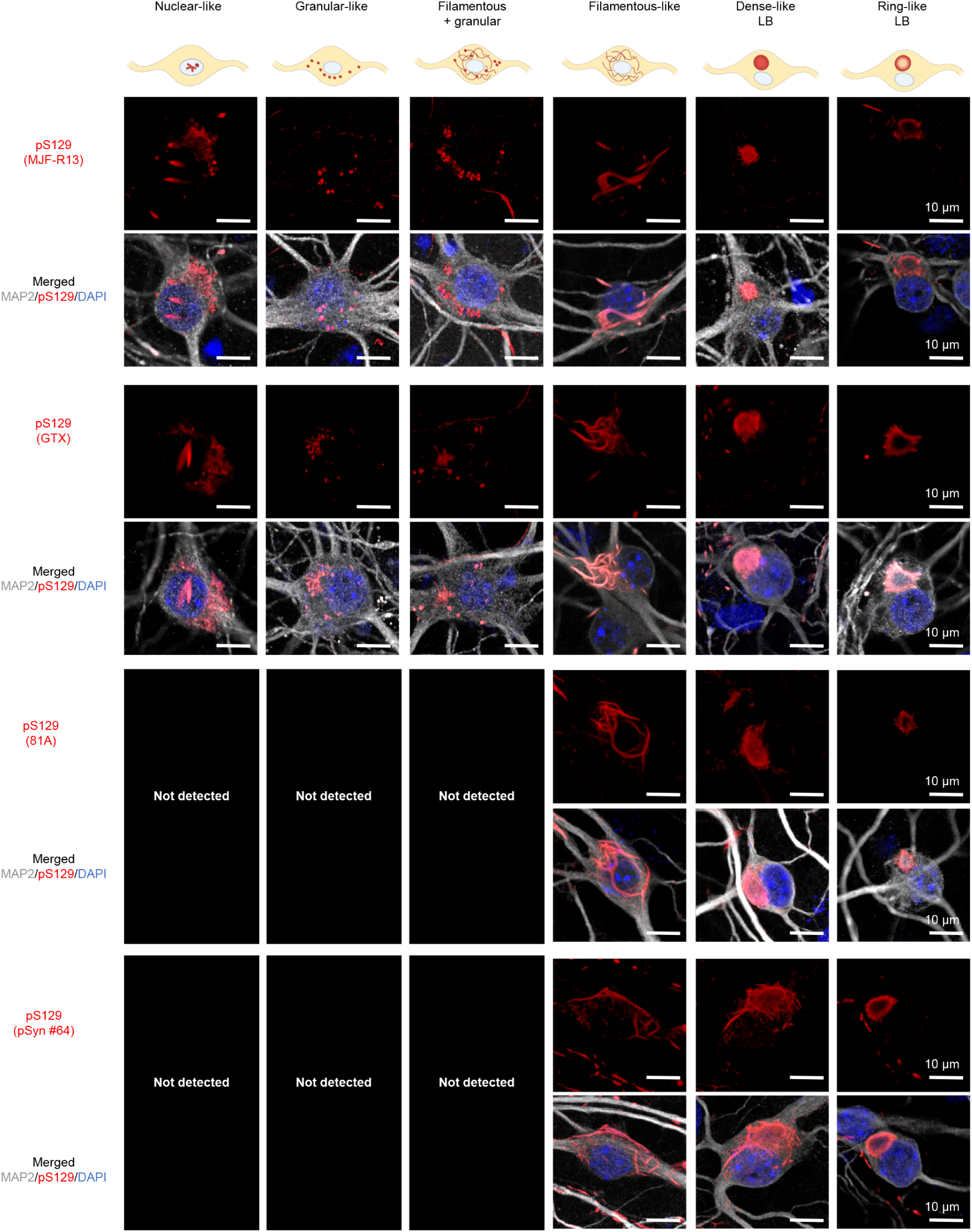
All the pS129 antibodies do not detect the morphological diversity of the pS129-aSyn seeded-aggregates formed in human PFFs-treated cortical neurons. After 21 days of treatment with 70 nM of human E83Q aSyn PFFs, primary cortical neurons were fixed, and ICC was performed. The seeded-aggregates were detected using the pS129 antibodies (MJF-R13, GTX, 81a or pSyn#64). Neurons were counterstained with the MAP2 antibody and the nucleus with DAPI staining. Scale bar = 10 μM. All four pS129 antibodies were able to distinguish the seeded-aggregates with the filamentous-like morphology and the LB-like inclusions with the dense core and the ring-like morphology. However, only the MJF-R13 and GTX antibodies were able to detect the nuclear, granular or granular and filamentous-like aggregates. The schematic representation of the different types of seeded-aggregates was created using BioRender.com.

We chose to perform this study in primary cortical neurons as the neuropathological characterization of the human brain of the E83Q mutation carrier showed the highest pathology in the cortex^37^. Confocal imaging shows that all the four pS129 antibodies were able to detect the newly seeded-aggregates with the filamentous-like morphology, the LB-like inclusions with the dense core resembling cortical LBs^15,38-43^ or LB-like inclusions with a ring-like morphology resembling brainstem/nigral LBs^15,17,18,38,44-46^, at D21 in both human E83Q and mouse WT PFF-treated cortical neurons (Figures 5 and S7). However, only the MJF-R13 and GTX antibodies could also detect the presence of the nuclear, granular, or granular and filamentous-like aggregates formed at D21 in the E83Q PFF-treated cortical neurons (Figure 5). These results are not consistent with a previous report by Grassi et al. ^47,48^, suggesting that the GTX antibody detects only dot-like/granular structures.

Taken together, these results demonstrate that not all commercially available pS129 antibodies can capture the aSyn pathological diversity in the neuronal seeding model to the same extent. This highlights the importance of using multiple antibodies to characterize cellular models of aSyn pathology and identify which antibody or combination of antibodies is the most appropriate for capturing the diversity of aggregates or specific aggregate morphology of interest.

### Specificity and cross-reactivity of the pS129 antibodies in primary neurons

Consistent with a previous study^10^, ICC combined with confocal imaging revealed that some of the pS129 antibodies showed unspecific immunoreactivity in the cytosol or the nucleus of primary neurons. This prompted us to assess further the specificity of the six antibodies for ICC and WB analyses in neurons lacking aSyn (KO neurons) and treated with PFFs. Under these conditions, the internalized PFFs undergo rapid cleavage at residue 114, thus eliminating the epitopes for all the pS129 antibodies. Several pS129 antibodies exhibited non-specific labelling in aSyn KO hippocampal and cortical primary neurons by ICC. For example, confocal imaging revealed that the pSyn#64 and MJF-R13 antibodies detect an unspecific and intense signal in the cytosol of the hippocampal (Figure S8A) and cortical (Figure S8B) primary neurons despite a very low laser intensity (∼1–2.5%) and using a photomultiplier (PMT) gain of around 600. On the other hand, the 81A and the LASH-EGT antibodies showed a similar background but only at much higher laser intensity (∼8%) (Figure S8). Under these setups, we also observed that the pSyn#64, MJF-R13, and the LASH-EGT antibodies had a strong reactivity in the nucleus of both the hippocampal and cortical KO (Figure S8) but also WT (Figure 4A) neurons. These findings are consistent with previous and recent reports demonstrating that MJF-R13, pSyn#64 and 81A antibodies show extensive staining and cross-reactivity in aSyn KO mice neuronal cultures^10^.

Conversely, no non-specific and/or background signal was observed in primary neurons stained with the GTX or the EP1536Y antibody, even with ∼8% laser intensity (Figure S8). Using neurons containing pS129 positive aggregates (Figure 4A), we found that adjusting the setup of the laser intensity and the PMT gain and offset helped eliminate the unspecific signal initially observed in the aSyn KO primary neurons (Figure S8C and D). Altogether, our findings underscore the critical importance of using the appropriate positive (recombinant pS129 or neurons containing pS129 positive aggregates) and negative (aSyn KO cells/tissues) controls to allow for identifying off-target proteins and signals recognized by pS129 antibodies, thus enabling more accurate interpretation of experimental results based on imaging.

### All pS129 antibodies cross-react with several nuclear proteins that could be mistaken for full-length, truncated, and oligomeric forms aSyn

Initial studies reported increased accumulation of pS129 with age in the cortical brain areas and dopaminergic neurons of aSyn transgenic mice^49,50^. In these studies, pre-treatment with phosphatases resulted in the disappearance of the pS129 immunoreactivity. Wakamatsu et al. also reported that nuclear aSyn is preferentially phosphorylated compared to cytosolic aSyn^50^. However, most of these studies were based on immunohisto-/cytochemistry and relied on the use of a single pS129 antibody^50-54^ (Figure S1). Furthermore, the cross-reactivity of pS129 antibodies towards nuclear proteins has not been systematically assessed. We previously reported that some PD-linked mutations, such as E46K, enhance aSyn nuclear localization and detection of pS129 by ICC but failed to detect an increase in pS129-aSyn by WB^9^. We speculated that this could be due to the presence of predominantly truncated and phosphorylated C-terminal fragments of aSyn in the nucleus. A recent study by Koss et al. reported strong pS129 staining in neuronal cells in the brain of healthy and DLB individuals^55^. Despite the strong signals detected by ICC, WB analysis showed that the levels of pS129 are minimal in the nuclear fraction (< 5%) relative to the total aSyn protein level^55^. The discrepancy between the ICC signal and levels of pS129 detected by WB suggests that the pS129 antibodies could potentially be cross-reacting with other nuclear proteins. Interestingly, the authors reported specific bands above that of the aSyn monomer, which is seen in the nuclear fraction of the DLB cases but not in the nuclear or cytoplasmic fractions from healthy controls. However, in the absence of the proper controls to establish the purity of the nuclear fractions and confirmation of pS129 detection by mass spectrometry, this data remains inconclusive. To help address such discrepancies and identify better tools for assessing nuclear pS129-aSyn, we evaluated the specificity of the six pS129 antibodies towards nuclear (N), cytosolic (C), and membrane-associated proteins (M) but also the soluble and insoluble fractions by performing biochemical subcellular fractionation of lysates from neurons (Figure 6) and mammalian cells (Figures S9 and S11).

First, we assessed by WB the pS129 levels in the nuclear, cytosolic, and membrane fractions (Figure 6A) and the soluble/insoluble fraction (Figure 6B) of the primary hippocampal neurons from WT and aSyn KO mice. We observed a strong cross-reactivity with several proteins of various sizes, particularly in the nuclear and membrane fractions (Figure 6A), where all pS129 antibodies show prominent detection of a 16–17 kDa band that exhibits similar SDS-PAGE mobility similar to that of full-length recombinant aSyn, and truncated aSyn species usually detected in neuronal aSyn seeding models. These bands were observed in the nuclear fractions from WT and KO neurons, thus establishing that they do not represent aSyn species. Interestingly these bands were absent in the cytosolic fractions. However, all pS129 antibodies also showed cross-reactivity to several proteins with molecular weight ranging from 25 to ∼200 kDa (Figure 6). Although the aSyn sequence does not contain nuclear localization signals, several studies have reported nuclear localization of neuronal aSyn in human and rodent brains^49-51^. However, the role of nuclear aSyn in health and disease remains a subject of intense debate. aSyn interactions with DNA and histones, regulation of gene expression^53^, and induction of transcriptional dysregulation^52,56^ of cell cycle-related genes have been proposed to be fundamental mechanisms contributing to aSyn-induced toxicity in PD and other synucleinopathies^57^. However, which forms of aSyn exist in the nucleus and the factors involved in regulating the aSyn life cycle in the nucleus or shuttling between the nucleus and cytoplasm remain unknown.

As mammalian cell lines are frequently used to study the cell biology of aSyn, investigate its aggregation mechanisms, and screen for modifiers of aSyn aggregation, we also evaluated the specificity of the pS129 antibodies towards soluble and insoluble proteins from HEK and HeLa cells transfected with WT aSyn plasmid or empty vectors. All six antibodies showed non-specific detection of several bands in both cell lines, with the 17 KDa species being the most abundant, mainly in the insoluble fractions (Figure S9). There was no difference in the bands detected in cells transfected with aSyn plasmids or an empty vector. Although the intensity of the 17 kDa band increased in cells overexpressing aSyn, monitoring aSyn levels using a pan-aSyn antibody suggests that it is unlikely that this band corresponds to phosphorylated aSyn proteins. Interestingly, more non-specific bands were detected in the soluble and insoluble fractions from HEK cells. Furthermore, EP1536Y detected more non-specific bands in the soluble fractions than the other five antibodies.

Interestingly, the same prominent nuclear bands pattern observed in primary neurons was also detected in the nuclear fractions from mammalian cells (HeLa and HEK293) transfected with WT aSyn or an empty vector (Figure S10). Notably, the pattern of proteins that cross-react with pS129 in the membrane fractions was distinct from that seen in the nuclear fractions. Furthermore, the non-specific bands seen in the nuclear fraction exhibit similar but not identical mobility to that of newly formed fibrils seen in many of the aSyn seeding-based neuronal models, including the appearance of two prominent bands with MW of ∼16–17 and ∼13–12 KDa, similar to that of full-length and truncated aSyn species, in addition to a band at ∼50 kDa and other HMW species. The small differences in size in relation to WT aSyn can be discerned more easily using Tricine gels (Figure S11). Taken together, these findings demonstrate that all pS129 antibodies show strong cross-reactivity with membrane and nuclear proteins. The pattern of pS129 non-specific immunoreactivity detected by WB could be mistaken for pS129 species seen in the aSyn neuronal seeding models. These findings underscore the importance of not only validating the pS129 antibodies but also using the appropriate 1) control samples (e.g., recombinant WT and pS129 and lysates from aSyn KO neurons or cells that do not express aSyn); and 2) SDS-PAGE gels that allow clear separation of proteins between 10-20 KDa (e.g., Tricine gels).

### Some pS129 antibodies display unspecific signals in aSyn KO mouse tissue

To further assess the specificity of the pS129 antibodies in brain tissues, we stained tissue from different brain regions, including the amygdala, striatum, cortex, hippocampus, and substantia nigra, from aSyn KO animals (Figure 7A). The EP1536Y, GTX, and LASH-EGT antibodies did not show any background staining in all five regions. The MJF-R13 and 81A antibodies showed high background in all five regions, especially in the hippocampus, substantia nigra (81A), and amygdala and striatum (MJF-R13). The MJF-R13 staining showed mostly dotted structures around the nuclei or staining the nucleus itself, whereas the 81A showed mostly neuritic staining, especially in the hippocampus and substantia nigra. The pSyn#64 antibody showed an intermediate background with the highest background signal seen in the striatum and hippocampus. The high background signal of the 81A and pSyn#64 antibodies could be because these two antibodies are raised in the same species (mouse) of the target tissue (mouse). On the other hand, this does not explain the strong neuritic staining present in the hippocampus of 81A-stained tissue (Figure 7A). In our hands, the EGT-LASH and GTX antibodies are highly specific to pS129 and do not recognize other proteins when used for IHC application. Our results are in line with a previous report by Delic et al., where MJF-R13 exhibiting the highest background revealed as punctate intracellular structures and the 81A antibody displaying off-target signal reminiscent of neuronal processes^4^.

**Figure 7.**
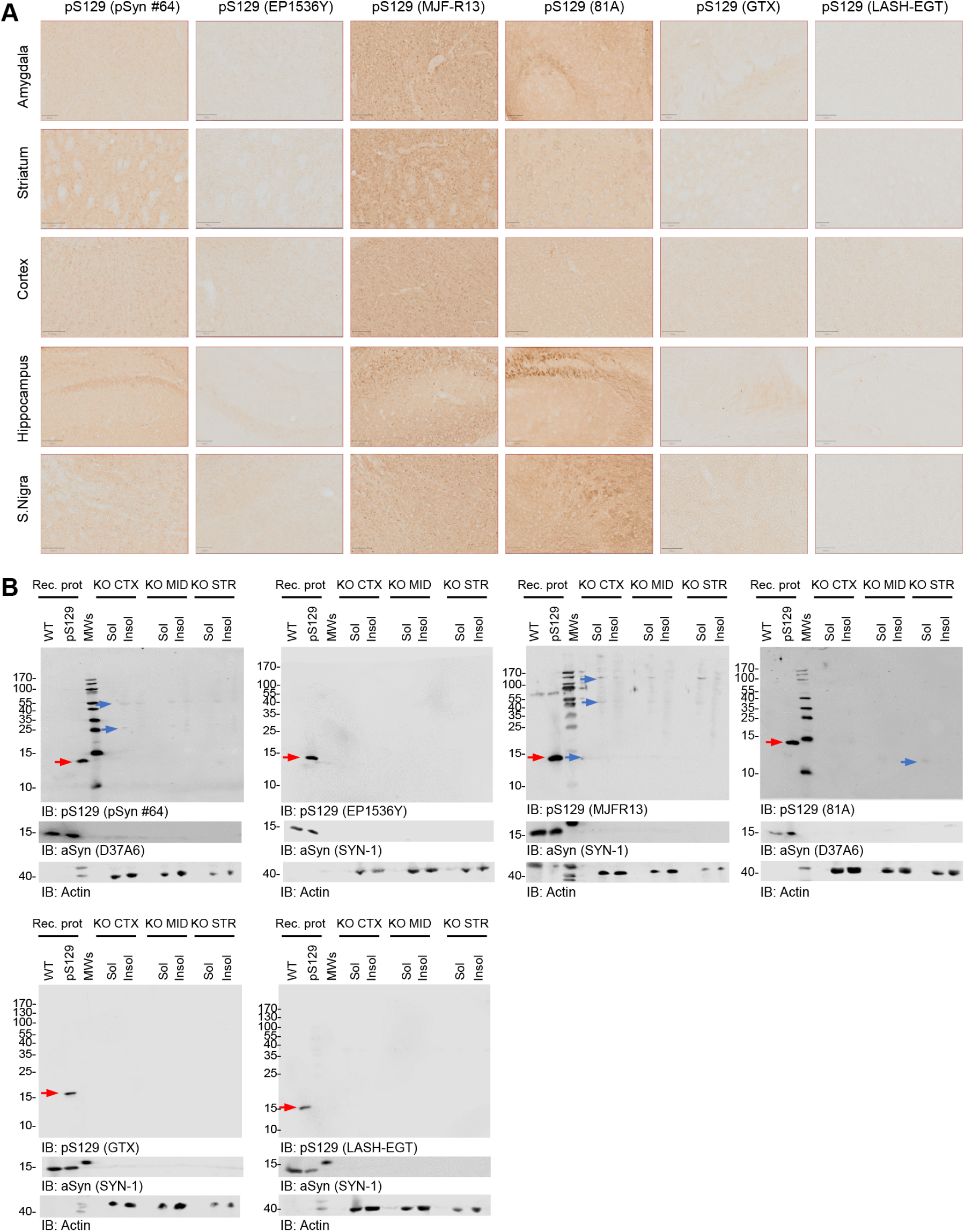
Assessment of pS129 antibodies detection in brain sections of aSyn KO mice. **A**. Brain coronal sections (50 μM thick) from different brain areas of aSyn KO mice (Amygdala, Striatum, Motor Cortex, Hippocampus and Substantia Nigra) have been examined to assess the different staining patterns of the six pS129 antibodies (pSyn#64, MJF-R13, 81a, EP1536Y, GTX and LASH-EGT). Scale bar = 100 μM. **B**. pS129 level of detection in the soluble and insoluble fractions extracted from the aSyn KO mice brains using the six pS129 antibodies (pSyn#64, MJF-R13, 81a, EP1536Y, GTX and LASH-EGT). Sequential biochemical extraction was performed on different brain regions (cortical/CTX; Midbrain/MID and Striatum/STR) derived from aSyn KO mice, and the level of pS129 was assessed in the soluble (0.1% Triton-soluble fraction) and the insoluble fraction (SDS-soluble fraction). 40 ng of recombinant aSyn WT or phosphorylated at residue S129 (pS129, indicated by the red arrow) were used as positive controls. Membranes were counterstained for total aSyn (SYN-1 or D37A6), and actin was used as a loading control. The red arrows indicate the pS129-aSyn positive bands. The blue arrows indicate the non-specific bands.

Consistent with our findings, Delic et al. reported that EP1536Y shows the highest specificity with little to no off-target labelling and the same order of background signal as shown in Figure 7A^4^. In a recent study, Arlinghaus et al. showed that three pS129 antibodies (MJF-R13, pSyn#64, and EP1536Y) showed cross-reactivity in brain slices or primary cultures (by IHC or ICC)^10^. Similar to what we report, Arlinghaus and colleagues show that MJF-R13 and pSyn#64 stained cell bodies throughout all regions analyzed, with no differences between WT and *Snca* KO mouse tissues^10^. Furthermore, 81A did not show cell body staining but displayed higher background in the cortex and substantia nigra than other areas. We also highlighted that 81A did not show somatic staining but rather a neuritic off-target signal. Nevertheless, we detected higher background in the substantia nigra, amygdala, and especially the hippocampus, with little to no background in the cortex. In contrast to our findings, these authors also report that EP1536Y displays somatic staining, whereas we detected little to no unspecific signal.

To better understand the sources of the background signals, we assessed the specificity of the antibodies in the cortex, midbrain, and striatal lysates from aSyn KO animals (Figure 7B). The three antibodies (EP1536Y, GTX, and LASH-EGT) that did not show a background signal by IHC also did not show any bands by WB. Even at high exposure, we only observed one HMW band at ∼120 kDa for the LASH-EGT antibody (Figure S12). In contrast, the MJF-R13 showed several bands ranging from 25 to 170 kDa in the soluble and insoluble fractions of lysates from the three brain regions, consistent with the high background signal observed by IHC (Figure 7A). The pSyn#64 antibody, at high exposure, showed a strong signal for a band with a molecular weight that runs slightly higher than aSyn and HMW bands. The 81A shows a single band with a molecular weight below 15 kDa in the soluble fractions from the midbrain and striatal tissue lysates (Figure 7B). These results are in line with those described by Delic and colleagues, which also revealed the highest off-target signal for MJF-R13^4^.

Nevertheless, in our hands, the EP1536Y antibody did not present any non-specific high molecular weight bands, probably due to differences in the composition of the extraction buffers used in the two studies. Similarly, Delic et al. report that the 81A antibody shows a single 75 kDa band when using SDS extraction buffer^4^, whereas in our case, a lower band appears at higher exposure in the Triton-soluble fraction, but not in the insoluble fraction (Figure S12). Overall, these findings are consistent between WB and IHC and demonstrate that some of the commonly used antibodies (e.g., 81A and MJF-R13) are not specific to aSyn and recognize other proteins in the brain. Among all the antibodies tested, EP1536Y and GTX appear to be the most specific for pS129.

### pS129 antibodies display different staining patterns in aSyn mPFFs-injected WT mice based on the brain region analyzed

To determine if the pS129 antibodies exhibit differential immunoreactivity towards different aSyn pathological aggregates, we stained brain slices from WT mPFFs injected mice 3 months after injection (Figure 8). In the motor cortex, all six antibodies were able to capture, to a different extent, neuritic and somatic aSyn aggregates (Figure 8). Nevertheless, EP1536Y showed the strongest staining of aSyn aggregates, with similar immunoreactivity towards neuritic or somatic aggregates. The appearance of these aggregates varies from skein-like structures to long serpentine neuritic and dense perinuclear aggregates. In contrast, MJF-R13 and pSyn#64 showed weaker staining, with a slight preference of MJF-R13 towards perinuclear aggregates and some partial fibrillary neuritic staining, whereas 81A greatly privileged this subtype (Figure 8).

**Figure 8.**
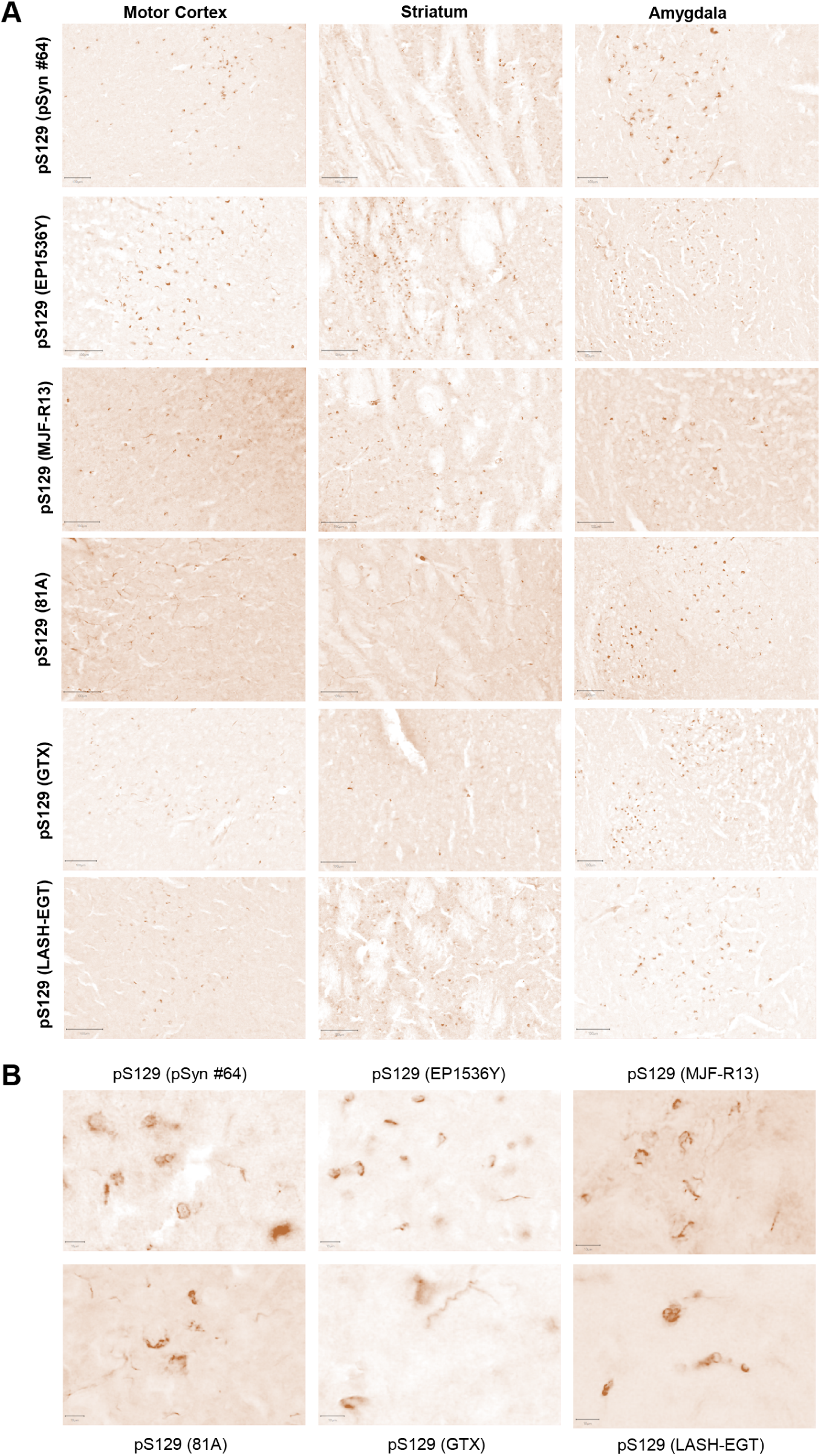
Evaluation of staining pattern of aSyn pathological aggregates in WT PFF-injected mice. **A** Brain coronal sections (50 μM-thick) from different brain areas of C57BL6/J PFF-injected mice (Amygdala, Striatum, Motor Cortex), 3 months post-injection, have been examined to assess the different staining patterns of the six pS129 antibodies (pSyn#64, MJF-R13, 81A, EP1536Y, GTX and LASH-EGT). Scale bar = 100 μM. **B**. Microphotographs of the amygdala stained with the six pS129 antibodies (pSyn#64, MJF-R13, 81A, EP1536Y, GTX and LASH-EGT) at higher magnification. Scale bar = 10 μM

In the striatum, pSyn#64 consistently showed a preference for detecting dense perinuclear aggregates and skein-like aggregates, although once again underestimating a load of aSyn pathology, compared to EP1536Y. MJF-R13 and LASH-EGT, in this brain region, highlighted mild aSyn pathology without any preference for neuritic vs somatic pathology and revealed diffuse cytoplasmic aggregates, as well as skein-like and neuritic tubular structures. On the other hand, GTX and pSyn#64 underperformed in terms of pathological load evaluation, compared to EP1536Y, which, once again, proved to be the antibody that most reliably detects pathological aggregates by also detecting very dense round cytoplasmic aggregates. Finally, in the amygdala, all antibodies were able to equally reveal the high level of pathological load, except for MJF-R13, which was only partially stained for pS129-positive inclusions. Interestingly, in this region, the pSyn#64 antibody displayed better the diversity of the aggregates (Figure 8). It is also interesting to note that EP1536Y displayed a preference towards dense intracytoplasmic inclusions over aggregates in neuronal processes, which appeared more prominent in the other antibodies. It is important to note that the GTX antibody showed highly variable staining patterns from animal to animal. Consistent with a previous study by Delic and colleagues, EP1536Y showed the highest signal specificity and captured better the different forms of aSyn aggregates. Nevertheless, in our hands, MJF-R13 did not reveal as much pathology load as reported by Delic and colleagues in the striatum. Under our experimental conditions, all the antibodies displayed a similar capacity to stain aSyn aggregates with a similar propensity towards long neuritic aggregates (Figure 8). Overall, EP1536Y proved to be the most reliable antibody to assess aSyn pathology in all brain regions in mouse tissue. The differential sensitivity and specificity of the antibodies in different brain regions could partially explain the experimental variability observed across different laboratories. Therefore, we recommend that the performance of new antibodies should be systematically assessed in different brain regions before they are used to assess and quantify aSyn pathology.

### Identification of putative candidate proteins that cross-react with pS129

Although several studies have shown that pS129 cross-reacts with selected proteins, a systematic analysis and cataloguing of all possible proteins that could cross-react with pS129 antibodies has not been performed. Therefore, we screened all known human and mouse phosphorylation sites annotated in Uniprot (https://www.uniprot.org) for sequence similarities to peptide sequences comprising not only S129 but all the other phosphorylation sites that have been detected in LBs and PD brains (Y39, S87, and Y125). In addition, we included peptide sequences comprising Y133 and Y136. Entries for humans (n = 7662) and mice (n = 7133) were retrieved as full text. A Perl script was then used to parse the entries and extract protein sequence fragments of 13 residues centred on the annotated phosphorylation sites to compare them to the six human aSyn fragments. Figure 9A shows the hits of only those whose residues preceding and following the phosphorylation site were perfect matches. For all protein fragments retrieved from Uniprot satisfying this condition (Figure S13), we computed the number of exact and highly similar matches to the six aSyn fragments shown in Figure 9A and the length of the continuous stretch of perfect or similar matches encompassing the phosphorylation site (see material and method section for details).

**Figure 9.**
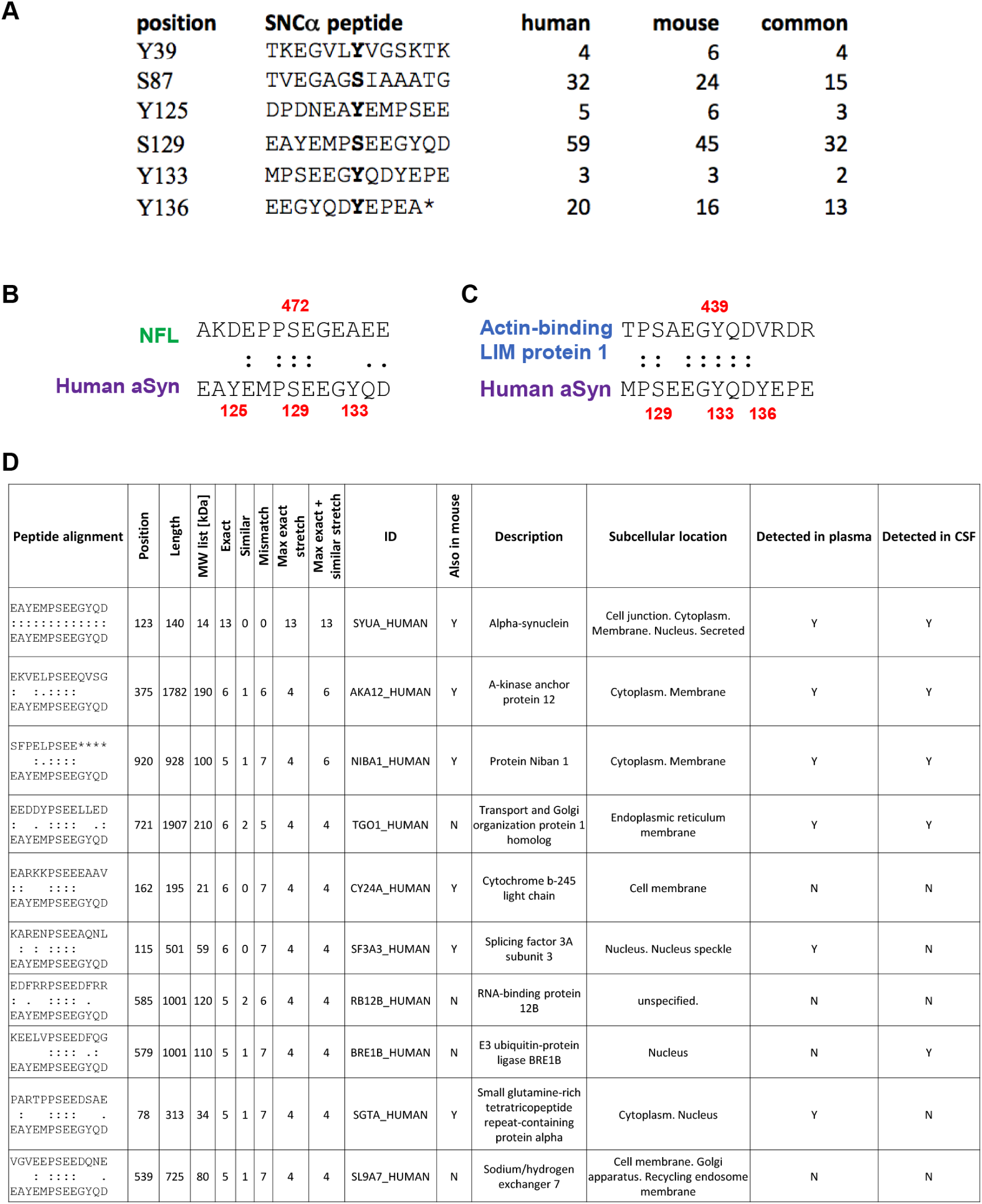
Identification of putative candidate proteins that could cross-react with aSyn antibodies. **A**. List of peptides used to search the Uniprot database to identify putative proteins that could cross-react against antibodies raised against the known aSyn phosphorylated residues highlighted in bold at the middle of each sequence. The number of candidate hits retrieved for human, mouse, or common to both organisms is indicated for each peptide. The * present in the last sequence indicates the C-terminal residue. **B**. Sequence alignment of the region around human aSyn pS129 and Neurofilament light polypeptide (NFL) pS472, which is known to cross-react with antibodies raised against pS129-aSyn. **C**. Sequence alignment of the region around human aSyn pY133 and Actin-binding LIM protein 1 pY439, illustrating the high level of similarity. **D**. List of the top ten candidate proteins that can cross-react with pS129 antibody.

The major number of hits was returned with the search using the S129 anchor peptide, with 59 and 45 human and mouse hits, respectively, of which 32 are in common (Figures 9A and S13). This is, for example, the case for the NFL, which has been reported to cross-react with antibodies raised against fragments of aSyn pS129^8^. Interestingly, NFL was ranked 44 (out of 59) and 35 (out of 45) for human and mouse, respectively (Figure S13A-B). Furthermore, it bears a modest local similarity with the aSyn fragment, where both protein fragments share a continuous stretch of three conserved residues, a fourth disjoint but conserved residue and two further residues with high similarity scores (Figure 9B). This indicates that several other hits have sequences that are locally even more similar to the aSyn S129 fragment than NFL, several bearing continuous stretches of four fully conserved residues as well as additional similar residues and may consequently also cross-react with pS129 antibodies, depending on their local structure and experimental conditions. Based on the local conservation metric, the best candidate for cross-reaction is the Actin-binding LIM protein 1, which shares a local stretch of five conserved residues with the phosphorylated Y133 aSyn region along with two additional fully conserved residues nearby Figure 9C.

Of note, the list of candidate proteins that could cross-react with antibodies raised against phosphorylated aSyn is likely not exhaustive because the hits reported are limited to phosphorylation sites annotated in Uniprot at the time of the search (Figure 9D and tables in Figure S13). For example, among the six aSyn seed fragments used to scan the database, only three identify aSyn as a candidate protein (S87, Y125, and S129). These three residues are currently annotated as phosphorylated in Uniprot https://www.uniprot.org. Only Y125 and S129 return aSyn as a hit for mice because an asparagine that cannot be phosphorylated is present at position 87 instead of a serine. The fact that a hit is found both in humans and in mice does not *per se* reflect a higher potential for cross-reactivity. Instead, this merely reflects that a phosphorylation site is not necessarily conserved between human and mouse proteins, as mentioned above for aSyn at position 87, or that the two residues surrounding the phosphorylation site are not strictly conserved between human and mouse, a condition imposed during our scanning strategy. Nevertheless, taken collectively, these hits represent a list of candidate proteins that should be considered for putative cross-reaction with antibodies raised against phosphorylated aSyn fragments.

As a next step, we investigated how many of these candidate proteins that could cross-react with pS129 antibodies were previously identified in biological specimens. From the total number of candidate proteins (n = 121) that could potentially cross-react with the different aSyn PTMs (i.e., pS129, pS87, pY39, pY125, pY1233, and pY136) (Figure S13), 47% (n = 57) and 31% (n = 38) were previously reported to be detected in human plasma and cerebrospinal fluid (CSF), respectively. Particularly, when focusing on the proteins (n = 59) that could react with pS129, the percentage of proteins previously identified in plasma was 45.8% (n = 27) and 32.2% (n = 19) in CSF. Several other proteins were also identified to potentially cross-react with the different aSyn PTMs, in plasma and CSF (Figure S13A).

Whether cross-reactivity with some of these candidate proteins could be contributing to the high variability of pS129 across the different biomarker-based studies remains unknown. This is in part because the levels of the phosphorylated forms of these proteins in biological fluids are unknown. However, their presence and a high potential for cross-reactivity with phosphorylated aSyn antibodies underscore the importance of validating the specificity of pS129 antibodies in biological fluids, which is rarely done. This could be done using CSF or plasma sample standards where the desired phosphorylated aSyn species have been depleted.

Although cross-reaction should be tested experimentally, we hope that the list of proteins we provide here (see tables in Figure S13B and S14) could help to better assess the specificity of future antibodies against disease-associated phosphorylation sites. The local alignments can also be scrutinized to check in more detail the similarity based on other criteria not considered during the ranking, such as the local charge.

## Discussion

Among all the research tools and reagents used by scientists daily, antibodies have the greatest influence on shaping our knowledge and current hypotheses about the pathophysiology of PD and neurodegenerative diseases. Therefore, it seems obvious that the use of poorly characterized antibodies or antibodies that show non-specific cross-reactivity could lead to misinterpretation of experimental observation and misdirection of time, investments, and resources^58,59^. The work presented here, combined with previous studies^4,5,8,10,21^, demonstrates that the development of antibodies that target specific post-translationally modified or aggregated forms (oligomers, fibrils, LNs or LBs) aSyn remains challenging. The level of cross-reactivity we detected also underscores the urgent need for better antibodies targeting pS129-aSyn and other modified forms of aSyn. In this section, we reflect on the implications of our work for investigating the mechanisms underpinning aSyn pathology formation and their role in the pathogenesis of PD. We also discuss how we can apply the lessons learned from previous efforts to develop PTM and conformation-specific aSyn antibodies to develop more robust pipelines for the development, characterization and validation of such antibodies.

Today, the histopathological staging of LB diseases^60-62^ and the assessment of pathology formation and spreading capacity in aSyn-based animal^63-66^ and cellular^67-69^ models of PD and related diseases rely mainly on antibodies targeting the C-terminal domain^60-62 70,71^ of the protein and more recently pS129 immunoreactivity^2,72-74^. Very often, such studies rely on a single aSyn antibody and, more recently, primarily pS129 antibodies^74,75^ (Figure S1).

It is now well established that aSyn in LBs is not only subjected to multiple PTMs, including phosphorylation, ubiquitination, truncation, and oxidative nitration^76,77^, but also exists as a mixture of different species with different PTM patterns and localization within LBs. Given that the majority (>90%) of aSyn in LBs is phosphorylated on S129 residue, it is plausible to assume that the detection of other PTMs within LBs suggests that aggregated forms of aSyn bearing these PTMs are also phosphorylated at S129^1^. Our results demonstrate that the presence of other PTMs in close proximity to pS129 dramatically influences antibody-based detection of monomeric^78^ and aggregated forms of pS129-aSyn in a site, PTM-type, and PTM-number dependent manner. These findings combined with increasing evidence demonstrating that aSyn aggregates bear multiple modifications and are enriched in C-terminally truncated species, suggest that it is unlikely that antibodies against pS129 or any single C-terminal (a.a. 115-140) targeting antibody are capable of capturing the diversity of aSyn species or aSyn pathology in the brain. It is noteworthy that several studies have reported that aSyn astroglial pathology is not detectable using C-terminal targeting aSyn antibodies but can be revealed with antibodies targeting the NAC domain of the protein^62,79-82^. In a recent study, Henderson et al. reported that mice treated with the Glucocerebrosidase inhibitor conduritol-β-epoxide CBE develop spheroid structures that are recognized by the 81A but not the EP1536Y pS129 antibody^83^. Although the molecular basis of this differential labelling by pS129 antibodies remains unclear, these observations underscore the importance of using multiple pS129 and aSyn antibodies to characterize aSyn species and pathology^4,8^.

### Specificity and sensitivity of the pS129 antibodies in peripheral tissues

Several studies have focused on using this pS129-aSyn as a diagnostic biomarker of aSyn pathology in peripheral tissue and biological fluids. pS129-aSyn has been detected in different peripheral tissues, such as colon^84^, skin^85^, and in plasma^86,87^. The enteric system (ENS) has been proposed to play a pivotal role in the initiation and propagation of PD pathology and the gut-brain axis in PD and other synucleinopathies. However, the lack of antibodies that robustly and reliably detect pS129-aSyn in the ENS has hampered efforts to elucidate the role of aSyn phosphorylation in regulating aSyn biology and pathology formation in the ENS. While several studies have reported on the detection of pS129 by immunofluorescence, biochemical evidence for the accumulation of pS129 in the ENS and peripheral tissue remains sparse and lacks validation. However, very little is known about the aggregation state of pS129-aSyn in peripheral tissues or biological fluids. One study by Preterre et al. reported that several pS129 antibodies, including two of the six used in this study (EPI1536Y and pSyn#64), failed to robustly detect pS129 by WB and lacked the specificity to detect pS129 in the gut^88^.

Lassozé et al. evaluated the specificity and sensitivity of five commercially available pS129 antibodies (sc-135638, Invitrogen: #482700, ab59264, EP1536Y, and D1R1R) in primary cultures of ENS^25^. They reported that one of the five antibodies, D1R1R, was the most specific (no detection of non-specific bands by WB) and reliable in detecting endogenous pS129 in primary rat cultures of ENS. Unfortunately, this study displays many shortcomings that make it difficult to draw any conclusions about their findings. First, none of the experiments included a proper positive control for pS129. For example, the antibodies were assessed for their ability to detect non-modified aSyn, but a positive control of recombinant site-specific pS129 was not included, although such standards are commercially available (MJFF and Proteos, Inc). Furthermore, no brain sample known to contain detectable levels of pS129-aSyn was included as controls in the WB analysis of the primary culture of ENS. Second, the specificity of the antibodies, including D1R1R, was not evaluated using primary cultures of ENS from *SNCA* KO mice. Third, a direct comparison of the performance of the five antibodies by immunofluorescence, under the condition of based and KCI-induced increase in pS129 levels, was not carried out or presented. In this regard, only the immunofluorescence data for D1R1R was presented. One surprising finding reported in this paper is their observation that three of the pS129 antibodies they used also recognize non-modified monomeric aSyn. This contrasts with our data (Figure 2) and others ^5^, where we clearly show using modified and unmodified proteins that EP1536Y recognize specifically pS129 but not the unmodified protein using standard protein concentrations of 40–100 ng. These observations and limitations underscore the importance of conducting proper characterization and validation of the antibodies before drawing conclusions or recommendations that could lead to misdirecting efforts and resources and add more confusion to the literature.

### Relevance of the pS129 antibodies in evaluating pS129-aSyn physiological function

Increasing evidence suggests that phosphorylation of aSyn regulates the structural properties^89^, interactome, and subcellular localization of many aspects of aSyn physiological functions^90^. Therefore, there is an urgent need to develop antibodies that are capable of detecting both physiological and pathogenic pS129 species. Recently, Arlinghaus et al. described a new Proximity Ligation Assay (PLA) that allows for the specific detection of endogenous soluble pS129-aSyn in primary neurons and in mouse and human brain sections with significantly reduced background signal and cross-reactivity^10^. Their results suggest that pS129 is ubiquitously expressed throughout the mouse brain and is elevated in the cell bodies within specific brain regions, suggesting more specific roles of pS129 in these brain regions. Despite the failure of the PLA assay to detect aggregated pS129 in human brain tissues, these results are encouraging and suggest that the use of a combination of pS129 and other aSyn antibodies allows for the development of more rapid, compared to PLA, and specific assays and methods for detecting endogenous pS129-aSyn. Furthermore, Killinger et al. showed a high abundance of pS129 in the mitral cell layer of the olfactory bulb^90^. Altogether, these findings suggest a potential physiological role for S129 phosphorylation in aSyn biology in health and disease and underscore the importance of developing more specific and sensitive antibodies against aSyn phosphorylated at S129 or bearing other modifications.

Increasing evidence shows that physiological and aggregated forms of aSyn exist in different conformational states and exhibit distinct interactomes (Figure 10). Although most existing structural data suggest that the regions harboring the majority of aSyn PTMs (extreme N- and C-terminal domains) remain flexible and accessible, this by no means guarantees that antibodies targeting PTMs in these regions would be capable of capturing this diversity of aSyn species. The presence of additional PTMs near the PTM of interest or interaction of aSyn species with other proteins could mask or interfere with the detection of specific aSyn species. Furthermore, aSyn within LBs exists in a complex environment where aSyn aggregates not only modified at multiple residues but interact with membranous organelles and a large number of proteins (Figure 10 A), mainly through the remaining flexible domains flanking the amyloid core. Furthermore, recent studies on aSyn fibrils obtained by amplifying brain-derived fibrils demonstrate that the C-terminal domain sequence comprising the residues S129 is highly structured and included in the core sequence of the fibrils (Figure 10 A). These observations suggest again that relying on only C-terminal targeting antibodies to detect, monitor, or quantify aSyn pathology may lead to underestimation or even misrepresentation of aSyn pathology.

**Figure 10.**
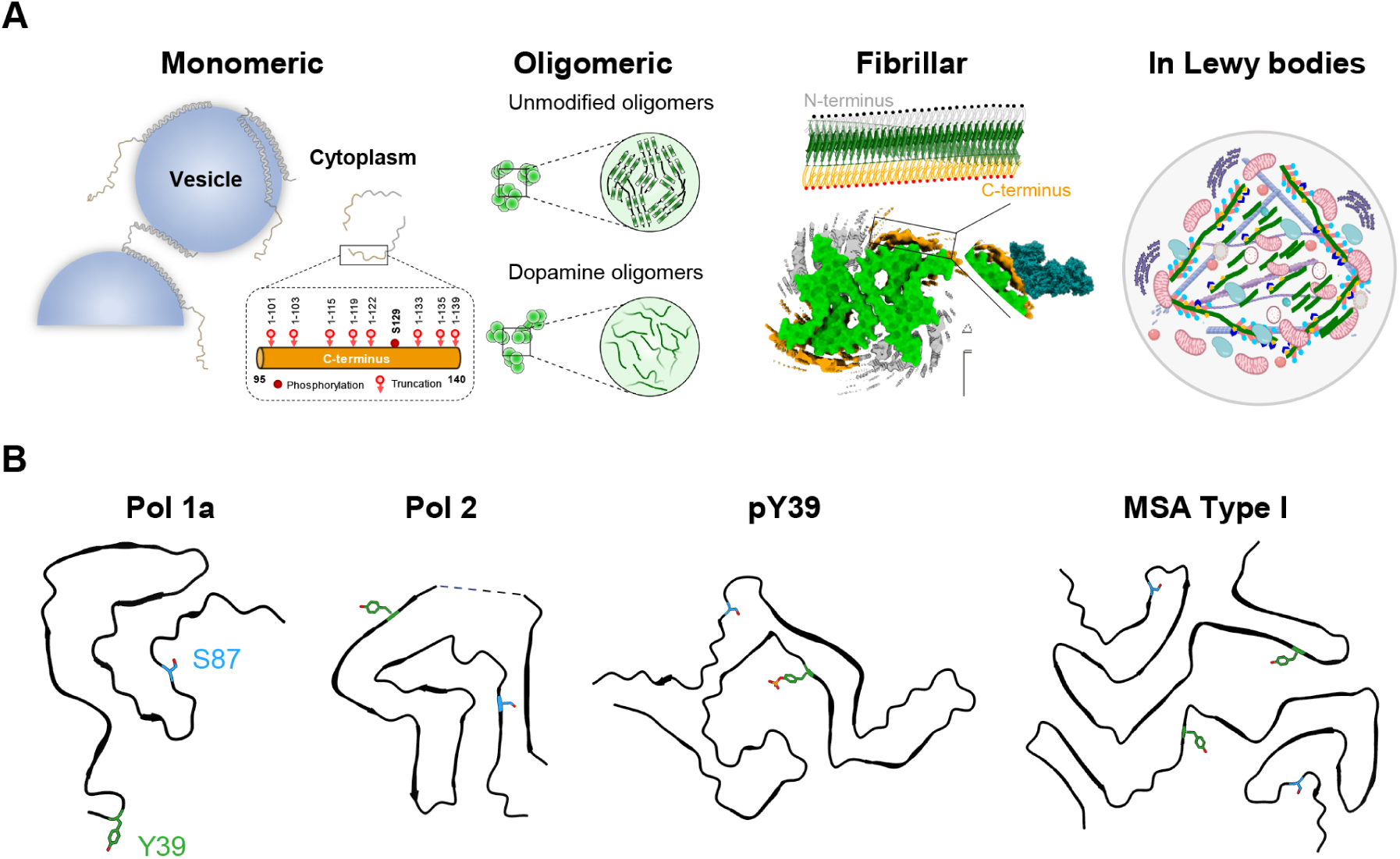
The multiple faces of aSyn and their susceptibility to post-translational modifications. **A**. The C-terminal domain of aSyn remains disordered and accessible in most aSyn species, from monomers to oligomers, fibrils and Lewy bodies; Monomeric forms of aSyn are predominantly disordered but adopt a-helical rich conformations when bound to vesicles. Despite the absence of an atomic model for aSyn oligomers, several types of aSyn oligomers of distinct conformations (disordered or b-sheet rich) and morphological properties have been observed during aSyn in vitro aggregation^91^. Near-atomic models for aSyn fibril core structures solved using cryo-EM (depicted model PDB:6CU7, EMDB: EMD-7618)^100^. On top, a schematic depiction of the fibrils showing the amyloid core sequence (green), disordered N-terminus (grey), and disordered C-terminus (orange). In all cases, the protein’s C-terminus, which harbors various sites for the protein’s truncation and/or phosphorylation, is disordered and accessible. aSyn fibrils found in pathological aggregates are ubiquitinated (black dots), mostly in the N-terminal domain, and phosphorylated, especially at S129 (red dots), and nitrated at multiple tyrosine residues (Y125/Y133/Y136). In LB, modified aSyn fibrils represent one of the key components of Lewy bodies and interact with a large number of proteins, membranous structure organelles, and lipids. Many of these interactions are mediated by N- and C-terminal domains that harbor several sites of PTMs. Recent studies demonstrated that these PTMs play key roles in regulating aSyn fibrils remodeling and LB formation and maturation^31,34^. **B**. A selection of aSyn fibrils’ cryo-EM core structures to illustrate the diversity of aSyn fibril structures and the positions of other disease-relevant PTM sites (Y39 in green and S87 in blue) present in the fibril core. One protofilament of the *in vitro* derived fibrils is displayed, including Polymorph 1a (Pol 1a, PDB-6CU7)^100^, Polymorph 2 (Pol 2, PDB-6SST)^101^ and fibrils from pY39 monomers (pY39, PDB-6L1T)^102^. An example of the fibrils derived from the postmortem brain of multiple system atrophy patients is also displayed (MSA Type I, PDB-6XYO)^103^. The LB representation in **A** was created using BioRender.com. The fibril core icon “Amyloid fibril” was adapted from BioRender.com, 2022.

### Systematic characterization and validation of pS129 antibodies

Based on our findings and previous results, we propose a pipeline for systematic characterization and validation of pS129 antibodies before their application in PD research or their use in biomarker discovery and validation (Figures 1A). We propose that all current and future antibodies against pS129 or other aSyn PTMs be screened first using aSyn protein standards containing the target PTM and proteins containing other PTMs that are known to co-occur with the target PTM. The antibodies that are insensitive to neighbouring PTMs should then be screened using brain tissues and brain lysates from aSyn KO mice or rats. Antibodies that show specificity for the aSyn species/modification of interest should then be tested and optimized (e.g., dilutions) for the desired applications (e.g., studies on the brain and peripheral tissues or studies on animal and cellular models of synucleinopathies). In these studies, the specificity of the antibodies should be further validated by including conditions where the antibody is preabsorbed against a protein or peptide fragment(s) bearing the target sequence(s)/modification(s), or the antibody is omitted. Finally, it is essential to conduct a systematic analysis of the human and mouse proteome to determine which protein candidates are likely to cross-react with aSyn or aSyn-PTM-specific antibodies (Figure 9). Knowing these proteins and whether they are present in the samples of interest (cells, tissues, or biological fluids) could help eliminate potential confounding factors and accelerate the development of specific tools and assays to detect and quantify aSyn species.

### “Trust but verify”

Our experience with antibodies has taught us that verifying the specificity of the antibodies saves not only time and money but also projects. Antibody batch-to-batch or lot-lot variability is more common than most people think or realize. This is because most antibodies are not rigorously characterized and validated by vendors before their distribution to research laboratories and that validation data of new antibody batches are rarely made available. Most problems with antibodies are usually identified in research laboratories after significant time and resources have been wasted. During the past 5 years, we have had several experiences where we received lots of pS129 antibodies (from at least three different vendors) that did not exhibit the specificity and performance of the original lots. This contributes negatively to experimental reproducibility and could also lead to findings and controversies that distract and delay projects to come to fruition. For example, recently, we conducted systematic characterization of 16 antibodies that were reported and used as aSyn oligomer-specific antibody^91^. We showed that none of these antibodies could distinguish between aSyn oligomers and fibrils, which raises questions about many of the claims made about the levels or role of aSyn oligomers in the pathogenesis of synucleinopathies based on these antibodies.

Therefore, to address many of the problems outlined above, we propose that researchers and reviewers should demand that 1) all new antibodies are thoroughly characterized and validated under physiologically relevant conditions and using the appropriate protein standards (Figures 1A and S15); 2) the epitopes of all the antibodies should be disclosed, which will improve experimental design and enable more accurate interpretation of the results; 3) all new antibody batches are subjected to the same level of characterization and validation; 4) the resulting data are shared widely^92^, and all validation data is included in publications or vendors’ material data sheets; 5) all commercial antibodies are sold with the appropriate protein standards to allow the users to independently validate the antibodies in-house; and 6) the publication of negative results is encouraged.

Finally, funding agencies and publishers should incentivize and reward researchers who conduct studies and publish antibody characterization, validation, and reproducibility studies. Perhaps it is time to embrace the proposal of establishing certification programs for commercial antibodies^58^.

## Material and Methods

### Reagents and Antibodies

All the antibodies used in this study are described in Figure S3.

### Expression and purification of aSyn

pT7-7 plasmids were used for the expression of recombinant human aSyn (WT, 1-133, 1-135) in E.coli and purified as previously described^93,94^.

### In vitro protein fibrillization

Lyophilized full-length WT aSyn protein was dissolved for in vitro studies in Tris buffer (50mM Tris, 150 mM NaCl, pH 7.5, TBS) or phosphate-buffered saline (PBS, Invitrogen, Switzerland) for the cellular studies. The pH of the solution was adjusted to 7.2-7.5, and the protein was subsequently filtered through 0.22um polypropylene filters. Fibrils formation was induced by incubating the recombinant protein for 5 days under agitation (1’000 rpm) on an orbital shaker at 37°C. After incubation, the fibrils were fragmented by sonication (20 sec, 20% amplitude, 1X pulse on, and 1X pulse off) and separated from the remaining soluble protein content (before and after sonication) by following a published protocol^94^. Sonicated aSyn fibrils were aliquoted and stored at -80°C. As established by our lab^34,94,95^, aSyn fibrils were systematically and thoroughly characterized by SDS-PAGE and Coomassie staining, Thioflavin T aggregation assay and their length distribution were estimated before and after sonication by electron microscopy. Each of these assays is detailed in the sections below.

### Monomeric phosphorylated aSyn proteins

Phosphorylation of the recombinant (WT, 1-113, 1-135) and semisynthetic proteins (pY39, pS129, pY125, pS129 were produced and purified as described previously^27^.

### Nitration of aSyn fibrils

Post-fibrillization nitration of sonicated WT aSyn fibrils was induced with TNM (Sigma-Aldrich, Switzerland) following a published protocol^32^. Following nitration, the excess of TNM from nitrated fibrils was removed by washing steps in PBS using the 100 kDa MW-cut-off filters (Millipore, Switzerland) (13’000 rpm, 30 min, 4°C). The success of post-fibrillization nitration was confirmed by Liquid chromatography-mass spectrometry (ESI-LC-MS) and TEM.

### Phosphorylation of nitrated fibrils by PLK3

Phosphorylation of nitrated aSyn fibrils was induced following a previously published protocol^12^. Briefly, WT nitrated aSyn fibrils were dissolved in the phosphorylation reaction buffer, and PLK3 (0,1 μg) was added and incubated at 30°C without shaking. After 18h of incubation, the reaction was analyzed by LC-MS.

### Mass spectrometry analysis

Mass spectrometry analysis of proteins was performed by liquid chromatography-mass spectrometry (LC-MS) on an LTQ system (Thermo Scientific, San Jose, CA) equipped with an electrospray ion source and the MS instrument was operated in positive ion mode as previously described ^94^.

### Characterization of aSyn fibrils by Transmission electron microscopy (TEM)

To examine the ultrastructure of aSyn fibrils, negative staining electron microscope images were taken using Formvar/carbon-coated 200-mesh copper grids (Electron microscopy Sciences, Switzerland) as previously described^94^. Activated grids were loaded with 5 μl of sample for 30 seconds, washed three times with ultrapure water, and then negatively stained with 1% uranyl acetate for 1 min. Excess liquid was removed, and grids were allowed to air dry. Imaging was carried out on a Tecnai Spirit BioTWIN electron microscope operating at 80 kV acceleration voltage and equipped with a digital camera (FEI Eagle, FEI). A total of 3 to 5 images for each sample were chosen and the length of fibrils quantified (average 50-100 nm) using Image J software (U.S. National Institutes of Health, Maryland, USA; RRID:SCR_001935).

### Characterization of aSyn fibrils by Thioflavin T (ThT)

The amyloid-like properties and the extent of fibrilization of aSyn fibrils were assessed by ThT fluorescence as explained previously^93,94^. The sonicated aSyn fibrils were resuspended in ThT solution (50 mM glycine pH 8.5, 10 μM ThT solution). ThT fluorescence was measured with a FLUOstar plate reader (BMG Labtech, Germany) at an excitation wavelength of 450 nm and an emission wavelength of 485 nm.

### Characterization of aSyn fibrils by SDS-PAGE and Coomassie blue staining

The amount of soluble aSyn (monomers and oligomers) in the aSyn fibril solution was assessed before and after sonication after high-speed centrifugation by SDS-PAGE and Coomassie blue staining as previously described^94^. As sonication of aSyn fibrils can lead to the release of small amounts of monomers, only preparations with residual levels of aSyn monomers lower than 5 % were used for the seeding in primary neurons^34^.

### Slot Blot Analysis

Monomeric or fibrillar aSyn proteins unmodified or subjected to different PTMs such as nitration, phosphorylation, or nitration/phosphorylation were loaded to a final concentration of 36 ng into the Slot Blot system (GE Healthcare, Switzerland). The system was assembled with a pre-wetting nitrocellulose membrane and two Whatman filters in PBS for 10 min, and each slot was washed with 200 μl PBS. Each protein (100 μL) was then loaded onto the membrane through the slots. The slot blot system was then disassembled, and the membrane was removed. The membrane was soaked in blocking buffer (Intercept blocking buffer, LicCOR, Germany)) at room temperature (RT) for 1 h before being incubated with the different primary pS129 antibodies at a dilution of 1/250. After 1 h, the membranes were extensively washed three times with PBS buffer containing 0.1% of Tween 20 (PBS-T) and then incubated at RT for 1 h with the respective secondary goat anti-mouse or anti-rabbit antibodies conjugated to Alexa fluor 680 or 800 dyes. The source and dilution of each antibody can be found in Figure S3. The membranes were then washed three times with PBS-T, and scanned on a LiCOR scanner (LiCOR, Germany). Afterward, the same membranes were stained with SYN-1 or Eurogentec 1-20 total aSyn antibodies. All the experiments were independently repeated three times.

### ELISA

pS129 constructs (EPFL) and non-phosphorylated aSyn (Anaspec) were coated in a 96 well microplate (Nunc Maxisorp) at 0,1 μg/mL into a PBS buffer (Lonza) and incubated overnight at 4°C. Tau441 (rPeptide) was included as a negative control protein. The microplates were washed once (PBS/0,05% Tween 20) followed by adding 300 μL of PBS/0,1%casein (Thermo Scientific) to block aspecific reactions. After 2 hours at RT, the buffer was removed, and plates were dried overnight at RT. In the next step, the monoclonal pS129-aSyn antibodies (81A from BioLegend and MJF-R13 from Abcam) were diluted to 100 ng/mL into assay diluent (PBS/0,1% casein/0,2% Triton-X705) and 100 μL was added to the wells of the microplate plate followed by 1 hour at RT. A blank was included where assay buffer was added without antibody. After another wash step, the antibodies were detected with 100 μL of a tracer antibody for respectively mouse (81A) and rabbit (MJF-R13) IgG species purchased to Jackson Immunoresearch (Goat anti-mouse H&L: Cat# 115-035-166; Donkey anti-rabbit: Cat#711-035-152) at a 5000 fold working dilution. After 5 wash steps, another 100 μL of a colorimetric substrate was added (generic substrate buffer Euroimmun ELISA test kits) and incubated for 0,5 hours at RT, and the reaction was stopped by adding 50 μL of 1M sulphuric acid. The OD 450-600 nm was further analyzed in a microplate reader (BMG Clariostar). All data points were tested in duplicate, and average values are reported in the table and figures.

### Primary culture of hippocampal neurons and treatment with mouse aSyn fibrils

Primary hippocampal neurons were prepared from P0 pups of WT mice C57BL/6JRj (Janvier, France) or aSyn knock-out (KO) (C57BL/6J OlaHsd, Envigo, France) and cultured as previously described^34^. For high content imaging and immunocytochemistry (ICC), the neurons were seeded respectively in black, clear-bottomed, 96-well plates (Falcon, Switzerland) at a density of 200,000 cells/mL or onto coverslips (CS) (VWR, Switzerland) at a density of 250,000 cells/mL. For WB analysis, the neurons were plated on 6-wells plates at a density of 300,000 cells/mL. All the plates were coated with poly-L-lysine 0.1% w/v in water (Brunschwig, Switzerland) before the plating. For the neuronal seeding model, the WT neurons after 6 days in culture (DIV 6) were treated with extracellular mouse aSyn PFFs at a final concentration of 70 nM and cultured for 14 or 21 days as previously described^31,33,34,69^. The mouse aSyn PFFs used in this study are from the same batch prepared and characterized in Mahul-Mellier et al., BioRxiv 2018^31^. The E83Q aSyn PFFs used in this study are from the same batch prepared, characterized and used in Kumar et al., BioRxiv 2021^36^. All procedures were approved by the Swiss Federal Veterinary Office (authorization numbers VD 3392 and VD 3496).

### Stereotaxic Injection of mouse aSyn fibrils

Male C57BL/6JRj mice (two to three months of age, three animals per cage) were housed at 23°C, 40% humidity, with dark/light cyle of 12 hours from 7 A.M to 7 P.M. with access to standard laboratory rodent chow and water for *in vivo* experiments ad libitum. All animal experimentation procedures were approved by the Cantonal Veterinary Authorities (Vaud, Switzerland) and were performed according to the European Communities Council Directive of 24 November 1986 (86/609EEC). Every effort was taken to minimize the number of animals and their stress. Mice (n=12) have been injected with 2.5 uL of mouse PFFs, prepared as previously described^94,96^ in the right dorsal striatum (AP: 0.6, ML: 2.0, DV: -2.6). In detail, animals were anesthetized using a Ketamine/Xylazine cocktail. Once fully anesthetized, the animal was placed on the stereotaxic frame, head shaved, viscotears was applied to his eyes and liquid betadine solution was applied on the scalp before incision. Skull was cleaned from connective tissue with a scalpel and cleaned to highlight the bregma. Without damaging the dura mater, a small hole was drilled into the skull. Once in location, 2 μl of fibrils were injected in the right hemisphere at a flow rate of 0.4 μl/min. The needle was kept in place for 5 minutes and then slowly retracted and cleaned from blood to avoid clogging. The animal was then sutured, betadine gel applied on the wound and kept in a heated observation cage until fully awake.

### Mammalian cell lines culture

HEK293 and HeLa cells were cultured at 95% air, and 5% CO2 in DMEM (for HEK293 and HeLa) supplemented with 10% Fetal bovine serum (Invitrogen, Switzerland) and 1% of Penicillin/Streptomycin (Invitrogen, Switzerland). The plasmid transfections were carried out by using the Effectene transfection reagent (Qiagen, Germany) according to the manufacturer’s protocol. Mammalian cells were transfected with either the empty vector (pAAV) or human WT aSyn pAAV vector (Addgene plasmid # 36055; http://n2t.net/addgene:36055 ; RRID:Addgene_36055).

### Subcellular fractionation of aSyn KO and WT primary neurons, brain homogenates and mammalian cell lines

#### Preparation of soluble and insoluble fractions from aSyn KO and WT primary neurons and from brain homogenates

At the indicated time, WT or aSyn KO primary cortical or hippocampal neurons (untreated or treated with WT aSyn mouse PFFs or with PBS as negative control)^31,33,34,69^ or cortexes, striata, amygdala and midbrains from aSyn KO mouse brains were lysed in 1% Triton X-100/ Tris-buffered saline (TBS) (50 mM Tris, 150 mM NaCl, pH 7.5) supplemented with protease inhibitor cocktail (Sigma-Aldrich, Switzerland), 1 mM phenylmethane sulfonyl fluoride (PMSF) (Sigma-Aldrich, Switzerland), and phosphatase inhibitor cocktails 2 and 3 (Sigma-Aldrich, Switzerland). Sequential biochemical fractionation of cell extracts was performed as described previously^31,34,69^. Cell lysates were sonicated using a fine probe [(0.5-s pulse at an amplitude of 20%, ten times (Sonic Vibra Cell, Blanc Labo, Switzerland], and then incubated on ice for 30 min before being centrifuged at 100,000 g for 30 min at 4°C. The supernatant (soluble fraction) was collected. The pellet was washed in 1% Triton X-100/TBS and sonicated [(0.5-s pulse at an amplitude of 20%, ten times (Sonic Vibra Cell, Blanc Labo, Switzerland] before being centrifuged for 30 min at 100,000 g. The supernatant was then discarded, and the pellet (insoluble fraction) was resuspended in 2% sodium dodecyl sulfate (SDS) diluted in TBS, pH 7.4 and finally sonicated using a fine probe [(0.5-s pulse at an amplitude of 20%, ten times (Sonic Vibra Cell, Blanc Labo, Switzerland].

#### Preparation of the total cell lysates from aSyn KO and WT primary neurons

Primary hippocampal or cortical neurons were lysed in 2% SDS/TBS supplemented with protease inhibitor cocktail, 1 mM PMSF, and phosphatase inhibitor cocktail 2 and 3 (Sigma-Aldrich, Switzerland) and boiled for 10 min as previously described^34^.

#### Preparation of the nuclear, cytosolic and membrane fractions from aSyn KO and WT primary neurons and mammalian cell lines

Subcellular fractionation of the HEK and HeLa mammalian cells lines or the primary neurons was performed as previously described^97,98^ and from Abcam protocol (https://www.abcam.com/protocols/subcellular-fractionation-protocol) with minor modifications. Briefly, cells cultured in 10 cm culture plate (∼5 million cells) were washed twice with ice-cold PBS and scraped with 500 μL fractionation buffer (20 mM HEPES at pH 7.4, 10 mM KCI, 2 mM MgCl2, 1 mM EDTA, 1 mM EGTA) supplemented extemporaneously with 1mM of DTT, protease inhibitor cocktail (Sigma-Aldrich, Switzerland), 1 mM phenylmethane sulfonyl fluoride (PMSF) (Sigma-Aldrich, Switzerland), and phosphatase inhibitor cocktail 2 and 3 (Sigma-Aldrich, Switzerland). The cells in suspension were then incubated for 15 min on ice before being lysed mechanically using a Dounce homogenizer (with type B pestle). After 20 strokes, the cell homogenate was left 20 min on ice before being centrifuged at 720 g at 4°C for 5 min. The supernatant (cytosolic and membranes fractions), which contains the cytoplasm, membranes, and mitochondria, was then transferred into a fresh tube and kept on ice for 20 min.

The pellet corresponding to the nuclear fraction was washed with 500 μL fresh fractionation buffer to disperse the pellet. The resulting suspension was poured into a Dounce homogenizer (with type B pestle) and subjected to 20 strokes followed by centrifugation at 720 g at 4°C for 10 min. The supernatant was discarded, and the final pellet (nuclear fraction) was resuspended in TBS supplemented with 0.1% SDS and sonicated briefly (3s on ice at a power setting of 60%).

The cytosolic fraction was isolated from the membrane fraction after ultracentrifugation of the supernatant at 100,000 g at 4°C for 1 h. The supernatant was collected and kept as the cytosolic fraction. The final pellet was washed by adding 400 μL of fractionation buffer, resuspended, and passed through Dounce homogenizer (with type B pestle-20 strokes) followed by a last ultracentrifugation step 100,000 g at 4°C for 45 min. The pellet contains the membranes fraction which was resuspended in TBS (50 mM Tris-HCl-pH 7.6, 150 mM NaCl) supplemented with 0.1% SDS.

### Tricine gels and Western blot analyses

The bicinchoninic acid (BCA) protein assay was performed to quantify the protein concentration in all the subcellular fractions (soluble, insoluble, nuclear, cytosolic, membrane and total cell extract) according to the supplier’s protocol (Pierce, Thermofisher, Switzerland). Laemmli buffer 4× (10% SDS, 50% glycerol, 0.05% bromophenol blue, 1M Tris-HCl pH 6.8 and 20% β-mercaptoethanol) was added to all the fractions. 20 to 30 μg of proteins for each fraction were then separated on a 16% tricine gel^99^. 0.5 μl of the PageRuler prestained protein ladder (10 to 180 kDa) from Thermofisher (Switzerland) was loaded for each gel. After two hours of running, the gels were then transferred onto a nitrocellulose membrane (Amersham, Switzerland) with a semi-dry system (Bio-Rad, Switzerland), and immunoblotting was performed as previously described^31,34^.

### Immunocytochemistry (ICC)

WT and aSyn KO primary hippocampal neurons untreated or treated with WT aSyn mouse^31^ or E83Q human PFFs^36^ were washed twice with PBS, fixed in 4% PFA for 20 min at RT, and then immunostained as previously described^31,34,95^. The antibodies used are indicated in the corresponding legend section of each figure. The source and dilution of each antibody can be found in Figure S3. The cells plated on CS were then examined with a confocal laser-scanning microscope (LSM 700, Carl Zeiss Microscopy, Germany) with a 40× objective and analyzed using Zen software (RRID:SCR_013672). The cells plated in black, clear-bottom, 96-well plates were imaged using the IN Cell Analyzer 2200 (with a ×10 objective). For each independent experiment, two wells were acquired per tested condition, and in each well, nine fields of view were imaged. Each experiment was reproduced at least three times independently.

### Quantitative high-content wide-field cell imaging analyses (HCA)

After WT mouse aSyn PFFs treatment for 10 days, the primary hippocampal neurons plated in black, clear-bottom, 96-well plates (BD, Switzerland) were washed twice with PBS, fixed in 4% paraformaldehyde (PFA) for 20 min at RT and then immunostained as described above. Images were acquired using the Nikon 10×/ 0.45, Plan Apo, CFI/60 of the IN Cell Analyzer 2200 (GE Healthcare, Switzerland), a high-throughput imaging system equipped with a high-resolution 16-bits sCMOS camera (2048×2048 pixels), using a binning of 2×2. For each independent experiment, duplicated wells were acquired per condition, and nine fields of view were imaged for each well. Each experiment was reproduced at least three times independently. Images were then analyzed using Cell profiler 3.0.0 software (RRID:SCR_007358) for identifying and quantifying the level of LB-like inclusions (stained with pS129 antibody) formed in neurons MAP2-positive cells, the number of neuronal cell bodies (co-stained with MAP2 staining and DAPI), or the number of neurites (stained with MAP2 staining). The pipeline of analysis is fully described in Mahul-Mellier et al., 2020^34^.

### Immunochemistry on brain slides

Adult aSyn KO (C57BL/6J OlaHsd, Envigo) and Wild Type (C57BL6/J, Janvier labs) mouse fibrils-injected mice (3 months post-injection) were anesthetized using a solution containing 100 mg/Kg of Ketamine and 10 mg/Kg of xylazine. Once fully anesthetized, animals were perfused with PBS and then 4% PFA in PBS. Brains were rapidly extracted and left in PFA 4% at 4° for 24 hours, after that, they were put in a 30% sucrose solution in PBS again at 4° for 24 hours. Brains were then frozen in isopentane, stored at -80°, and subsequently cut into 50 μm-thick coronal slides with seriality of one every four sections and stored in a PBS-azide 0,1% solution at 4° until further processing. Free-floating coronal slices from 5 different animals were rinsed 3 times for 5 minutes in PBS and then incubated 10 minutes in a 10% hydrogen peroxide solution in PBS, then washed again twice for 5 minutes in PBS, left in a 2.5% Normal Horse Serum blocking solution for 30 minutes (ImmPRESS, Vector laboratories, Germany) and then incubated overnight at 4° shaking with primary antibodies (Figure S3) diluted in blocking solution. The following day, slices were rinsed 3 times in PBS and then incubated 1 hour at RT with either ImmPRESS® HRP Horse Anti-Mouse IgG Polymer (Vector Laboratories Cat# MP-7402, RRID:AB_2336528) or ImmPRESS® HRP Horse Anti-Rabbit IgG Polymer (Vector Laboratories Cat# MP-7401, RRID:AB_2336529), according to primary antibody’s host species. After incubation, slices were rinsed again 4 times for 5 minutes in PBS and revealed using a DAB substrate kit (Vector Laboratories Cat# SK-4100, RRID:AB_2336382). Stained slices (at least 5 per region) were then mounted on gelatinized slides, coverslipped and imaged at 20x using an VS120-SL Olympus slide scanner microscope (Olympus, Germany). All procedures were approved by the Swiss Federal Veterinary Office (authorization number VD2067).

### Identification of putative antibody cross-reaction candidates

All human and mouse phosphorylation sites annotated in Uniprot were retrieved as full text entries using the https://www.uniprot.org server on June 15^th^, 2020 by entering the following terms in the search box for UniProtKB: (annotation:(type:mod_res phosphoserine) OR annotation:(type:mod_res phosphotyrosine)) (organism:”Homo sapiens (Human) [9606]” OR organism:”Mus musculus (Mouse) [10090]”) AND reviewed:yes

A perl script was used to parse the entries, extract protein sequence fragments of 13 residues centered on each documented annotated phosphorylation site and compare them to the six human aSyn fragments shown in Figure 9A. Residues immediately preceding and following the phosphorylation site were requested to be perfect matches. For all protein fragments retrieved from Uniprot satisfying this condition (Figure S13), we computed the number of exact and highly similar matches to the six aSyn fragments and the length of the longest continuous stretch of perfect or similar matches encompassing the phosphorylation site. High similarity criteria were defined as a PAM120 matrix score of two matching residues strictly greater than

1. This would include for example Glu ≅ Gln (score = 2) and Tyr ≅ Phe (score = 4) but exclude Ile ≠Leu (score = 1) and Ile ≠Phe (score = 0). For each of the six aSyn fragments, hits were sorted by the number of continuous exact match stretch, then the number of continuous exact+similar match stretch, then the number of exact matches, then the number of similar matches, then the Molecular Weight.

Subcellular location annotations were retrieved by parsing the “CC -!- SUBCELLULAR LOCATION:” fields of the corresponding Uniprot entries.

Three online databases ie. The Human Protein Atlas, https://www.proteinatlas.org/; Proteomics DB, https://www.proteomicsdb.org/proteomicsdb/; CSF proteome resource, https://proteomics.uib.no/csf-pr/ were used to evaluate if the proteins were previously identified in Human plasma or CSF.

## Acknowledgments

We thank the CIME facility (EPFL) for using their electron microscopy facility and Dr. Fabien Kuttler (PTCB, EPFL) for developing the original HCA pipeline.

## Author contributions

Conceptualization: HAL

Experimental design: HAL

Methodology: ALMM, SN, RNH, YJ, FA, SD, SMG, RB, JR, MB, PM, ES, CI, AC

Investigation: HAL, ALMM, SN, RNH, YJ, FA, SD, SMG, RB, JR, MB, PM, ES, CI, AC

Data curation: HAL, ALMM, SN, RNH, YJ, FA, SD, SMG, RB, JR, MB, PM, ES, CI, AC

Formal analysis: HAL, ALMM, SN, RNH, FA, SD, SMG, RB, PM, ES, CI

Validation: HAL, ALMM, SN, RNH, FA, SD, SMG, RB, PM, ES, CI

Visualization: HAL, ALMM, SN, RNH, YJ, FA, SD, SMG, RB, JR, MB, PM, AS, ES, CI, AC

Software: PM, ES, CI

Supervision: HAL

Writing—original draft: HAL

Writing—review & editing: HAL, ALMM, SN, PM

Funding acquisition: HAL

Project administration: HAL

Resources: HAL

## Data and materials availability

All data are available in the main text or supplementary materials.

