## Supplemental Information for "Neighbouring modifications interfere with the detection of phosphorylated alpha-synuclein at Serine 129: Revisiting the specificity of pS129 antibodies"

**Affiliations:**

**Running title:** Revisiting the specificity of pS129 antibodies

**Keywords:** Synuclein, phosphorylation, antibodies, aggregation, posttranslational modifications.

**Legends**

**Figure S1**

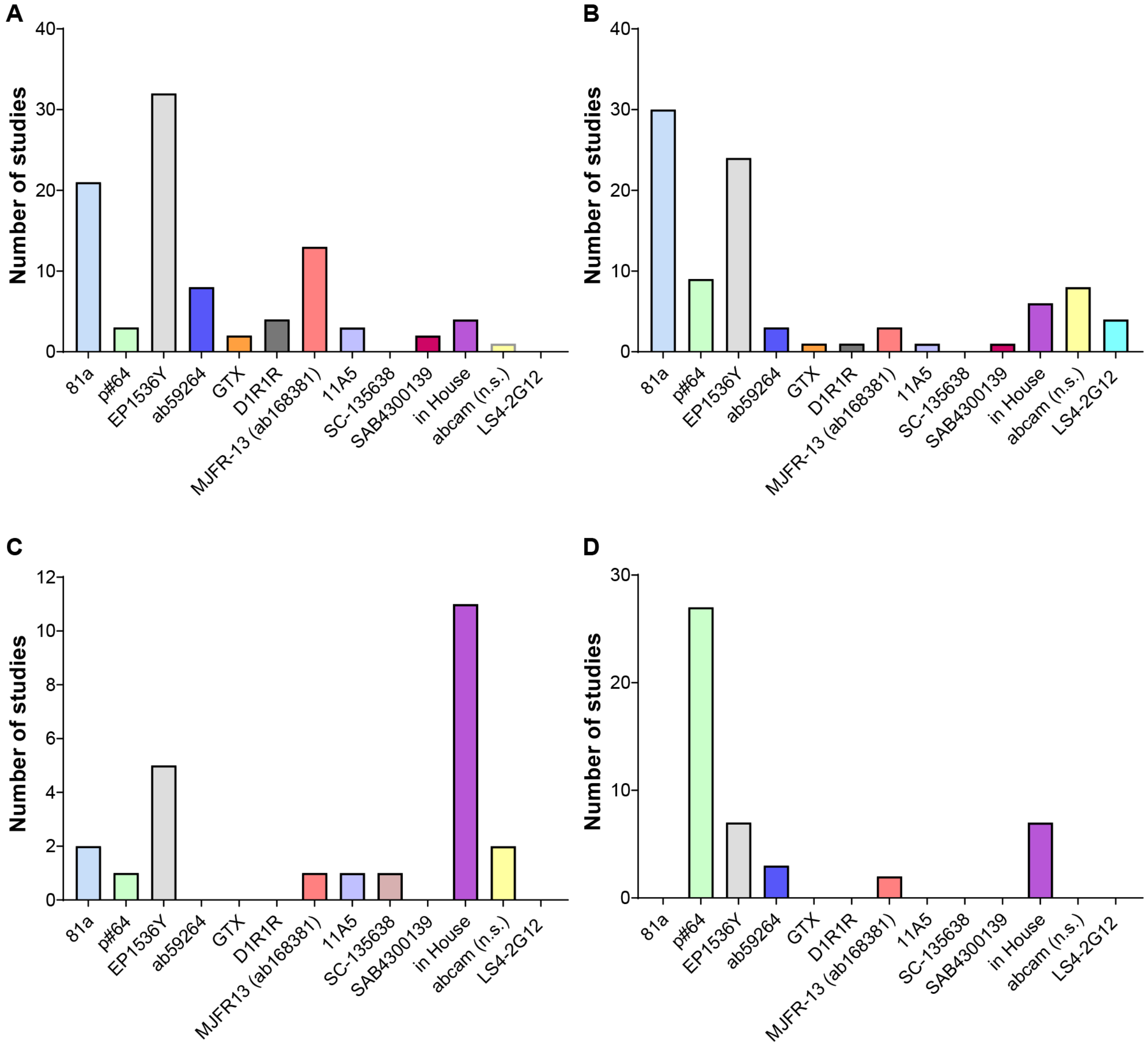

#### Figure S1

##### E. Seeding model cell culture studies

| References | Year of publication | pS129 antibody(-ies) used |
|---|---|---|
| **Luk et al.** | 2009 | 81A |
| **Volpicelli-Daley et al.** | 2011 | 81A |
| **Tanik et al.** | 2013 | 81A |
| **Aulic et al.** | 2014 | MJF-R13 (ab168381);EP1536Y; ab59264 |
| **Tran et al.** | 2014 | 81A |
| **Volpicelli-Daley et al.** | 2014 | 81A; EP1536Y |
| **Ma et al.** | 2016 | MJF-R13 (ab168381);EP1536Y; ab59264 |
| **Mao et al.** | 2016 | 81A; MJF-R13 (ab168381) |
| **Samuel et al.** | 2016 | 11A5 |
| **Abdelmotilib et al.** | 2017 | 81A |
| **Karampetsou et al.** | 2017 | p#64, EP1536Y |
| **Karpowicz et al.** | 2017 | 81A; in-house |
| **Tapias et al.** | 2017 | EP1536Y |
| **Froula et al.** | 2018 | 81A; EP1536Y |
| **Grassi et al.** | 2018 | MJF-R13 (ab168381); GTX |
| **Henderson et al.** | 2018 | MJF-R13 (ab168381); in-house |
| **Mahul-Mellier et al.** | 2018 | 81A; MJF-R13 (ab168381) |
| **Peng et al.** | 2018 | 81A |
| **Polinski et al.** | 2018 | 81A |
| **Sorrentino et al.** | 2018 | EP1536Y |
| **Terada et al.** | 2018 | p#64; EP1536Y |
| **Wang et al.** | 2018 | 81A; MJF-R13 (ab168381); EP1536Y |
| **Chen et al.** | 2019 | anti-pS129 (Abcam) (n.s) |
| **Elfarrash et al.** | 2019 | D1R1R;11A5 |
| **Froula et al.** | 2019 | EP1536Y |
| **Gao et al.** | 2019 | MJF-R13 (ab168381); EP1536Y |
| **Grassi et al.** | 2019 | GTX |
| **Gribaudo et al.** | 2019 | 81A; EP1536Y; ab59264 |
| **McGlinchey et al.** | 2019 | ab59264 |
| **Volpicelli-Daley et al.** | 2019 | EP1536Y |
| **Wang et al.** | 2019 | EP1536Y |
| **Wu et al.** | 2019 | 81A |
| **Ardah et al.** | 2020 | EP1536Y |
| **Bengoa-Vergniory et al.** | 2020 | EP1536Y |
| **Challis et al.** | 2020 | ab59264 |
| **Courte et al.** | 2020 | EP1536Y; ab59264; D1R1R |
| **Gegg et al.** | 2020 | EP1536Y |
| **Guan et al.** | 2020 | p#64; in-house |
| **Henderson et al.** | 2020 | MJF-R13 (ab168381); EP1536Y |
| **Henrich et al.** | 2020 | EP1536Y |
| **Mahul-Mellier et al.** | 2020 | 81A; MJF-R13 (ab168381); SAB4300139 |
| **Morgan et al.** | 2020 | EP1536Y |
| **Apicco et al.** | 2021 | EP1536Y |
| **Burbidge et al.** | 2021 | MJFR13 (ab168381) |
| **Courte et al.** | 2021 | EP1536Y; ab59264; D1R1R |
| **Elfarrash et al.** | 2021 | D1R1R;11A5 |
| **Emmenegger et al.** | 2021 | 81A; EP1536Y |
| **Mahul-Mellier et al.** | 2021 | 81A; MJF-R13 (ab168381); SAB4300139 |
| **Marotta et al.** | 2021 | 81A |
| **Pantazopoulou et al.** | 2021 | EP1536Y |
| **Stykel et al.** | 2021 | 81A; EP1536Y |
| **Tanudjojo et al.** | 2021 | EP1536Y; in-house |
| **Trinkaus et al.** | 2021 | EP1536Y |
| **Vajhøj et al.** | 2021 | EP1536Y |
| **Zhang et al.** | 2021 | EP1536Y |
| **Awa et al.** | 2022 | 81A; EP1536Y |
| **Vasili et al.** | 2022 | MJF-R13 (ab168381);EP1536Y; ab59264 |

### Figure S1

#### F. in vivo and mouse strains studies

| References | Year of publication | pS129 antibody(-ies) used |
|---|---|---|
| **Luk et al.** | 2012a | In-house (pSyn 6.2); in-house (SynpSer129) |
| **Luk et al.** | 2012b | In-house (pSyn 6.2); in-house (SynpSer129) |
| **Masuda-Suzukake et al.** | 2013 | p#64 , in house (1175 provided by Akiyama) |
| **Guo et al.** | 2013 | 81A |
| **Tran et al.** | 2014 | 81A |
| **Sacino et al.** | 2014A | 81A |
| **Sacino et al.** | 2014B | 81A |
| **Sacino et al.** | 2014C | 81A |
| **Osterberg et al.** | 2015 | 81A |
| **Paumier et al.** | 2015 | 81A |
| **Mao et al.** | 2016 | MJF-R13 (ab168381); SAB4300139; 81A |
| **Abdelmotilib et al.** | 2017 | 81A |
| **Harms et al.** | 2017 | 81A |
| **Sorrentino et al.** | 2017 | 81A; EP1536Y |
| **Karampetsou et al.** | 2017 | not specified; in-house |
| **Duffy et al.** | 2018 | 81A |
| **Luna et al.** | 2018 | 81A |
| **milanese et al.** | 2018 | ab59264 |
| **Uemura et al.** | 2018 | 81A, p#64, EP1536Y |
| **Yun et al.** | 2018 | 81A; MJF-R13 (ab168381) |
| **Ayers et al.** | 2018 | LS4-2G12 |
| **Sorrentino et al..** | 2018 | LS4-2G12 |
| **Okuzumi et al.** | 2018 | ab59264 |
| **Rey et al.** | 2018 | EP1536Y |
| **Lee et al.** | 2018 | MJF-R13 (ab168381) |
| **Manfredsson et al.** | 2018 | 81A |
| **Ren et al..** | 2019 | anti-pS129 (Abcam) (n.s) |
| **Durante et al.** | 2019 | EP1536Y |
| **Mavroeidi et al.** | 2019 | EP1536Y; 11A5; LS4-2G12 |
| **Patterson et al.** | 2019 | 81A |
| **Earls et al.** | 2019 | EP1536Y |
| **Chaterjee et al.** | 2019 | EP1536Y |
| **Bieri et al.** | 2019 | EP1536Y |
| **Yabuki et al.** | 2020 | anti-pS129 (Abcam) (n.s); p#64 |
| **Zhang et al.** | 2020 | p#64 |
| **Jia et al.** | 2020 | EP1536Y |
| **Kim et al.** | 2020 | 81A |
| **Kim et al.** | 2020 | EP1536Y |
| **Creed et al.** | 2020 | 81A |
| **Schaser et al.** | 2020 | 81A; EP1536Y |
| **Ham et al.** | 2020 | p#64 |
| **Wu et al.** | 2020 | 81A |
| **Howe et al.** | 2021 | MJFR13 (ab168381); GTX50222 |
| **Underwood et al.** | 2021 | EP1536Y |
| **Xu et al.** | 2021 | anti-pS129 (Abcam) (n.s) |
| **Bergkvist et al.** | 2021 | EP1536Y |
| **Ferreira et al.** | 2021 | 81A, EP1536Y, and LS4-2G12 |
| **Marotta et al.** | 2021 | EP1536Y |
| **Sykioti et al.** | 2021 | EP1536Y |
| **Demmings et al.** | 2021 | anti-pS129 (Abcam) (n.s) |
| **Lai et al.** | 2021 | EP1536Y |
| **Kweon et al.** | 2021 | anti-pS129 (Abcam) (n.s) |
| **Zhang et al.** | 2021 | p#64 |
| **Dutta et al.** | 2021 | anti-pS129 (Abcam) (n.s) |
| **Lee et al.** | 2021 | 81A |
| **Ghosh et al.** | 2021 | 81A; ab59264 |
| **Apicco et al.** | 2021 | EP1536Y |
| **Kuan et al.** | 2021 | 81A |
| **Zhang et al.** | 2021 | EP1536Y |
| **George et al.** | 2021 | anti-pS129 (Abcam) (n.s) |
| **Cole et al.** | 2021 | 81A |
| **Gentzel et al.** | 2021 | p#64 |
| **Ma et al.** | 2021 | 81A |
| **Thomsen etal** | 2021 | D1R1R |
| **Jimenez-ferrer et al.** | 2021 | EP1536Y |
| **Arkan et al.** | 2021 | EP1536Y |
| **Pantayopoulou et al.** | 2021 | EP1536Y |
| **Merghani et al.** | 2021 | anti-pS129 (Abcam) (n.s) |
| **Szego et al.** | 2021 | p#64 |
| **Boutros et al.** | 2021 | EP1536Y |
| **Okuda et al.** | 2022 | p#64; EP1536Y |
| **Thakur et al.** | 2017 | 81A; pSyn 5038 (Ser129) EMD (Millipore)* |
| **Kakoty 2021** | 2021 | LB509 (ab27766) * |

#### Figure S1

##### G. Bodyfluids biomarker-based studies

| References | Year of publication | pS129 antibody(-ies) used |
|---|---|---|
| **Foulds et al.** | 2011 | EP1536Y; in house |
| **Wang Y et al.** | 2012 | anti-pS129 (Abcam) (n.s); in house |
| **Foulds et al.** | 2012 | EP1536Y; in house |
| **Foulds et al.** | 2013 | EP1536Y |
| **Stewart et al.** | 2015 | In house |
| **Majbour et al.** | 2016a | In house |
| **Majbour et al.** | 2016b | In house |
| **van Steenoven et al.** | 2018 | In house |
| **Cariulo et al.** | 2019 | MJF-R13 (8-8) (ab168381) |
| **Lin et al.** | 2019 | 81A |
| **Elhadi et al.** | 2019 | pSyn#64 (WAKO) |
| **Tian et al.** | 2019 | 81A |
| **Majbour et al.** | 2020 | In house |
| **Constantinides et al.** | 2020 | In house |
| **Li et al.** | 2020 | sc-135638 |
| **Majbour et al.** | 2021 | In house |
| **Schulz et al.** | 2021 | PRTA-11A5; PRTA-23E8 |
| **Chen et al.** | 2021 | Anti-pS129 (Abcam) (n.s.) |

##### H. Gastrointestinal studies

| References | Year of publication | pS129 antibody(-ies) used |
|---|---|---|
| **Lebouvier et al.** | 2008 | pSyn#64 |
| **Beach et al.** | 2010 | in-house |
| **Lebouvier et al.** | 2010 | pSyn#64 |
| **Pouclet et al.** | 2011 | pSyn#64 |
| **Boettner et al.** | 2012 | pSyn#64 |
| **Pouclet et al.** | 2012 | pSyn#64 |
| **Pouclet et al.** | 2012 | pSyn#64 |
| **Beach et al.** | 2013 | in-house |
| **Folgoas et al.** | 2013 | pSyn#64 |
| **Adler et al.** | 2014 | in-house |
| **Gelpi et al.** | 2014 | pSyn#64 |
| **Gray et al.** | 2014 | ab59264 |
| **Hilton et al.** | 2014 | pSyn#64 |
| **Ito et al.** | 2014 | pSyn#64, in-house |
| **Masuda et al.** | 2014 | pSyn#64 |
| **Aldecoa et al.** | 2015 | pSyn#64 |
| **Clairembault et al.** | 2015 | pSyn#64 |
| **Mu et al.** | 2015 | pSyn#64 |
| **Sanchez-Ferro et al.** | 2015 | pSyn#64 |
| **Sprenger et al.** | 2015 | pSyn#64 |
| **Visanji et al.** | 2015 | ab59264 |
| **Adler et al.** | 2016 | in-house |
| **Antunes et al.** | 2016 | ab59264 |
| **Beach et al.** | 2016 | in-house |
| **Chung et al.** | 2016 | EP1536Y |
| **Corbille et al.** | 2016 | pSyn#64, in-house |
| **Stokholm et al.** | 2016 | MJF-R13 |
| **Vilas et al.** | 2016 | pSyn#64 |
| **Barrenschee et al.** | 2017 | pSyn#64 |
| **Carletti et al.** | 2017 | pSyn#64 |
| **Corbille et al.** | 2017 | pSyn#64 |
| **Kim et al.** | 2017 | pS129-aSyn (Santa Cruz) |
| **Rouaud et al.** | 2017 | pSyn#64 |
| **Shin et al.** | 2017 | EP1536Y |
| **Beach et al.** | 2018 | MJF-R13 |
| **Fernandez-Arcos et al.** | 2018 | pSyn#64 |
| **Iranzo et al.** | 2018 | pSyn#64 |
| **Killinger et al.** | 2018 | EP1536Y |
| **Lee et al.** | 2018 | EP1536Y |
| **Ruffmann et al.** | 2018 | pSyn#64 |
| **Shin et al.** | 2018 | EP1536Y |
| **Fenyi et al.** | 2019 | pSyn#64 |
| **Leclair-Visonneau et al.** | 2019 | pSyn#64 |
| **Shin et al.** | 2019 | EP1536Y |

**Figure S1. pS129 antibodies used across human studies and animal and cellular seeding models (related to Figures 1 to 8)**

**A-D.** Graphs represent the pS129 antibodies used across the different number of reports in the cell culture studies using the seeding model (**A**), in the animal studies using the seeding model (**B**), in the bodyfluids biomarker-based studies (**C**), and in the gastrointestinal tract (GIT) studies (**D**). In-house corresponds to the pS129 antibodies generated from different individual groups. **E-H.** Tables list the individual studies that used pS129-specific antibodies displayed in the graphs **A-D,** in the cell culture studies using the seeding model (**E**), in the animal studies using the seeding model (**F**), in the bodyfluids biomarker-based studies (**G**) and in GIT studies (**H**). In-house corresponds to the pS129 antibodies generated from different individual groups. Abcam (n.s.): catalogue number was not provided in the original manuscript. The red asterisk indicates studies in which total aSyn antibodies have been used to detect pS129 pathology.

#### Figure S2

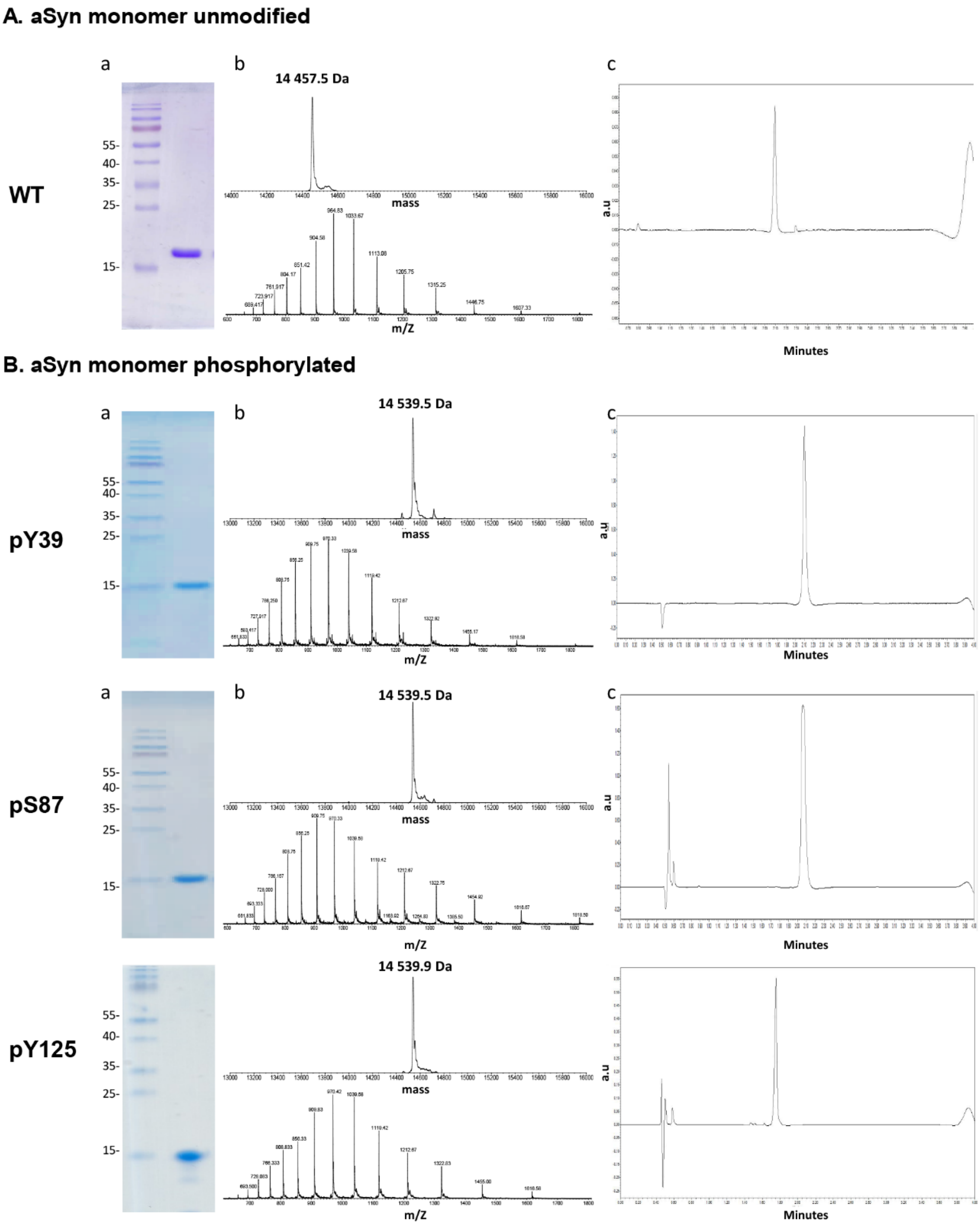

A. aSyn monomer unmodified
a
b
c
WT
55-
40-
35-
25-
15-
14 457.5 Da
mass
m/Z
a.u
Minutes
B. aSyn monomer phosphorylated
pY39
14 539.5 Da
pS87
14 539.5 Da
pY125
14 539.9 Da

#### Figure S2

##### C. aSyn monomer phosphorylated (semi-synthetic)

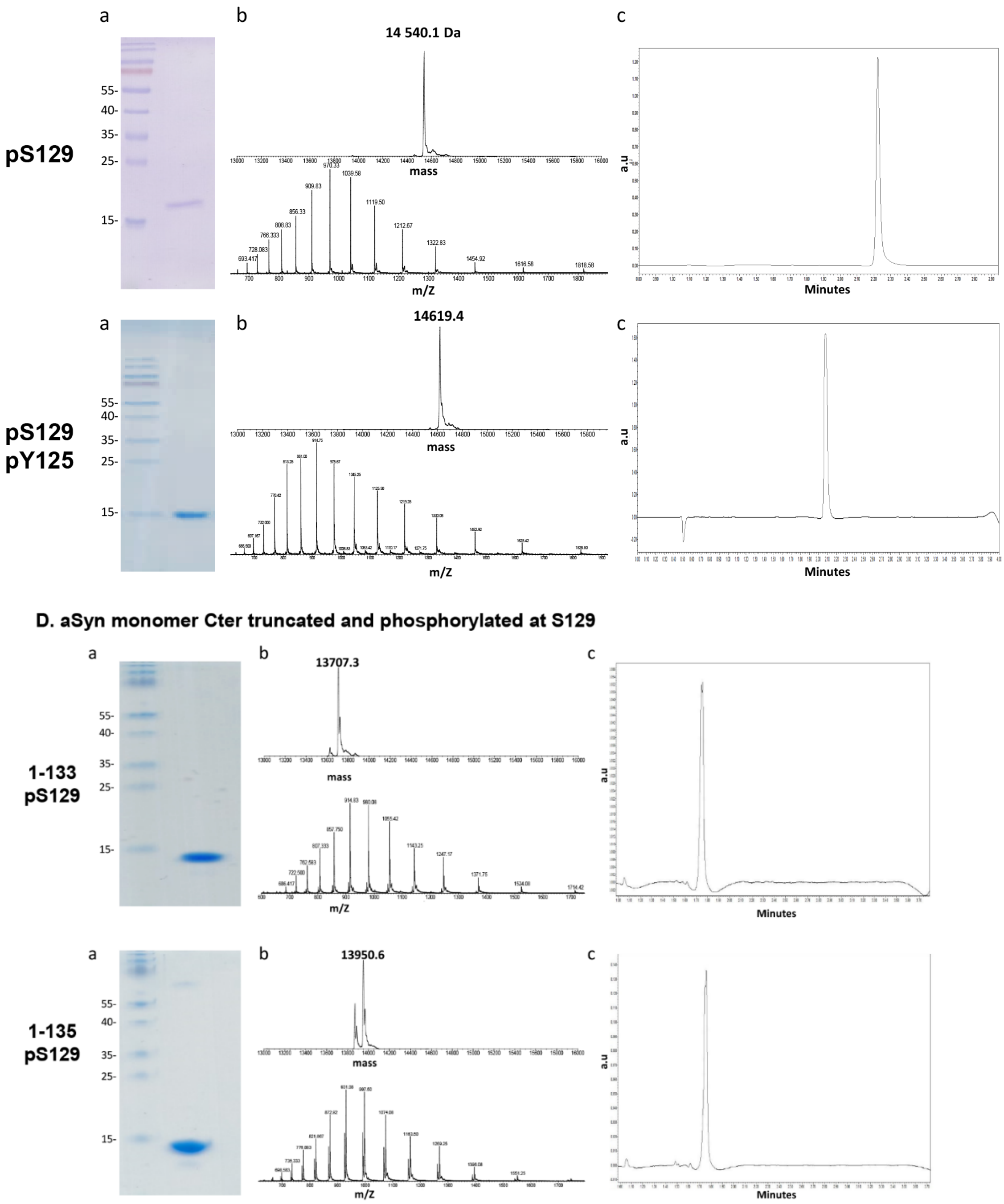

**Figure S2. Preparation and characterization of recombinant monomeric aSyn species (related to Figures 1 to 8)**

WT and mutants human aSyn recombinant proteins were produced in *E. coli* and purified by anion exchange chromatography and size-exclusion chromatography, followed by a final chromatographic step using reverse-phase HPLC, as previously described[1]. The purity of recombinant monomeric aSyn was systematically assessed by SDS-PAGE and Coomassie staining (a), ESI-LC/MS (b), that also showed the expected mass and UPLC (c). **A.** Human aSyn unphosphorylated (WT). **B.** Human aSyn phosphorylated at Y39 (pY39), S87 (pS87) or Y125 (pY125). **C.** Human aSyn phosphorylated at Y125 or S129 (pS129) residues or diphosphorylated (pS129 pY125). **D.** Human aSyn C-terminally truncated after the residue 133 (1-133) or 135 (1-135) and phosphorylated at S129.

#### Figure S3

**A**

| Primary Antibody | Catalog # | Company | Clone | RRID | Host | Concentration | WB dilution | ICC dilution | Epitope |
|---|---|---|---|---|---|---|---|---|---|
| anti-aSyn **total** | sc-10717 | Santa Cruz | FI-140 | - | Rabbit | 0.2 mg/mL | 1:500 | - | **1-140** |
| anti-aSyn **total** | 610787 | BD | SYN-1 | RRID:AB_398108 | Mouse | 0.25 mg/mL | 1:1000 | 1:1000 | **91-99** |
| anti-aSyn **total** | 4179 | Cell signalling | D37A6 | RRID:AB_1904156 | Rabbit | 1 mg/mL | 1:1000 | - | **Near Glu105** |
| anti-aSyn **total** | ab131508 | Abcam | - | RRID:AB_11155736 | Rabbit | 1 mg/mL | 1:1000 | 1:500 | **134-138** |
| anti-aSyn **total** | LASHUEL | - | LASH-EGT1-20 | - | Rabbit | Not provided | 1:1000 | 1:1000 | **1-20** |
| anti-aSyn **total** | LASHUEL | - | LASH-BL-A15110B | - | Mouse | | 1:1000 | 1:500 | **34-45** |

**B**

| Primary Antibody | Catalog # | Company | Clone | RRID | Host | Concentration | WB dilution | ICC dilution | HTS | IHC | Epitope |
|---|---|---|---|---|---|---|---|---|---|---|---|
| anti-**pS129**-aSyn | 825701 | BioLegend | p-syn/81A | RRID:AB_2564891 | Mouse | 1.0 mg/mL | 1:1000 | 1:2000 | 1:1000 | 1:10000 | Peptide (residues 124-134) including phosphorylated Ser129 of human aSyn |
| anti-**pS129**-aSyn | 015-25191 | Wako | pSyn #64 | RRID:AB_2537218 | Mouse | 1.0 mg/mL | 1:1000 | 1:2000 | 1:1000 | 1:1000 | residues 124-134 |
| anti-**pS129**-aSyn | ab168381 | Abcam | MJF-R13 | RRID:AB_2728613 | Rabbit | 4.229 mg/mL | 1:3000 | 1:3000 | 1:1000 | 1:2000 | The exact sequence is proprietary |
| anti-**pS129**-aSyn | ab51253 | Abcam | EP1536Y | RRID:AB_869973 | Rabbit | 2.578 mg/mL | 1:500 | 1:500* | 1:500* | 1:1000 | Synthetic peptide within human aSyn (aa 100 to the C-terminus). The exact sequence is proprietary. |
| anti-**pS129**-aSyn | GTX82738 | GeneTex | - | AB_11176838 | Rabbit | 1 mg/mL | 1:500 | 1:1000 | 1:1000 | 1:1000 | Synthetic phospho-peptide corresponding to amino acid residues surrounding Ser129 conjugated to KLH |
| anti-**pS129**-aSyn | Lashuel | - | LASH-EGT-pS129 | - | Rabbit | 0.1 mg/mL | 1:500 | 1:500 | 1:500 | 1:500 | A-Y-E-M-P-**SP**-E-E-G-Y-Q-D |
| anti-**pS129**-aSyn | ab59264 | Abcam | - | AB_2270761 | Rabbit | 1 mg/mL | 1:500 | 1:500 | 1:500 | 1:1000 | Synthetic phosphopeptide derived from human aSyn around the S129 residue (M-P-**S**P-E-E). |
| anti-**pY125**-aSyn | 558246 | BD | - | AB_647301 | Mouse | 0.1 mg/mL | 1:1000 | - | - | - | P-D-N-E-A-**Y**P-E-M-P-S-E-E-G |
| anti-**pY125**-aSyn | Lashuel | - | LASH-EGT-pY125 | | Rabbit | 0.1 mg/mL | 1:500 | - | - | - | Phosphorylated peptide corresponding to the region including Y125 residue of aSyn |
| anti-**pY39**-aSyn | 849202 | Biolegend | A15119B | AB_2650702 | Mouse | 0.5 mg/mL | 1:1000 | - | - | - | KLH-conjugated α-synuclein synthetic peptide containing pY39 |
| anti-**pS87**-aSyn | Lashuel | - | 128005 | - | Rabbit | 0.1 mg/mL | 1:1000 | - | - | - | H2N- C A Q K T V E G A G S(p) I A A -CONHH |

* Due to lot-to-lot variability between 2019 and 2020, the ICC application was removed by Abcam from the EP1536Y antibody datasheet

**C**

| Primary Antibody | Catalog # | Company | Clone | RRID | Host | Concentration | WB dilution | ICC dilution |
|---|---|---|---|---|---|---|---|---|
| anti-**actin** | ab6276 | Abcam | AC-15 | RRID:AB_2223210 | Mouse | 2.2 mg/mL | 1:5000 | - |
| anti-**MAP2** | ab92434 | Abcam | - | RRID:AB_2138147 | Chicken | Not provided | Not tested | 1:2000 |
| anti-**a-Tubulin** | T6199 | Sigma | DM1A | RRID:AB_477583 | Mouse | Not provided | 1:5000 | - |
| anti-**Lamin B1** | ab133741 | Abcam | EPR8985 | RRID:AB_2616597 | Rab Mab | 0.33 mg/mL | 1:2000 | - |
| anti-**E-Cadherin** | #3195 | Cell Signaling | 24E10 | RRID:AB_2291471 | Rab Mab | Not provided | 1:1000 | - |

**D**

| Secondary Antibody | Catalog # | Company | RRID | Concentration | WB dilution | ICC dilution |
|---|---|---|---|---|---|---|
| Goat anti-**mouse Alexa Fluor 680** | A21058 | Invitrogen | RRID:AB_2535724 | 2 mg/mL | 1:5000 | - |
| Goat anti-**rabbit Alexa Fluor 680** | A21109 | Invitrogen | RRID:AB_2535758 | 2 mg/mL | 1:5000 | - |
| Goat anti-**mouse Alexa Fluor 800** | 926-32210 | Li-Cor | RRID:AB_621842 | 1 mg/mL | 1:5000 | - |
| Goat anti-**rabbit Alexa Fluor 800** | 926-32211 | Li-Cor | RRID:AB_621843 | 1 mg/mL | 1:5000 | - |
| Donkey anti-**rabbit Alexa Fluor 647** | A31573 | Invitrogen | RRID:AB_2536183 | 2 mg/mL | - | 1:800 |
| Donkey anti-**mouse Alexa Fluor 647** | A31571 | Invitrogen | RRID:AB_162542 | 2 mg/mL | - | 1:800 |
| Goat anti-**chicken Alexa Fluor 568** | A11041 | Invitrogen | RRID:AB_2534098 | 2 mg/mL | - | 1:500 |
| Goat anti-**mouse Alexa Fluor 488** | A-11029 | Invitrogen | RRID:AB_2534088 | 2 mg/mL | - | 1:800 |
| Donkey anti-**chicken Alexa Fluor 488** | 703-545-155 | Jackson Immunoresearch | RRID:AB_2340375 | 1 mg/mL | - | 1:400 |
| Donkey anti-**rabbit Alexa Fluor 488** | A21206 | Invitrogen | RRID:AB_2535792 | 2 mg/mL | - | 1:800 |

**Figure S3. List of the antibodies used in this study (related to Figures 1 to 9)**

**A.** Antibodies used for the detection of total aSyn.

**B.** Antibodies used for the detection of α-syn phosphorylated at residue S129.

**C.** Other antibodies used in the study.

**D.** Secondary antibodies used for immunoblotting or confocal imaging.

#### Figure S4

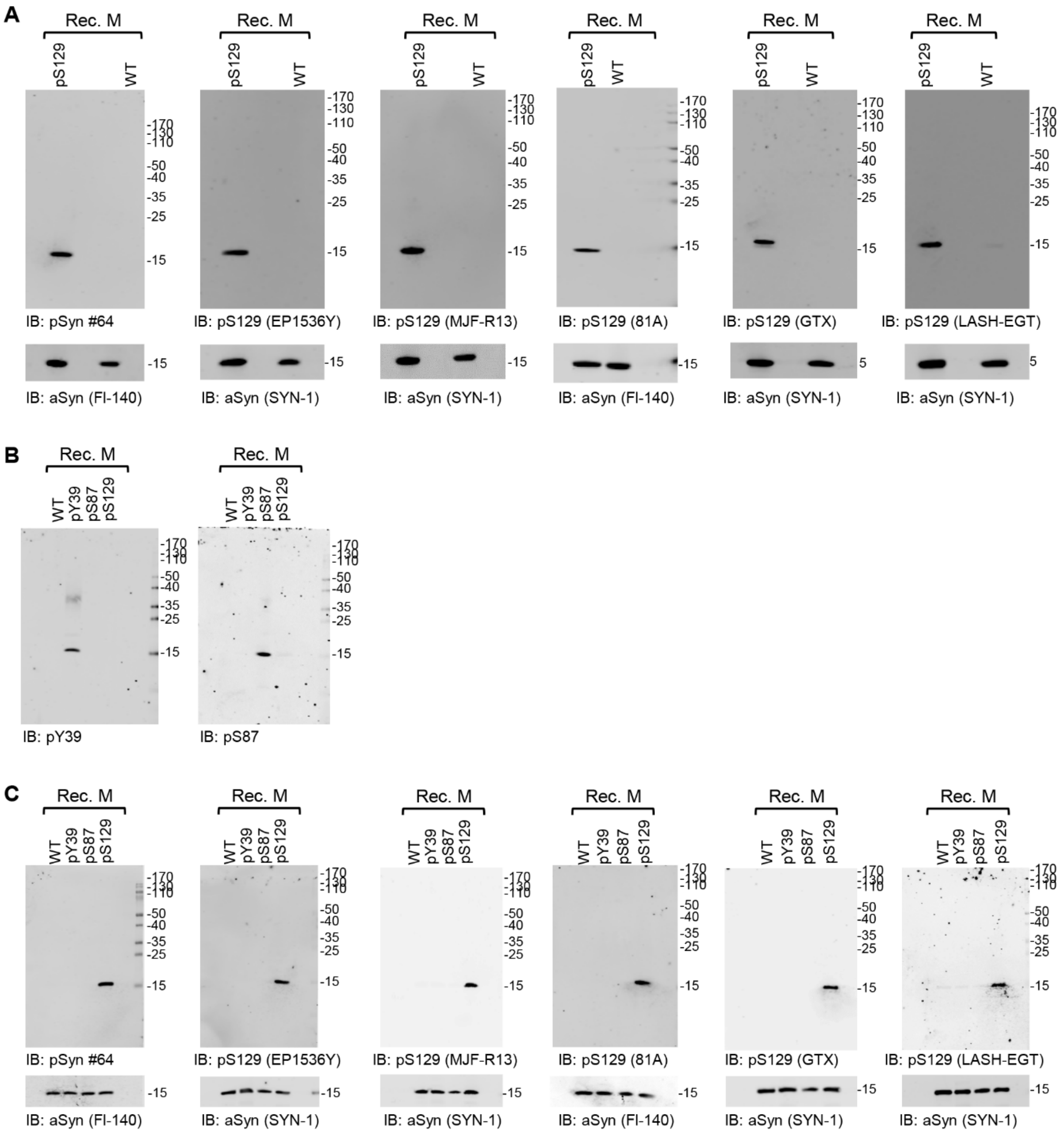

**Figure S4. The selected pS129 antibodies recognize all specifically pS129-aSyn but not unmodified aSyn** (related to Figure 2)

**A.** Forty nanograms of human recombinant monomeric aSyn unphosphorylated (WT) or phosphorylated at the S129 residue (pS129) were detected by WB using commercial antibodies (pSyn#64, MJF-R13, 81a, EP1536Y and GTX) or the LASH-EGT homemade pS129 antibody (respective epitopes listed in Figure S3) combined with total aSyn antibodies (Fl-140 or SYN-1).

**B-C.** Forty nanograms of human recombinant monomeric aSyn (WT) or phosphorylated at the Y39 (pY39) or S87 (pS87) or S129 residues (pS129) and WB was performed using the pY39 antibody (**B**) or the LASH-EGT homemade pS87 antibody (**B**) (respective epitopes listed in Figure S3) or the pS129 antibodies (**C**) used in **A** combined with total aSyn antibodies (FI-140 or SYN-1, **C**).

#### Figure S5

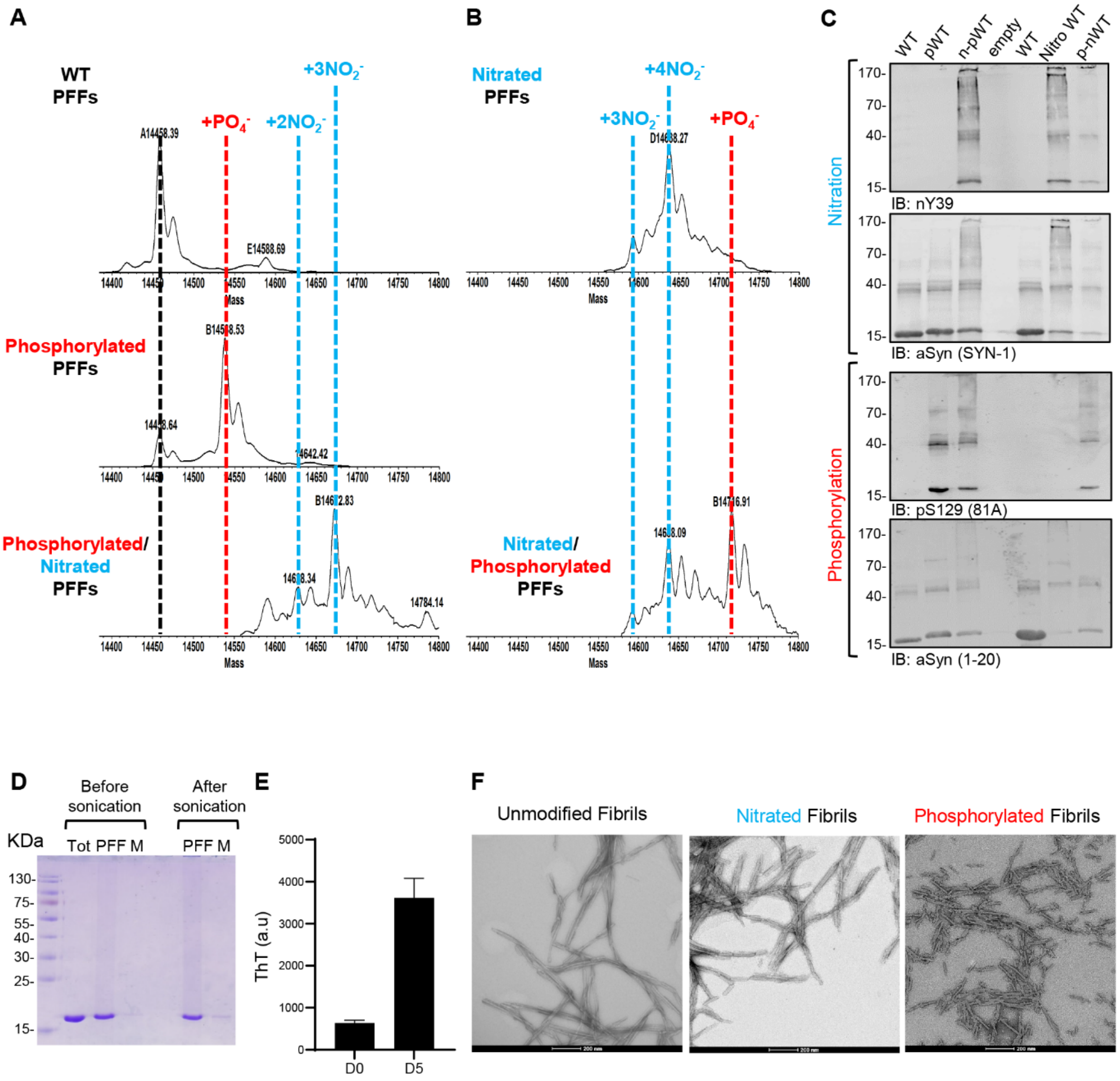

**Figure S5. Preparation and characterization of recombinant nitrated and/or phosphorylated monomeric and fibrillar aSyn species (related to Figure 3)**

**A.** WT aSyn fibrils were incubated for 18 h at 30°C with the phosphorylating enzyme PLK3. Afterwards, the phosphorylated fibrils were incubated for 2 h with the nitric oxide (NO) donor TNM. **B.** WT aSyn fibrils were incubated for 2 h with TNM and afterwards incubated for 18 h at 30°C with PLK3. The product reactions were analyzed by ESI-LC/MS. **C.** WB analyses were performed on the samples from **A** and **B**. The presence of the different types of aSyn fibrils was detected by the total aSyn antibody (SYN-1 or 1-20). As expected, the nY39 antibody only detected the nitrated PFFs, while the pS129 antibody (81A) specifically detected the phosphorylated PFFs.

**D-F.** Characterization of the aSyn PFFs. **D.** The amount of monomer that did not fibrillize or that was released after sonication by SDS-PAGE gel and Coomassie blue staining. **E.** Fibril formation was also confirmed by ThT fluorometry after sonication. All data represent the average ± SD (n=3). **F.** After sonication, aSyn WT, nitrated and/or phosphorylated PFFs were analyzed by transmission electron microscopy (TEM) imaging. Scale bars = 200 nm.

#### Figure S6

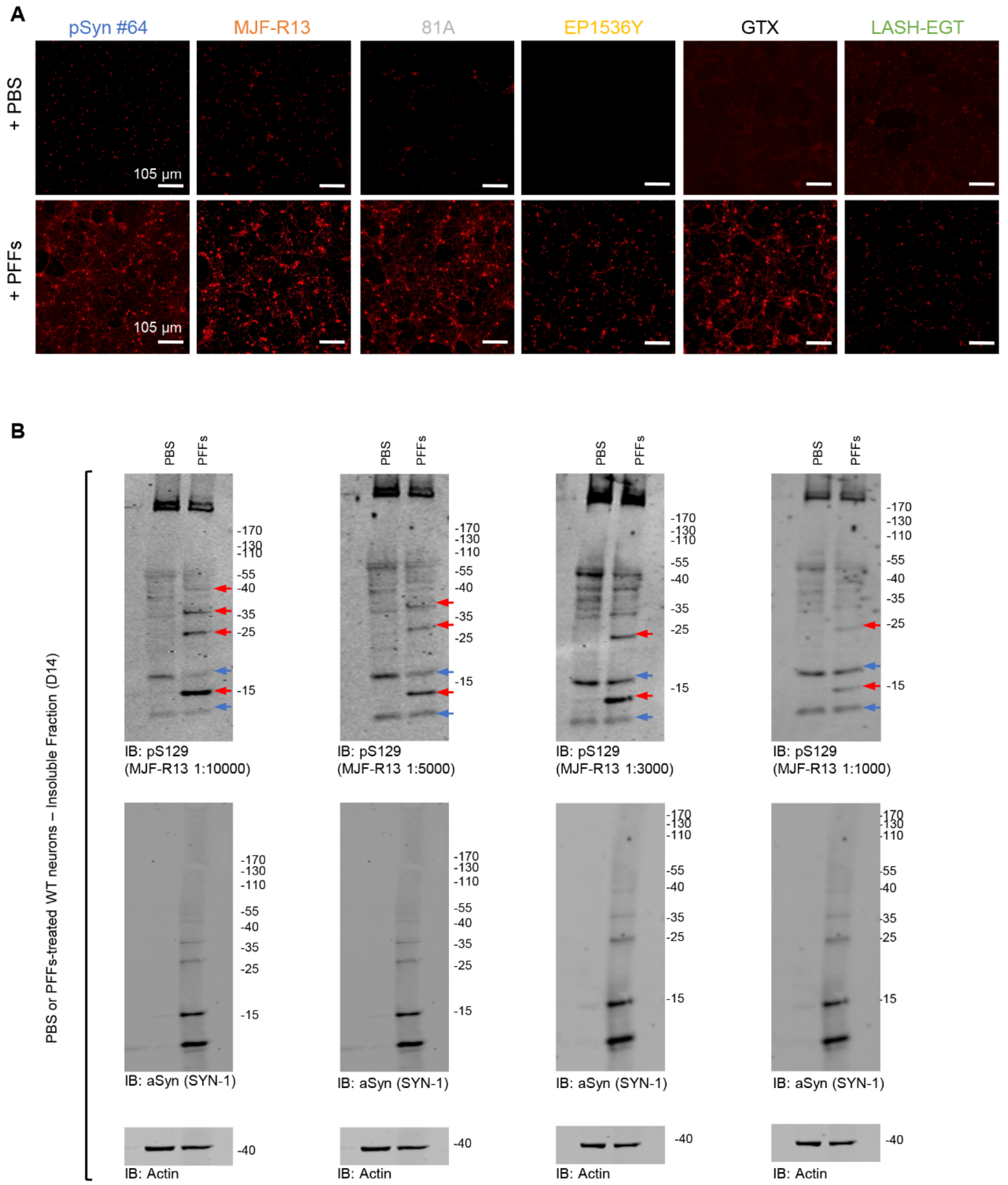

**Figure S6. Assessment of the pS129 antibodies detection in the PFFs-seeded WT primary hippocampal neurons (related to Figure 4)**

**A.** After 14 days of treatment with 70 nM of mouse aSyn PFFs or PBS buffer (negative control), primary hippocampal neurons were fixed, and ICC was performed. Newly formed fibrils were detected using the commercial antibodies (pSyn#64, MJF-R13, 81A, EP1536Y and GTX) or the LASH-EGT homemade pS129 antibody. Neurons were counterstained with MAP2 antibody and the nucleus with DAPI staining. Images were acquired by a high-throughput wide-field cell imaging system (In Cell Analyser 2200), and pS129 intensity in MAP2-positive neurons was quantified (see quantification in Figure 4B-C). Scale bar = 105 µM. **B.** Optimization of the dilution of the MJF-R13 pS129 antibody on the insoluble fractions (SDS-soluble fraction) of primary hippocampal neurons treated with PBS or with 70 nM of mouse aSyn PFFs for 14 days (D14). The optimal concentration of the MJF-R13 antibody is at 1/10 000 for the lot

GR3227218 (Abcam). The red arrows indicate the pS129-aSyn positive bands. The blue arrows indicate the non-specific bands.

**Figure S7**

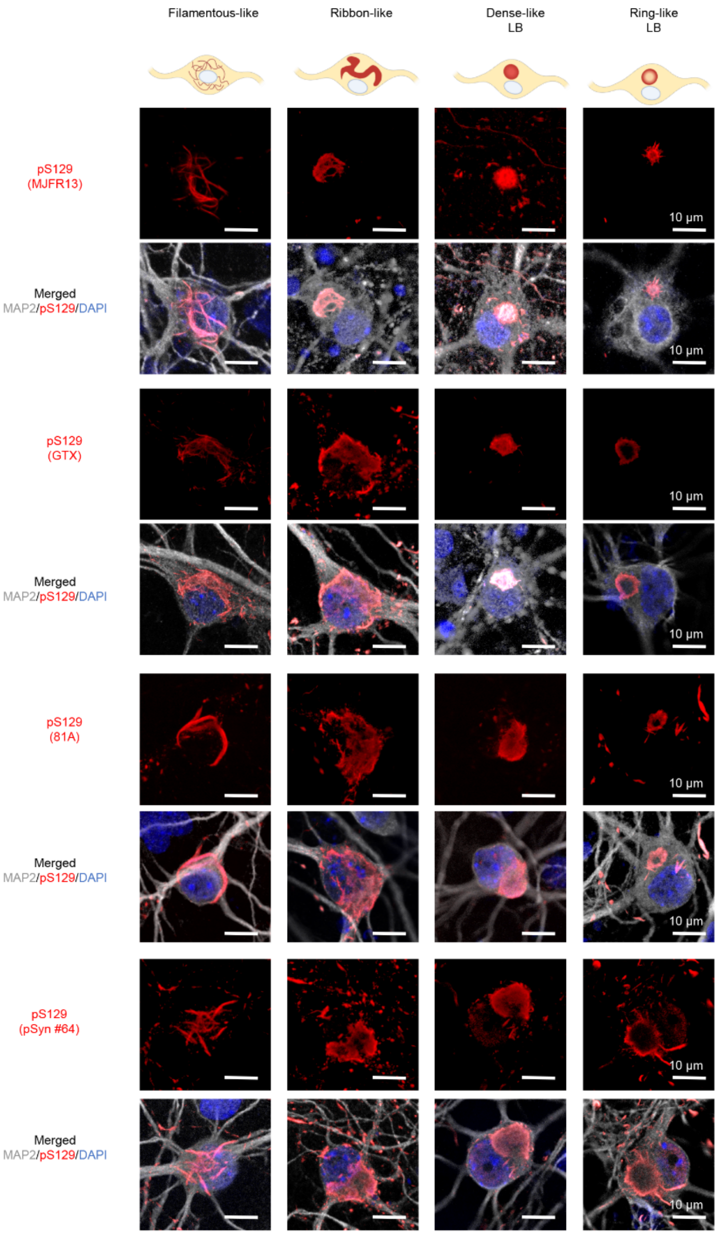

**Figure S7. All the pS129 antibodies detect the same types of seeded-aggregates in mouse PFFs-treated cortical neurons (related to Figure 6)**

After 21 days of treatment with 70 nM of mouse aSyn PFFs, primary cortical neurons were fixed, and ICC was performed. The seeded-aggregates were detected using the pS129 antibodies (MJF-R13, GTX, 81A or pSyn#64). Neurons were counterstained with the MAP2 antibody and the nucleus with DAPI staining. Scale bar = 10 μM. All four pS129 antibodies were able to distinguish the seeded-aggregates with the filamentous or the ribbon-like morphology and the LB-like inclusions with the dense core and the ring-like morphology. The schematic representation of the different types of seeded-aggregates was created using BioRender.com.

**Figure S8**

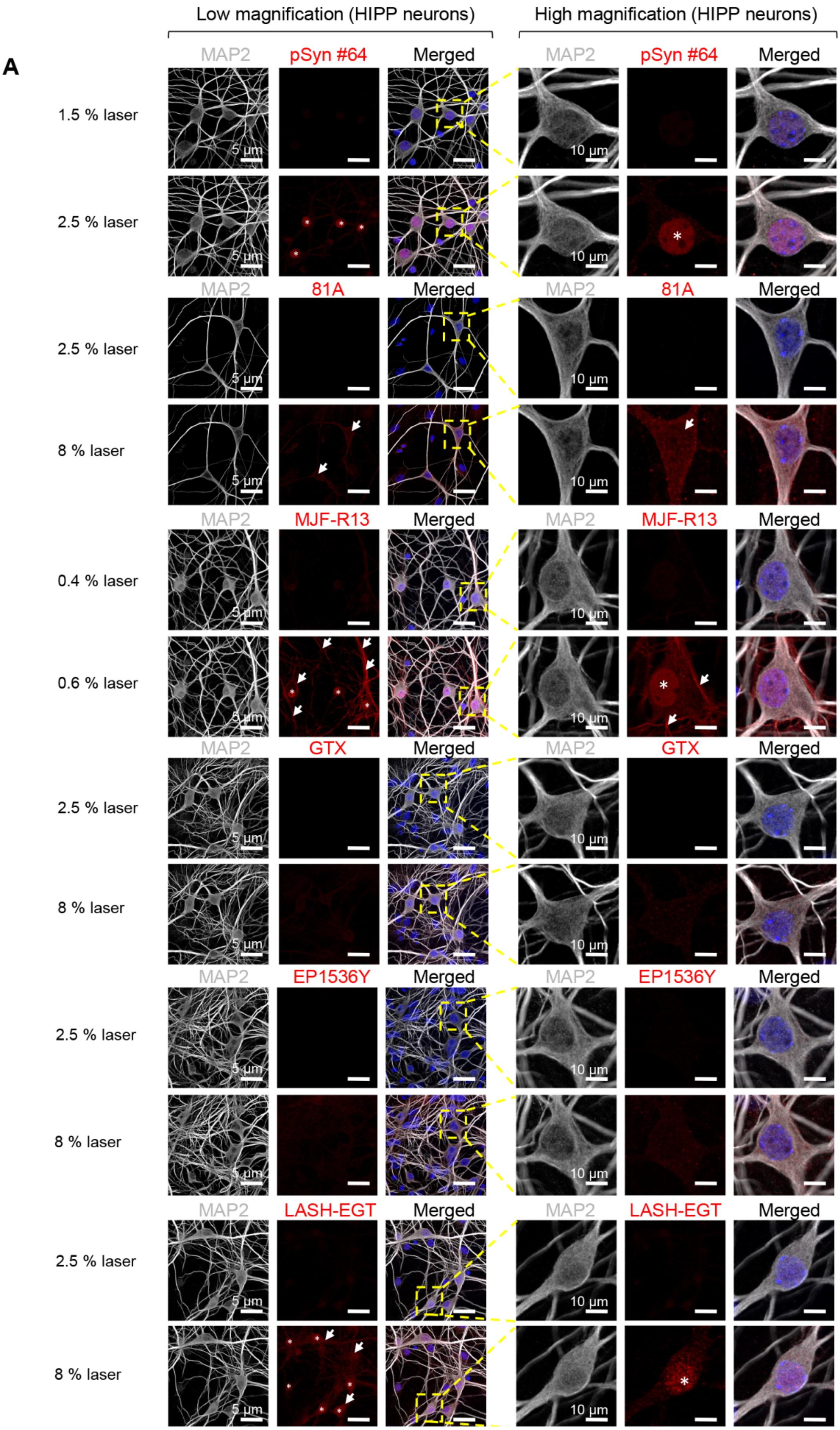
Low magnification (HIPP neurons)
High magnification (HIPP neurons)
A
MAP2
pSyn #64
Merged
1.5 % laser
2.5 % laser
81A
8 % laser
MJF-R13
0.4 % laser
0.6 % laser
GTX
EP1536Y
LASH-EGT
5 µm
10 µm

**Figure S8**

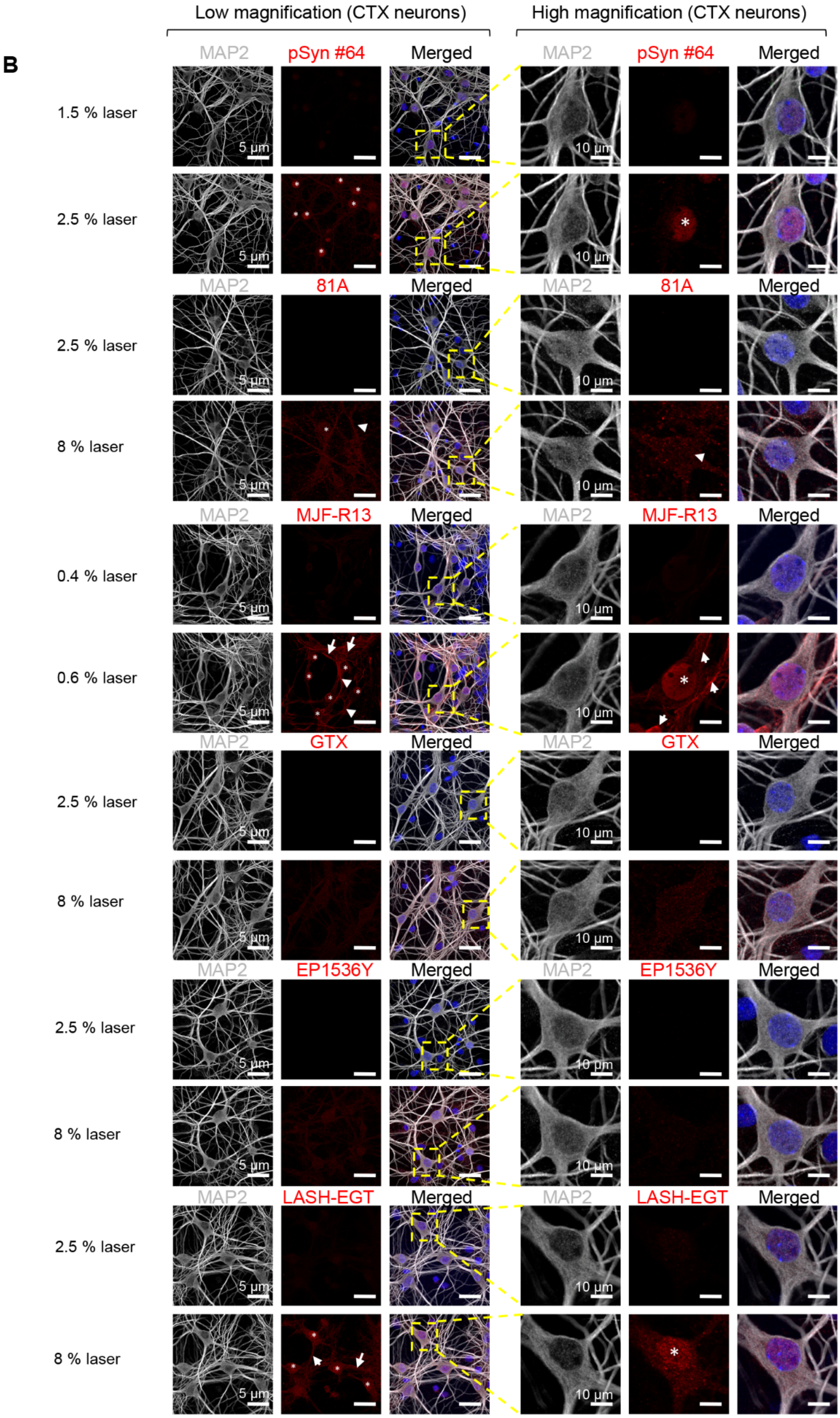
Low magnification (CTX neurons)
High magnification (CTX neurons)
B
MAP2
pSyn #64
Merged
1.5 % laser
2.5 % laser
81A
2.5 % laser
8 % laser
MJF-R13
0.4 % laser
0.6 % laser
GTX
2.5 % laser
8 % laser
EP1536Y
2.5 % laser
8 % laser
LASH-EGT
2.5 % laser
8 % laser
5 μm
10 μm

**Figure S8**

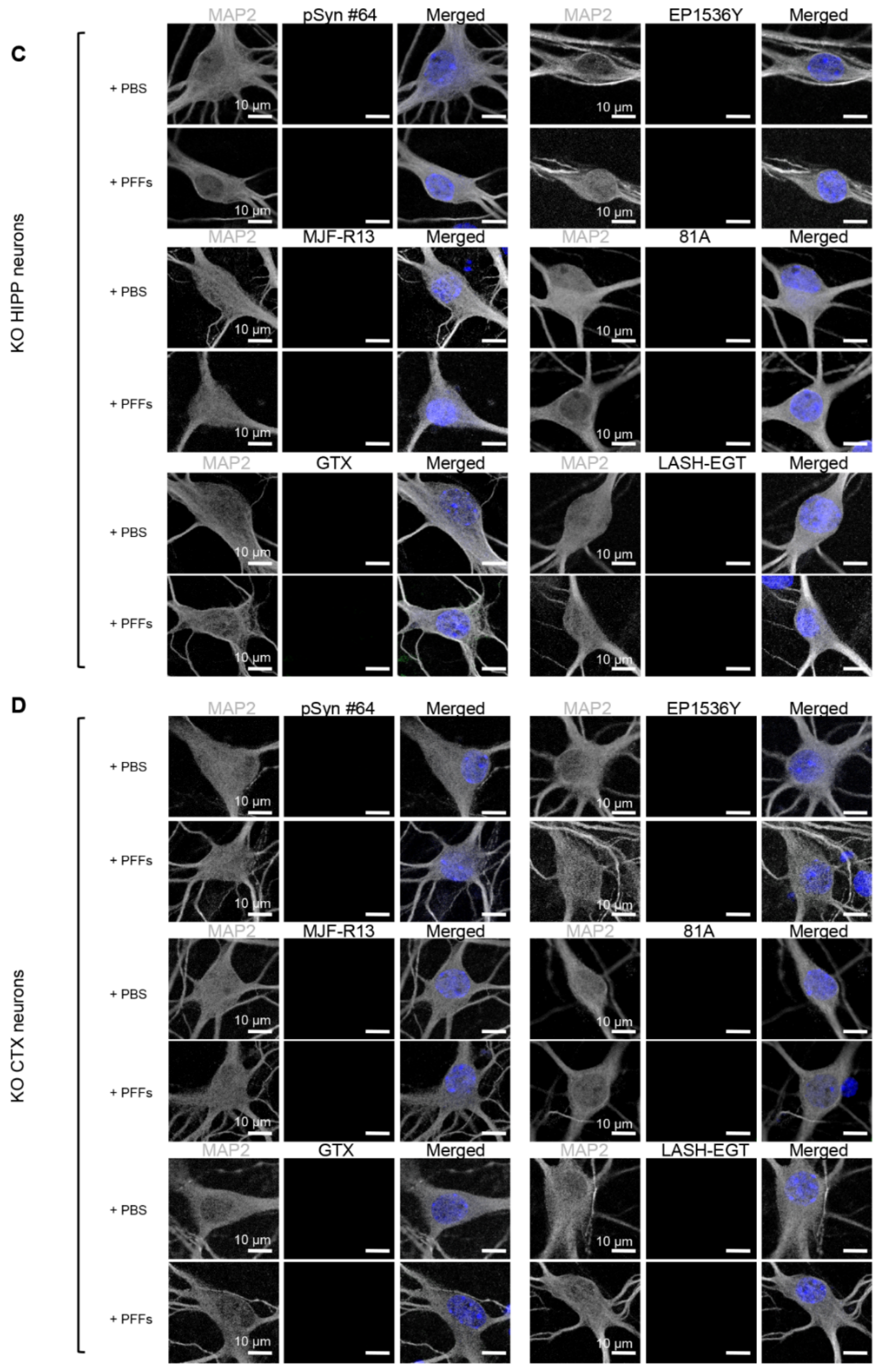

**Figure S8. Assessment of the pS129 antibodies background and cross-reactivity in aSyn KO primary neurons by ICC and confocal imaging** (related to Figures 4-6)

**A-B.** At DIV14, hippocampal (**A**) or cortical (**B**) aSyn KO primary neurons were fixed, and ICC was performed using the six pS129 antibodies (pSyn#64, MJF-R13, 81A, EP1536Y, GTX and LASH-EGT). Neurons were counterstained with the MAP2 antibody and the nucleus with DAPI staining. Confocal imaging was performed using different laser powers to reveal the background and the potential cross-reactivity of pS129 antibodies in the nucleus (white star), cytosol and neuritic extensions (white arrows) of neurons lacking aSyn expression. Scale bars = 5 and 10 μM. **C-D.** At DIV14, hippocampal (**C**) or cortical (**D**) aSyn KO primary neurons were treated for 24 h with 70 nM of mouse aSyn PFFs before being fixed. ICC was performed using the six pS129 antibodies (pSyn#64, MJF-R13, 81A, EP1536Y, GTX and LASH-EGT). Neurons were counterstained with the MAP2 antibody and the nucleus with DAPI staining. Confocal imaging was performed using the same laser intensity and the PMT gain and offset used when imaging WT neurons containing pS129 positive aggregates (Figure 4). This helped eliminate the unspecific signal initially observed in the aSyn KO primary neurons shown in **A** and **B** panels.

**Figure S9**

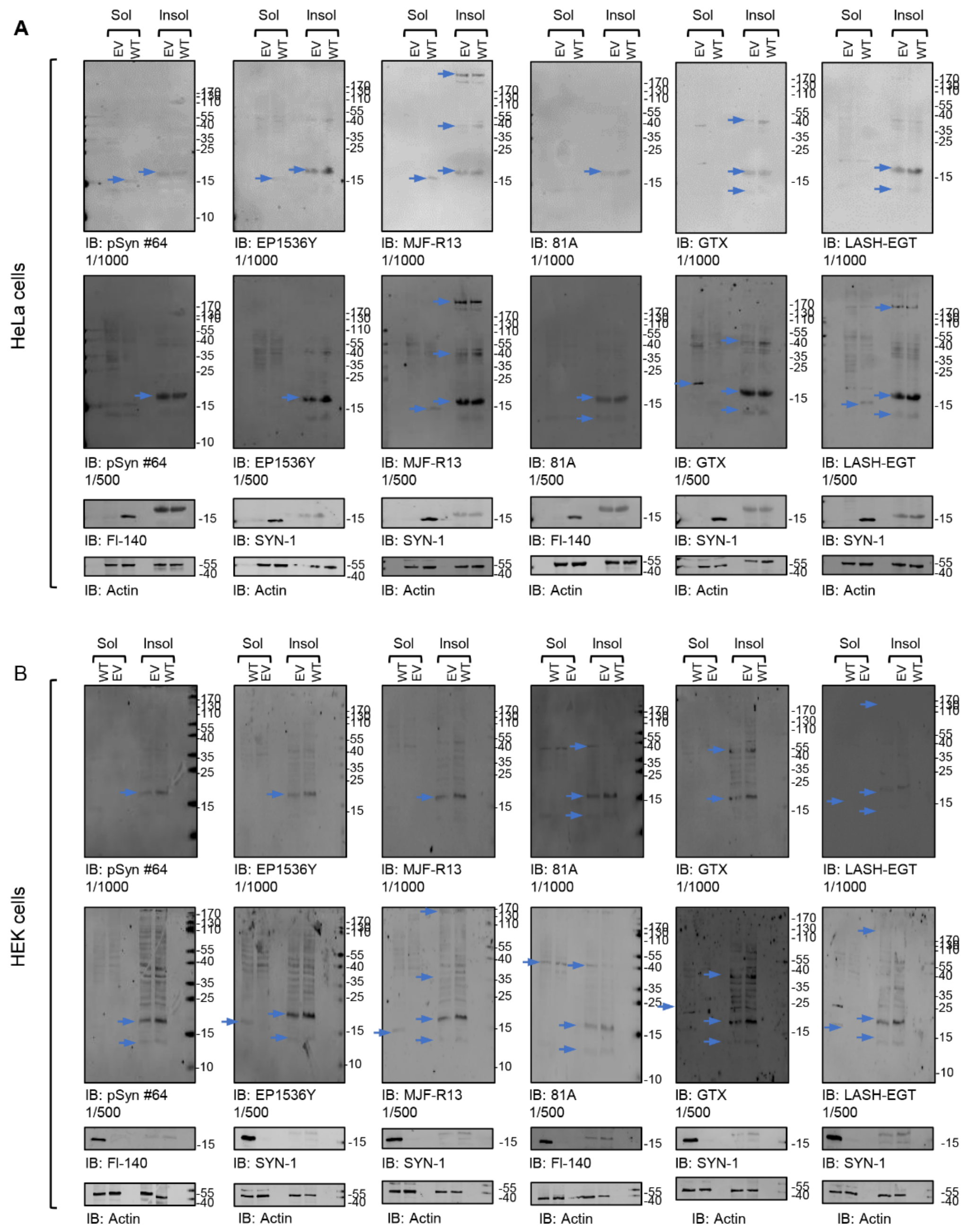

**Figure S9. Assessment of the pS129 antibodies detection in the soluble and insoluble fractions of Hela and HEK293 mammalian cell lines by WB (related to Figures 5 and 7)**

HeLa and HEK293 cells were transfected with WT aSyn (WT) or empty (EV) vectors. After 48 h, the transfected cells were lysed and sequential biochemical extraction. pS129 level of detection in the soluble (0.1% Triton-soluble fraction, sol) and the insoluble fraction (SDS-soluble fraction, insol) were extracted from the HeLa (**A**) or HEK293 cells (**B**) using the six pS129 antibodies (pSyn#64, MJF-R13, 81A, EP1536Y, GTX and LASH-EGT) at a dilution of 1/1000. Forty nanograms of recombinant aSyn WT or phosphorylated at residue S129 (pS129) were used as positive controls. Membranes were counterstained by total aSyn antibodies (SYN-1 or FI-140). Actin was used as a loading control. The red arrows indicate the pS129-aSyn positive bands. The blue arrows indicate the non-specific bands.

**Figure S10**

**A HeLa cells**

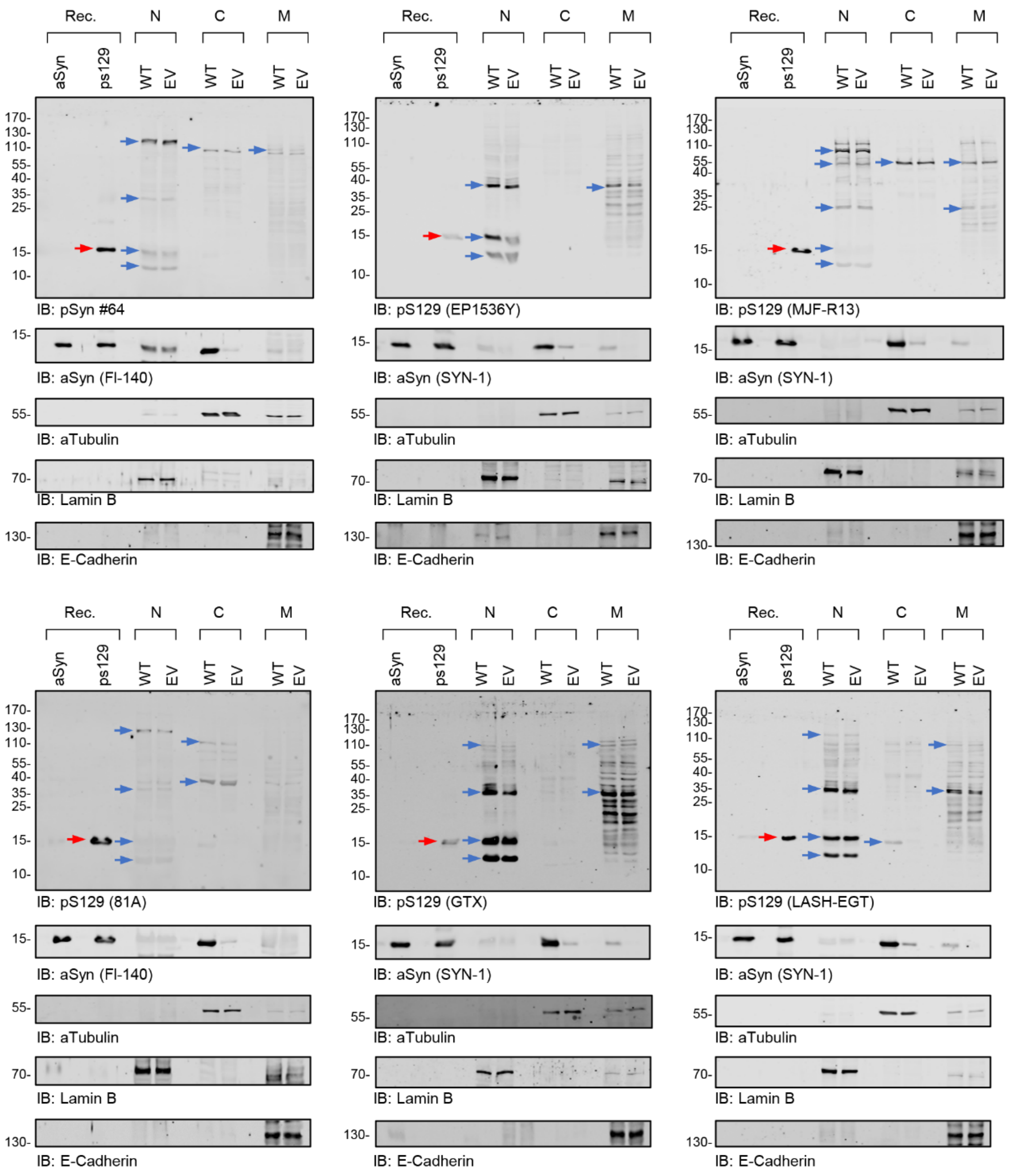

**Figure S10**

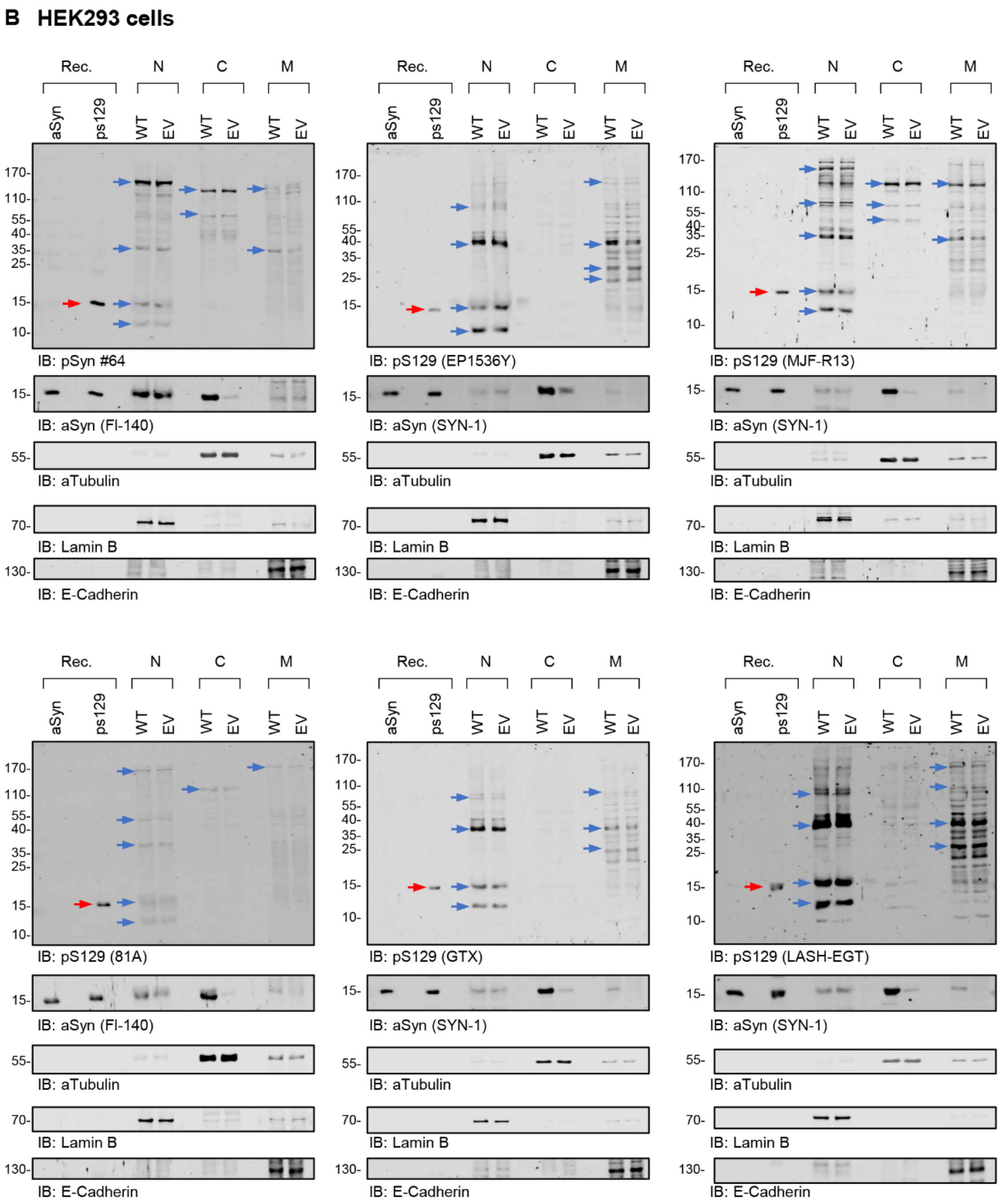

**Figure S10. Assessment of the pS129 antibodies detection in the nuclear, cytosolic and membranous fractions of Hela and HEK293 mammalian cell lines by WB (related to Figures 5 and 7)**

HeLa and HEK293 cells were transfected with WT aSyn or empty (EV) vectors. After 48 h, the transfected cells were lysed and sequential biochemical extraction. pS129 level of detection in the nuclear (N), cytosolic (C), and membrane (M) fractions (A-B) or total cell (C) lysates extracted from the HeLa (**A**) or HEK293 cells (**B**) using the six pS129 antibodies (pSyn#64, MJF-R13, 81A, EP1536Y, GTX and LASH-EGT) at a dilution of 1/1000. Forty nanograms of recombinant aSyn WT or phosphorylated at residue S129 (pS129, indicated by a red arrow) were used as positive controls. Membranes were counterstained by total aSyn antibodies (SYN-1 or FI-140). aTubulin, Laminin B and E-Cadherin were used as loading control for respectively the cytosolic, nuclear and membrane fractions. The red arrows indicate the pS129-aSyn positive bands. The blue arrows indicate the non-specific bands.

**Figure S11**

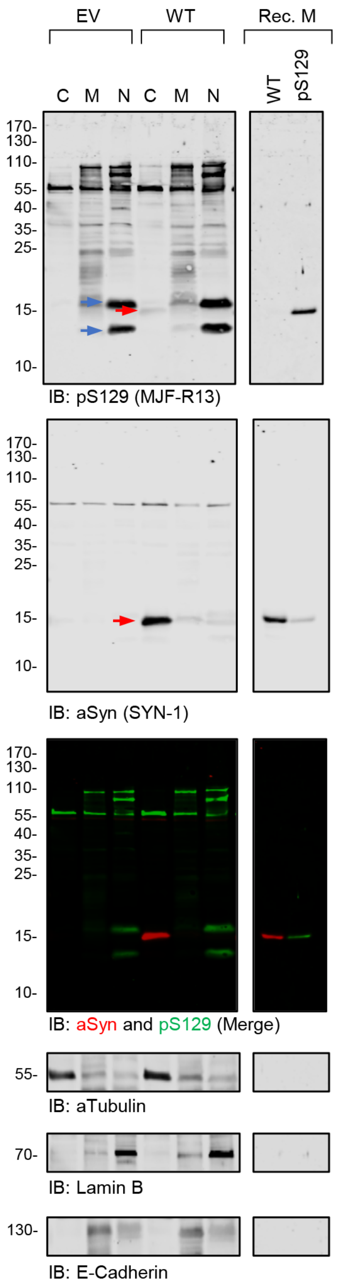

**Figure S11. The commercial tricine gel allows resolving the aSyn band from the non-specific bands (related to Figures 4, 5 and 7)**

HeLa cells were transfected with WT aSyn or empty (EV) vectors. After 48 h, the transfected cells were lysed, subcellular fractionation was performed, and proteins from the nuclear (N), cytosolic (C) and membrane (M) fractions were separated using commercial 16% tricine gel (Invitrogen). Forty nanograms of recombinant aSyn WT or phosphorylated at residue S129 (pS129) were used as positive controls. Membranes were co-stained with the MJF-R13 (pS129) and SYN-1 (total aSyn). Actin was used as a loading control. The red arrows indicate the pS129-aSyn positive bands. The blue arrows indicate the non-specific bands.

**Figure S12**

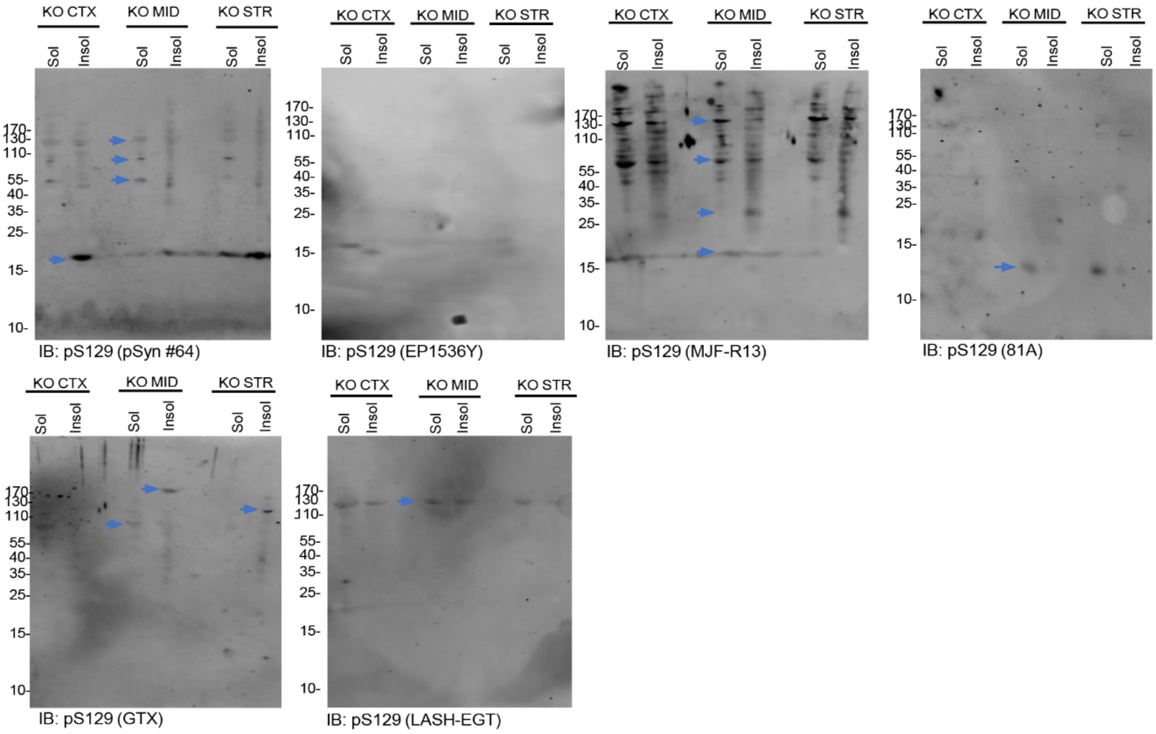

**Figure S12. Assessment of the pS129 antibodies detection in brain sections of aSyn KO mice (related to Figure 7B)**

pS129 level of detection in the soluble and insoluble fractions extracted from the aSyn KO mice brains using the six pS129 antibodies (pSyn#64, MJF-R13, 81a, EP1536Y, GTX and LASH-EGT). Sequential biochemical extraction was performed on different brain regions (cortical/CTX; Midbrain/MID and Striatum/STR) derived from aSyn KO mice, and the level of pS129 was assessed in the soluble (0.1% Triton-soluble fraction) and the insoluble fraction (SDS-soluble fraction). Membranes were counterstained for total aSyn (SYN-1 or D37A6), and actin was used as a loading control. The red arrows indicate the pS129-aSyn positive bands. The blue arrows indicate the non-specific bands. Membranes shown in Figure S12 are the same as in Figure 7B but acquired at a higher exposure.

#### Figure S13

A

| | Total | S129 | S87 | Y39 | Y125 | Y133 | Y136 |
|---|---|---|---|---|---|---|---|
| **Previously detected in plasma** | 57 | 27 | 13 | 2 | 4 | 2 | 11 |
| **Previously detected in CSF** | 38 | 19 | 10 | 1 | 3 | 0 | 7 |
| **Combined Proteins identified** | 121 | 59 | 32 | 4 | 5 | 3 | 20 |

##### B S129 human – part 1

| Peptide alignment | Position | Length | MW list [kDa] | Exact | Similar | Mismatch | Max exact stretch | Max exact + similar stretch | ID | Also in mouse | Description | Subcellular location | Detected in plasma | Detected in CSF |
|---|---|---|---|---|---|---|---|---|---|---|---|---|---|---|
| EAYEMPSEEGYQD<br>::::::::::::<br>EAYEMPSEEGYQD | 123 | 140 | 14 | 13 | 0 | 0 | 13 | 13 | SYUA_HUMAN | Y | Alpha-synuclein | Cell junction. Cytoplasm. Membrane. Nucleus. Secreted | Y | Y |
| EKVELPSEEQVSG<br>: :.::::<br>EAYEMPSEEGYQD | 375 | 1782 | 190 | 6 | 1 | 6 | 4 | 6 | AKA12_HUMAN | Y | A-kinase anchor protein 12 | Cytoplasm. Membrane | Y | Y |
| SFPELPSEE****<br> :.::::<br>EAYEMPSEEGYQD | 920 | 928 | 100 | 5 | 1 | 7 | 4 | 6 | NIBA1_HUMAN | Y | Protein Niban 1 | Cytoplasm. Membrane | Y | Y |
| EEDDYPSEELLED<br>: . :::: .:<br>EAYEMPSEEGYQD | 721 | 1907 | 210 | 6 | 2 | 5 | 4 | 4 | TGO1_HUMAN | N | Transport and Golgi organization protein 1 homolog | Endoplasmic reticulum membrane | Y | Y |
| EARKKPSEEEAAV<br>:: ::::<br>EAYEMPSEEGYQD | 162 | 195 | 21 | 6 | 0 | 7 | 4 | 4 | CY24A_HUMAN | Y | Cytochrome b-245 light chain | Cell membrane | N | N |
| KARENPSEEAQNL<br> : : ::::<br>EAYEMPSEEGYQD | 115 | 501 | 59 | 6 | 0 | 7 | 4 | 4 | SF3A3_HUMAN | Y | Splicing factor 3A subunit 3 | Nucleus. Nucleus speckle | Y | N |
| EDFRRPSEEDFRR<br>: . :::: .<br>EAYEMPSEEGYQD | 585 | 1001 | 120 | 5 | 2 | 6 | 4 | 4 | RB12B_HUMAN | N | RNA-binding protein 12B | unspecified. | N | N |
| KEELVPSEEDFQG<br> :::: .:<br>EAYEMPSEEGYQD | 579 | 1001 | 110 | 5 | 1 | 7 | 4 | 4 | BRE1B_HUMAN | N | E3 ubiquitin-protein ligase BRE1B | Nucleus | N | Y |
| PARTPPSEEDSAE<br> : :::: .<br>EAYEMPSEEGYQD | 78 | 313 | 34 | 5 | 1 | 7 | 4 | 4 | SGTA_HUMAN | Y | Small glutamine-rich tetratricopeptide repeat-containing protein alpha | Cytoplasm. Nucleus | Y | N |
| VGVEEPSEEDQNE<br> : :::: .<br>EAYEMPSEEGYQD | 539 | 725 | 80 | 5 | 1 | 7 | 4 | 4 | SL9A7_HUMAN | N | Sodium/hydrogen exchanger 7 | Cell membrane. Golgi apparatus. Recycling endosome membrane | N | N |
| LPEVPPSEEEEQE<br> :::: :.<br>EAYEMPSEEGYQD | 809 | 1127 | 130 | 5 | 1 | 7 | 4 | 4 | TEX2_HUMAN | Y | Testis-expressed protein 2 | Endoplasmic reticulum membrane. Nucleus membrane | N | N |
| KTERKPSEEEYVI<br> :::: :<br>EAYEMPSEEGYQD | 634 | 776 | 88 | 5 | 0 | 8 | 4 | 4 | ARHG6_HUMAN | N | Rho guanine nucleotide exchange factor 6 | Cell projection | N | N |
| TKNEEPSEEEIDA<br> : ::::<br>EAYEMPSEEGYQD | 115 | 783 | 87 | 5 | 0 | 8 | 4 | 4 | DDX21_HUMAN | Y | Nucleolar RNA helicase 2 | Cytoplasm. Mitochondrion. Nucleus | Y | N |
| PRESRPSEEREWD<br> :::: :<br>EAYEMPSEEGYQD | 1192 | 1382 | 170 | 5 | 0 | 8 | 4 | 4 | EIF3A_HUMAN | Y | Eukaryotic translation initiation factor 3 subunit A | Cytoplasm | Y | N |
| PPAGRPSEEPEPD<br> :::: :<br>EAYEMPSEEGYQD | 374 | 432 | 47 | 5 | 0 | 8 | 4 | 4 | PHAG1_HUMAN | Y | Phosphoprotein associated with glycosphingolipid-enriched microdomains 1 | Cell membrane | Y * | Y |
| KIQRMPSESAAQS<br> :::: :<br>EAYEMPSEEGYQD | 113 | 514 | 57 | 5 | 0 | 8 | 4 | 4 | PLRG1_HUMAN | Y | Pleiotropic regulator 1 | Nucleus. Nucleus speckle | N | N |
| RGEDVPSEEEEEE<br> . :::: ..<br>EAYEMPSEEGYQD | 178 | 412 | 47 | 4 | 3 | 6 | 4 | 4 | NOB1_HUMAN | N | RNA-binding protein NOB1 | Nucleus | N | N |
| SGSQGPSEEKPAP<br> . ::::<br>EAYEMPSEEGYQD | 386 | 1782 | 190 | 4 | 1 | 8 | 4 | 4 | AKA12_HUMAN | Y | A-kinase anchor protein 12 | Cytoplasm. Membrane | Y | Y |
| DIIRQPSEEEIIK<br>. ::::<br>EAYEMPSEEGYQD | 110 | 130 | 15 | 4 | 1 | 8 | 4 | 4 | PEA15_HUMAN | Y | Astrocytic phosphoprotein PEA-15 | Cytoplasm | Y | Y |

### Figure S13

#### B S129 human – part 2

| Peptide alignment | Position | Length | MW list [kDa] | Exact | Similar | Mismatch | Max exact stretch | Max exact + similar stretch | ID | Also in mouse | Description | Subcellular location | Detected in plasma | Detected in CSF |
|---|---|---|---|---|---|---|---|---|---|---|---|---|---|---|
| TRKRGPSEETGGA<br> ::::<br>EAYEMPSEEGYQD | 571 | 726 | 81 | 4 | 0 | 9 | 4 | 4 | CCNT1_HUMAN | N | Cyclin-T1 | Nucleus | N | N |
| FHLMAPSEEDHSI<br> ::::<br>EAYEMPSEEGYQD | 138 | 259 | 28 | 4 | 0 | 9 | 4 | 4 | IBP1_HUMAN | N | Insulin-like growth factor-binding protein 1 | Secreted | Y | Y |
| LCTKAPSEEDSLS<br> ::::<br>EAYEMPSEEGYQD | 32 | 258 | 29 | 4 | 0 | 9 | 4 | 4 | RT18B_HUMAN | N | 28S ribosomal protein S18b, mitochondrial | Mitochondrion | N | N |
| KELPPPSEEIKTG<br> ::::<br>EAYEMPSEEGYQD | 531 | 657 | 76 | 4 | 0 | 9 | 4 | 4 | THOC1_HUMAN | Y | THO complex subunit 1 | Cytoplasm, Nucleus, Nucleus matrix and Nucleus speckle | N | N |
| AMLHLPSEQGAPE<br> .:::.: .<br>EAYEMPSEEGYQD | 37 | 419 | 48 | 4 | 3 | 6 | 3 | 6 | NEMO_HUMAN | Y | NF-kappa-B essential modulator | Cytoplasm and Nucleus | Y * | N |
| EETGSPSEDGMQS<br>: :::.: :<br>EAYEMPSEEGYQD | 53 | 501 | 56 | 6 | 1 | 6 | 3 | 5 | PWP1_HUMAN | N | Periodic tryptophan protein 1 homolog | Cytoplasm and Nucleus | N | N |
| MSRALPSEDEEGQ<br> .:::.<br>EAYEMPSEEGYQD | 8299 | 8797 | 1000 | 3 | 2 | 8 | 3 | 5 | SYNE1_HUMAN | Y | Nesprin-1 | Cytoplasm, Golgi apparatus, Nucleus, Nucleus envelope, Nucleus outer membrane | Y | Y # |
| EPEEGPSEDDKAE<br>: : :::. .<br>EAYEMPSEEGYQD | 4940 | 5596 | 630 | 5 | 2 | 6 | 3 | 4 | MDN1_HUMAN | N | Midasin | Cytoplasm and Nucleus | N | N |
| DSSPLPSEKSDED<br>. .::: .:<br>EAYEMPSEEGYQD | 1354 | 1710 | 200 | 4 | 3 | 6 | 3 | 4 | CHD1_HUMAN | Y | Chromodomain-helicase-DNA-binding protein 1 | Cytoplasm and Nucleus | N | N |
| ETQRKPSEDEVLN<br>: :::. .<br>EAYEMPSEEGYQD | 1052 | 1231 | 140 | 4 | 2 | 7 | 3 | 4 | EHBP1_HUMAN | Y | EH domain-binding protein 1 | Cytoplasm, Endosome, Membrane | N | N |
| YSVQEPSEDSEEE<br> . :::. ..<br>EAYEMPSEEGYQD | 1178 | 1374 | 140 | 3 | 4 | 6 | 3 | 4 | LMTK1_HUMAN | Y | Serine/threonine-protein kinase LMTK1 | Cytoplasm and Membrane | N | N |
| QYNQVPSEDFERT<br>. . :::.<br>EAYEMPSEEGYQD | 317 | 365 | 40 | 3 | 3 | 7 | 3 | 4 | CXAR_HUMAN | N | Coxsackievirus and adenovirus receptor | Basolateral cell membrane, Cell junction,<br>Cell membrane, Secreted | Y | Y |
| DSSRLPSEGPRPA<br>. .:::<br>EAYEMPSEEGYQD | 36 | 206 | 22 | 3 | 2 | 8 | 3 | 4 | BLVRB_HUMAN | Y | Flavin reductase (NADPH) | Cytoplasm | Y | Y |
| DQQSLPSEPEETL<br>. .:::<br>EAYEMPSEEGYQD | 341 | 543 | 58 | 3 | 2 | 8 | 3 | 4 | RETR2_HUMAN | N | Reticulophagy regulator 2 | Membrane | N | Y # |
| RTQTLPSENSEES<br> .::: .<br>EAYEMPSEEGYQD | 54 | 641 | 71 | 3 | 2 | 8 | 3 | 4 | RIOX1_HUMAN | N | Ribosomal oxygenase 1 | Nucleus | N | N |
| KHQSDPSEDEDER<br> :::. .<br>EAYEMPSEEGYQD | 123 | 496 | 57 | 3 | 2 | 8 | 3 | 4 | SGK3_HUMAN | Y | Serine/threonine-protein kinase Sgk3 | Cytoplasmic vesicle, Early endosome and<br>Recycling endosome | N | N |
| KSVSTPSEAGSQD<br> ::: : ::<br>EAYEMPSEEGYQD | 101 | 645 | 71;73 | 6 | 0 | 7 | 3 | 3 | DC1I1_HUMAN | N | Cytoplasmic dynein 1 intermediate chain 1 | Chromosome and Cytoplasm | N | N |
| EEEEEPSEGLEEE<br>: : ::: ..<br>EAYEMPSEEGYQD | 335 | 1035 | 110 | 5 | 2 | 6 | 3 | 3 | GGYF1_HUMAN | Y | GRB10-interacting GYF protein 1 | Unspecified | N | N |
| DLKESPSEGSLQP<br>. : ::: :<br>EAYEMPSEEGYQD | 2 | 512 | 58 | 5 | 1 | 7 | 3 | 3 | ASIC2_HUMAN | N | Acid-sensing ion channel 2 | Cell membrane | N | N |

### Figure S13

#### B S129 human – part 3

| Peptide alignment | Position | Length | MW list [kDa] | Exact | Similar | Mismatch | Max exact stretch | Max exact + similar stretch | ID | Also in mouse | Description | Subcellular location | Detected in plasma | Detected in CSF |
|---|---|---|---|---|---|---|---|---|---|---|---|---|---|---|
| EAASGPSESPSPP<br>:: :::<br>EAYEMPSEEGYQD | 75 | 814 | 92 | 5 | 0 | 8 | 3 | 3 | EIF3B_HUMAN | N | Eukaryotic translation initiation factor 3 subunit B | Cytoplasm | Y | N |
| EKNSTPSEPGSGR<br>: ::: :<br>EAYEMPSEEGYQD | 196 | 240 | 27 | 5 | 0 | 8 | 3 | 3 | HDGF_HUMAN | N | Hepatoma-derived growth factor | Cytoplasm, Nucleus and Secreted | Y | N |
| EPDCNPSEAASEE<br>: ::: ..<br>EAYEMPSEEGYQD | 158 | 783 | 87 | 4 | 2 | 7 | 3 | 3 | DDX21_HUMAN | Y | Nucleolar RNA helicase 2 | Cytoplasm, Mitochondrion and Nucleus | Y | N |
| TVRYAPSEAGLHE<br> ::: : ..<br>EAYEMPSEEGYQD | 1829 | 2647 | 280 | 4 | 2 | 7 | 3 | 3 | FLNA_HUMAN | Y | Filamin-A | Cell projection, Cytoplasm and Perikaryon | Y | Y |
| PAERTPSEIQFHQ<br> : ::: ..<br>EAYEMPSEEGYQD | 322 | 820 | 92 | 4 | 2 | 7 | 3 | 3 | M4K2_HUMAN | Y | Mitogen-activated protein kinase kinase kinase kinase 2 | Basolateral cell membrane, Cytoplasm and Golgi apparatus membrane | N | N |
| AKDEPPSEGEAEE<br> : ::: ..<br>EAYEMPSEEGYQD | 466 | 543 | 62 | 4 | 2 | 7 | 3 | 3 | NFL_HUMAN | Y | Neurofilament light polypeptide | Cell projection and Cytoplasm | Y | Y |
| KPGEEPSEYTDEE<br> : ::: ..<br>EAYEMPSEEGYQD | 202 | 223 | 24 | 4 | 2 | 7 | 3 | 3 | PGRC2_HUMAN | Y | Membrane-associated progesterone receptor component 2 | Endoplasmic reticulum, Membrane and Nucleus envelope | Y | N |
| KRVRRPSESDKED<br> ::: .:<br>EAYEMPSEEGYQD | 621 | 1782 | 190 | 4 | 1 | 8 | 3 | 3 | AKA12_HUMAN | Y | A-kinase anchor protein 12 | Cytoplasm and Membrane | Y | Y |
| EDEGIPSENEEEK<br>: ::: .<br>EAYEMPSEEGYQD | 2339 | 2813 | 310 | 4 | 1 | 8 | 3 | 3 | AKP13_HUMAN | Y | A-kinase anchor protein 13 | Cytoplasm, Membrane and Nucleus | Y | N |
| HAFSSPSESPDST<br> :. :::<br>EAYEMPSEEGYQD | 1159 | 1429 | 160 | 4 | 1 | 8 | 3 | 3 | ANR50_HUMAN | N | Ankyrin repeat domain-containing protein 50 | Endosome | N | N |
| ERLQNPSESSEPI<br>: . :::<br>EAYEMPSEEGYQD | 25 | 185 | 22 | 4 | 1 | 8 | 3 | 3 | CASA1_HUMAN | N | Alpha-S1-casein | Secreted | N | N |
| HGAASPSEKGAHP<br> ::: : .<br>EAYEMPSEEGYQD | 13 | 602 | 66 | 4 | 1 | 8 | 3 | 3 | CKAP4_HUMAN | N | Cytoskeleton-associated protein 4 | Cell membrane, Cytoplasm and Endoplasmic reticulum membrane | Y | N |
| GSTYTPSEAGNEL<br> ::: : .<br>EAYEMPSEEGYQD | 208 | 256 | 29 | 4 | 1 | 8 | 3 | 3 | MAF1_HUMAN | N | Repressor of RNA polymerase III transcription MAF1 homolog | Cytoplasm and Nucleus | N | N |
| DADSEPSESESAS<br>.: :::<br>EAYEMPSEEGYQD | 502 | 814 | 92 | 4 | 1 | 8 | 3 | 3 | UBP45_HUMAN | N | Ubiquitin carboxyl-terminal hydrolase 45 | Photoreceptor inner segment | N | N |
| DGLDTPSENSNEF<br>. . ::: .<br>EAYEMPSEEGYQD | 83 | 314 | 36 | 3 | 3 | 7 | 3 | 3 | BNIP2_HUMAN | Y | BCL2/adenovirus E1B 19 kDa protein-interacting protein 2 | Cytoplasm | N | N |
| AVCDEPSEPEEEE<br> . ::: ..<br>EAYEMPSEEGYQD | 194 | 1360 | 150 | 3 | 3 | 7 | 3 | 3 | MSH6_HUMAN | N | DNA mismatch repair protein Msh6 | Chromosome and Nucleus | Y | N |
| DLSDCPSEPLSDE<br>. . ::: .<br>EAYEMPSEEGYQD | 211 | 1905 | 210 | 3 | 3 | 7 | 3 | 3 | MTCL1_HUMAN | N | Microtubule cross-linking factor 1 | Apical cell membrane, Cytoplasm, Lateral cell membrane and Midbody | N | Y # |
| RPKHRPSEADEEE<br> ::: ..<br>EAYEMPSEEGYQD | 652 | 1153 | 130 | 3 | 2 | 8 | 3 | 3 | AP3D1_HUMAN | N | AP-3 complex subunit delta-1 | Cytoplasm and Golgi apparatus membrane | Y | Y |
| NKFRTPSELHNDN<br> . ::: .<br>EAYEMPSEEGYQD | 461 | 566 | 60 | 3 | 2 | 8 | 3 | 3 | CSPG5_HUMAN | Y | Chondroitin sulfate proteoglycan 5 | Cell junction, Cell membrane, Cell surface, Endoplasmic reticulum membrane and Golgi apparatus membrane | N | Y |
| GTKPPPSEGSDEE<br> ::: ..<br>EAYEMPSEEGYQD | 113 | 446 | 49 | 3 | 2 | 8 | 3 | 3 | GRWD1_HUMAN | N | Glutamate-rich WD repeat-containing protein 1 | Chromosome and Nucleus | N | N |
| RIFQVPSEMTEDI<br> .. :::<br>EAYEMPSEEGYQD | 123 | 167 | 19 | 3 | 2 | 8 | 3 | 3 | SPT19_HUMAN | Y | Spermatogenesis-associated protein 19, mitochondrial | Mitochondrion outer membrane | N | N |

### Figure S13

#### B S87 human – part 1

| Peptide alignment | Position | Length | MW list [kDa] | Exact | Similar | Mismatch | Max exact stretch | Max exact + similar stretch | ID | Also in mouse | Description | Subcellular location | Detected in plasma | Detected in CSF |
|---|---|---|---|---|---|---|---|---|---|---|---|---|---|---|
| TVEGAGSIAAATG<br>:::::::::::::<br>TVEGAGSIAAATG | 81 | 140 | 14 | 13 | 0 | 0 | 13 | 13 | SYUA_HUMAN | N | Alpha-synuclein | Cell junction. Cytoplasm. Membrane. Nucleus. Secreted | Y | Y |
| VDRGAGSIREAGG<br> ::::: : :<br>TVEGAGSIAAATG | 33 | 106 | 12 | 7 | 0 | 6 | 5 | 5 | ATIF1_HUMAN | Y | ATPase inhibitor, mitochondrial | Mitochondrion | Y * | N |
| ADESTGSIAKRLQ<br> : ::::<br>TVEGAGSIAAATG | 33 | 364 | 39 | 5 | 0 | 8 | 4 | 4 | ALDOA_HUMAN | Y | Fructose-bisphosphate aldolase A | Cytoplasm | Y | Y |
| FDSLAGSIPATKV<br> :::: :<br>TVEGAGSIAAATG | 13 | 593 | 66 | 5 | 0 | 8 | 4 | 4 | CPNE5_HUMAN | Y | Copine-5 | Cell projection. Perikaryon | N | N |
| ATVLSGSIAGLEG<br> :::: :<br>TVEGAGSIAAATG | 981 | 2054 | 230 | 5 | 0 | 8 | 4 | 4 | MY18A_HUMAN | Y | Unconventional myosin-XVIIIa | Cell surface. Cytoplasm. Endoplasmic reticulum-Golgi intermediate compartment. Golgi apparatus. Postsynaptic Golgi apparatus | Y | N |
| LLEEQGSIALRQE<br> : ::::<br>TVEGAGSIAAATG | 1035 | 2472 | 280 | 5 | 0 | 8 | 4 | 4 | SPTN1_HUMAN | Y | Spectrin alpha chain, non-erythrocytic 1 | Cytoplasm | Y | Y |
| TEIRAGSISSEEV<br>: ::::<br>TVEGAGSIAAATG | 1695 | 3051 | 330 | 5 | 0 | 8 | 4 | 4 | BD1L1_HUMAN | N | Biorientation of chromosomes in cell division protein 1-like 1 | Chromosome | N | N |
| AEDSDGSIASWEL<br> . ::::<br>TVEGAGSIAAATG | 172 | 483 | 55 | 4 | 1 | 8 | 4 | 4 | SMRD3_HUMAN | Y | SWI/SNF-related matrix-associated actin-dependent regulator of chromatin subfamily D member 3 | Nucleus | N | N |
| ELSPAGSISKNSP<br> :::: .<br>TVEGAGSIAAATG | 617 | 639 | 73 | 4 | 1 | 8 | 4 | 4 | PARN_HUMAN | N | Poly(A)-specific ribonuclease PARN | Cytoplasm. Nucleus | N | N |
| HSPRAGSISPGSP<br> :::: .<br>TVEGAGSIAAATG | 824 | 1409 | 150 | 4 | 1 | 8 | 4 | 4 | TNS2_HUMAN | N | Tensin-2 | Cell junction. Cell membrane | N | N |
| ALKEAGSIVRLYV<br> ::::<br>TVEGAGSIAAATG | 136 | 724 | 81 | 4 | 0 | 9 | 4 | 4 | DLG4_HUMAN | Y | Disks large homolog 4 | Cell junction. Cell membrane. Cell projection. Cytoplasm | N | Y |
| FSTFAGSITGPLY<br> ::::<br>TVEGAGSIAAATG | 412 | 916 | 100 | 4 | 0 | 9 | 4 | 4 | NFM_HUMAN | N | Neurofilament medium polypeptide | Cell projection. Cytoplasm | Y | Y |
| IQARLGSIAEIDL<br> ::::<br>TVEGAGSIAAATG | 130 | 876 | 100 | 4 | 0 | 9 | 4 | 4 | SRRT_HUMAN | Y | Serrate RNA effector molecule homolog | Cytoplasm. Nucleus | Y | Y # |
| QLNRAGSISTLDS<br> ::::<br>TVEGAGSIAAATG | 174 | 1216 | 130 | 4 | 0 | 9 | 4 | 4 | RELCH_HUMAN | Y | RAB11-binding protein RELCH | Golgi apparatus. Recycling endosome | N | N |
| NLSEAGSIKKGER<br> ::::<br>TVEGAGSIAAATG | 198 | 1438 | 160 | 4 | 0 | 9 | 4 | 4 | CLIP1_HUMAN | N | CAP-Gly domain-containing linker protein 1 | Cell projection. Cytoplasm. Cytoplasmic vesicle membrane | Y | N |
| TVSCTGSIRYKTL<br>:: ::: :<br>TVEGAGSIAAATG | 600 | 763 | 83 | 6 | 0 | 7 | 3 | 3 | PLPR4_HUMAN | N | Phospholipid phosphatase-related protein type 4 | Membrane | N | N |
| SASSEGSIHVAMG<br>. ::: : :<br>TVEGAGSIAAATG | 164 | 1011 | 110 | 5 | 1 | 7 | 3 | 3 | RASL3_HUMAN | N | RAS protein activator like-3 | Cytoplasm | N | N |
| EVIRKGSITEYTA<br> : ::: :<br>TVEGAGSIAAATG | 108 | 314 | 36 | 5 | 0 | 8 | 3 | 3 | BNIP2_HUMAN | Y | BCL2/adenovirus E1B 19 kDa protein-interacting protein 2 | Cytoplasm | N | N |

### Figure S13

#### B S87 human – part 2

| Peptide alignment | Position | Length | MW list [kDa] | Exact | Similar | Mismatch | Max exact stretch | Max exact + similar stretch | ID | Also in mouse | Description | Subcellular location | Detected in plasma | Detected in CSF |
|---|---|---|---|---|---|---|---|---|---|---|---|---|---|---|
| TALGGGSIRQSPL<br>: : :::<br>TVEGAGSIAAATG | 228 | 503 | 51 | 5 | 0 | 8 | 3 | 3 | WIPF1_HUMAN | N | WAS/WASL-interacting protein family member 1 | Cell projection, Cytoplasm, Cytoplasmic vesicle | Y | N |
| AEEDNGSIGEETD<br> : ::: :<br>TVEGAGSIAAATG | 654 | 685 | 77 | 5 | 0 | 8 | 3 | 3 | STIM1_HUMAN | Y | Stromal interaction molecule 1 | Cell membrane, Cytoplasm, Endoplasmic reticulum membrane, Sarcoplasmic reticulum | Y | Y |
| GNPGRGSIKTVAG<br> : ::: :<br>TVEGAGSIAAATG | 164 | 979 | 110 | 5 | 0 | 8 | 3 | 3 | UBP37_HUMAN | N | Ubiquitin carboxyl-terminal hydrolase 37 | Unspecified | N | N |
| IIPHSGSIEKAEI<br> . ::: :<br>TVEGAGSIAAATG | 219 | 372 | 42 | 4 | 1 | 8 | 3 | 3 | ZN830_HUMAN | N | Zinc finger protein 830 | Chromosome, Nucleus, Nucleus speckle | N | N |
| EEEEEGSIMNGST<br> : ::: .<br>TVEGAGSIAAATG | 558 | 675 | 81 | 4 | 1 | 8 | 3 | 3 | NEXN_HUMAN | Y | Nexilin | Cell junction and Cytoplasm | Y | N |
| ESQGKGSISEDEL<br> .: :::<br>TVEGAGSIAAATG | 344 | 776 | 84 | 4 | 1 | 8 | 3 | 3 | RTN1_HUMAN | N | Reticulon-1 | Endoplasmic reticulum membrane,<br>Golgi apparatus membrane | Y | Y |
| NKYGRGSISLNSS<br> : ::: .<br>TVEGAGSIAAATG | 679 | 800 | 91 | 4 | 1 | 8 | 3 | 3 | M3K20_HUMAN | N | Mitogen-activated protein kinase kinase kinase 20 | Cytoplasm and Nucleus | N | N |
| SLSKEGSIGGSGG<br>. ::: :<br>TVEGAGSIAAATG | 1108 | 1343 | 150 | 4 | 1 | 8 | 3 | 3 | SYGP1_HUMAN | Y | Ras/Rap GTPase-activating protein SynGAP | Unspecified | N | N |
| SVATVGSICDLNL<br>.: :::<br>TVEGAGSIAAATG | 2107 | 2602 | 280 | 4 | 1 | 8 | 3 | 3 | FLNB_HUMAN | N | Filamin-B | Cytoplasm | Y | Y |
| MVLSRGSIFPMSV<br> : ::: .<br>TVEGAGSIAAATG | 931 | 2701 | 310 | 4 | 1 | 8 | 3 | 3 | ITPR2_HUMAN | Y | Inositol 1,4,5-trisphosphate receptor type 2 | Endoplasmic reticulum membrane | N | N |
| GTDIGGSIRFPSS<br> . ::: .<br>TVEGAGSIAAATG | 235 | 579 | 63 | 3 | 2 | 8 | 3 | 3 | FAAH1_HUMAN | Y | Fatty-acid amide hydrolase 1 | Cytoplasm and Endomembrane system | N | N |
| SGSHQGSIQELST<br>. ::: .<br>TVEGAGSIAAATG | 546 | 630 | 71 | 3 | 2 | 8 | 3 | 3 | KCND2_HUMAN | N | Potassium voltage-gated channel subfamily D member 2 | Cell junction, Cell membrane, Cell projection, Perikaryon | N | N |
| ELQREGSIETLSN<br> . ::: .<br>TVEGAGSIAAATG | 880 | 928 | 100 | 3 | 2 | 8 | 3 | 3 | BCAS3_HUMAN | N | Breast carcinoma-amplified sequence 3 | Cytoplasm and Nucleus | N | Y # |
| SRLSRGSISSTSE<br>. ::: .<br>TVEGAGSIAAATG | 668 | 1821 | 210 | 3 | 2 | 8 | 3 | 3 | PHIP_HUMAN | N | PH-interacting protein | Nucleus | N | N |

#### Figure S13

##### B Y39 human

| Peptide alignment | Position | Length | MW list [kDa] | Exact | Similar | Mismatch | Max exact stretch | Max exact + similar stretch | ID | Also in mouse | Description | Subcellular location | Detected in plasma | Detected in CSF |
|---|---|---|---|---|---|---|---|---|---|---|---|---|---|---|
| EPDEILYVNMDEG<br> . .:::<br>TKEGVLYVGSKTK | 815 | 894 | 98 | 3 | 2 | 8 | 3 | 4 | UFO_HUMAN | Y | Tyrosine-protein kinase receptor UFO | Cell membrane | Y | Y |
| ASDGKLYVSSESR<br> .: ::: : ..<br>TKEGVLYVGSKTK | 179 | 1130 | 120 | 5 | 3 | 5 | 3 | 3 | ABL1_HUMAN | Y | Tyrosine-protein kinase ABL1 | Cytoplasm, Mitochondrion, Nucleus, Nuclear membrane | Y * | N |
| DHEEHLYVNTQGL<br> : ::: .<br>TKEGVLYVGSKTK | 408 | 582 | 62 | 4 | 1 | 8 | 3 | 3 | SHC2_HUMAN | Y | SHC-transforming protein 2 | Unspecified | N | N |
| GGEDPLYVARRLV<br> : ::: .<br>TKEGVLYVGSKTK | 528 | 665 | 72 | 4 | 1 | 8 | 3 | 3 | WRIP1_HUMAN | Y | ATPase WRNIP1 | Cytoplasm and Nucleus | N | N |

##### B Y125 human

| Peptide alignment | Position | Length | MW list [kDa] | Exact | Similar | Mismatch | Max exact stretch | Max exact + similar stretch | ID | Also in mouse | Description | Subcellular location | Detected in plasma | Detected in CSF |
|---|---|---|---|---|---|---|---|---|---|---|---|---|---|---|
| DPDNEAYEMPSEE<br>:::::::::::::<br>DPDNEAYEMPSEE | 119 | 140 | 14 | 13 | 0 | 0 | 13 | 13 | SYUA_HUMAN | Y | Alpha-synuclein | Cell junction, Cytoplasm, Membrane, Nucleus, Secreted | Y | Y |
| HYNGEAYEDDEHH<br> . ::::<br>DPDNEAYEMPSEE | 375 | 397 | 45 | 4 | 1 | 8 | 4 | 4 | DNJA1_HUMAN | Y | DnaJ homolog subfamily A member 1 | Cytoplasm, Membrane, Microsome, Mitochondrion, Nucleus | Y | Y |
| YVDLQAYEDPAQG<br> : .::: : .<br>DPDNEAYEMPSEE | 599 | 976 | 110 | 5 | 2 | 6 | 3 | 4 | EPHA1_HUMAN | Y | Ephrin type-A receptor 1 | Cell membrane | Y | N |
| DLEVPAYEDIFRD<br>: . ::: .<br>DPDNEAYEMPSEE | 124 | 566 | 66 | 4 | 2 | 7 | 3 | 3 | CDC45_HUMAN | N | Cell division control protein 45 homolog | Cytoplasm, Nucleus | N | N |
| FKIKNAYEESLDQ<br> ::: ..<br>DPDNEAYEMPSEE | 1488 | 1941 | 220 | 3 | 2 | 8 | 3 | 3 | MYH2_HUMAN | N | Myosin-2 | Cytoplasm | Y | Y # |

##### B Y133 human

| Peptide alignment | Position | Length | MW list [kDa] | Exact | Similar | Mismatch | Max exact stretch | Max exact + similar stretch | ID | Also in mouse | Description | Subcellular location | Detected in plasma | Detected in CSF |
|---|---|---|---|---|---|---|---|---|---|---|---|---|---|---|
| TPSAEGYQDVRDR<br> :: :::::<br>MPSEEGYQDYEPE | 433 | 778 | 88 | 7 | 0 | 6 | 5 | 5 | ABLM1_HUMAN | Y | Actin-binding LIM protein 1 | Cytoplasm | Y | N |
| SAPTSGYQEFVHA<br> :::..<br>MPSEEGYQDYEPE | 569 | 825 | 90 | 3 | 2 | 8 | 3 | 5 | IL4RA_HUMAN | Y | Interleukin-4 receptor subunit alpha | Cell membrane, Secreted | Y | N |
| LLAKKGYQERDLE<br>. :::. . :<br>MPSEEGYQDYEPE | 58 | 441 | 49 | 4 | 3 | 6 | 3 | 4 | P2RX6_HUMAN | N | P2X purinoceptor 6 | Membrane | N | N |

### Figure S13

**B** **Y136 human**

| Peptide alignment | Position | Length | MW list [kDa] | Exact | Similar | Mismatch | Max exact stretch | Max exact + similar stretch | ID | Also in mouse | Description | Subcellular location | Detected in plasma | Detected in CSF |
|---|---|---|---|---|---|---|---|---|---|---|---|---|---|---|
| GGYVQDYEDFM*<br> :::: :<br>EEGYQDYEPEA* | 248 | 258 | 29 | 5 | 0 | 7 | 4 | 4 | EIF3J_HUMAN | N | Eukaryotic translation initiation factor 3 subunit J | Cytoplasm | Y | Y |
| LEKIQDYEKMPE<br> : ::::<br>EEGYQDYEPEA* | 490 | 569 | 65 | 5 | 0 | 7 | 4 | 4 | IL1R1_HUMAN | Y | Interleukin-1 receptor type 1 | Cell membrane, Membrane, Secreted | Y | Y |
| PVPNPDYEPIRK<br> ::::<br>EEGYQDYEPEA* | 182 | 207 | 23 | 4 | 0 | 8 | 4 | 4 | CD3E_HUMAN | Y | T-cell surface glycoprotein CD3 epsilon chain | Cell membrane | N | N |
| MLRLQDYEEKTK<br> ::::<br>EEGYQDYEPEA* | 348 | 586 | 69 | 4 | 0 | 8 | 4 | 4 | EZRI_HUMAN | Y | Ezrin | Apical cell membrane, Cell projection, Cytoplasm | Y | Y |
| EEDEEDYESSAK<br>:: .::: :<br>EEGYQDYEPEA* | 64 | 1025 | 120 | 6 | 1 | 5 | 3 | 4 | LCAP_HUMAN | Y | Leucyl-cystinyl aminopeptidase | Cell membrane and Secreted | Y | Y |
| VVGGEDYEEVDR<br> : .:::<br>EEGYQDYEPEA* | 300 | 904 | 99 | 4 | 1 | 7 | 3 | 4 | ZN598_HUMAN | N | E3 ubiquitin-protein ligase ZNF598 | Unspecified | N | N |
| DDSDEDYEKVPL<br>.. .:::<br>EEGYQDYEPEA* | 442 | 561 | 62 | 3 | 3 | 6 | 3 | 4 | 3BP2_HUMAN | Y | SH3 domain-binding protein 2 | Unspecified | N | N |
| DLDDEDYEEDTP<br>. .::: .<br>EEGYQDYEPEA* | 166 | 391 | 44 | 3 | 3 | 6 | 3 | 4 | REQU_HUMAN | Y | Zinc finger protein ubi-d4 | Cytoplasm and Nucleus | N | N |
| DDDEHDYEEILE<br>.. .:::<br>EEGYQDYEPEA* | 626 | 643 | 72 | 3 | 3 | 6 | 3 | 4 | THMS2_HUMAN | Y | Protein THEMIS2 | Unspecified | N | N |
| PTDNEDYEHDDE<br> .::: .<br>EEGYQDYEPEA* | 168 | 561 | 62 | 3 | 2 | 7 | 3 | 4 | 3BP2_HUMAN | Y | SH3 domain-binding protein 2 | Unspecified | N | N |
| EEGAPDYENLQE<br>::: :::<br>EEGYQDYEPEA* | 249 | 262 | 28 | 6 | 0 | 6 | 3 | 3 | LAT_HUMAN | N | Linker for activation of T-cells family member 1 | Cell membrane | Y | N |
| EELQDDYEDMME<br>:: :::<br>EEGYQDYEPEA* | 2 | 911 | 100 | 5 | 0 | 7 | 3 | 3 | B3AT_HUMAN | N | Band 3 anion transport protein | Basolateral cell membrane and Cell membrane | Y | Y |
| EEPEPDYEAIQT<br>:: :::<br>EEGYQDYEPEA* | 381 | 432 | 47 | 5 | 0 | 7 | 3 | 3 | PHAG1_HUMAN | Y | Phosphoprotein associated with glycosphingolipid-enriched microdomains 1 | Cell membrane | Y * | Y |
| DEPEGDYEEVLE<br>.: :::<br>EEGYQDYEPEA* | 391 | 486 | 54 | 4 | 1 | 7 | 3 | 3 | HCLS1_HUMAN | Y | Hematopoietic lineage cell-specific protein | Cytoplasm, Membrane and Mitochondrion | Y | N |
| TSNFDDYEEEEI<br> . ::: :<br>EEGYQDYEPEA* | 325 | 351 | 41 | 4 | 1 | 7 | 3 | 3 | KAPCA_HUMAN | Y | cAMP-dependent protein kinase catalytic subunit alpha | Cell membrane, Cell projection, Cytoplasm, Cytoplasmic vesicle, Membrane, Mitochondrion and Nucleus | Y | N |
| TSNFDDYEEEDI<br> . ::: :<br>EEGYQDYEPEA* | 325 | 351 | 41 | 4 | 1 | 7 | 3 | 3 | KAPCB_HUMAN | N | cAMP-dependent protein kinase catalytic subunit beta | Cell membrane, Cytoplasm, Membrane and Nucleus | Y | N |
| SSKEDDYESDAA<br> ::: .:<br>EEGYQDYEPEA* | 364 | 5183 | 570 | 4 | 1 | 7 | 3 | 3 | UBR4_HUMAN | Y | E3 ubiquitin-protein ligase UBR4 | Cytoplasm, Membrane, Nucleus | Y | N |
| DPERSDYEEQQL<br>. ::: .<br>EEGYQDYEPEA* | 65 | 394 | 44 | 3 | 2 | 7 | 3 | 3 | ALKB5_HUMAN | N | RNA demethylase ALKBH5 | Nucleus speckle | N | N |
| DDFDSDYENPDE<br>.. :::<br>EEGYQDYEPEA* | 66 | 456 | 50 | 3 | 2 | 7 | 3 | 3 | BLNK_HUMAN | Y | B-cell linker protein | Cell membrane and Cytoplasm | N | Y |
| DGPAVDYENQDV<br>. ::: .<br>EEGYQDYEPEA* | 316 | 403 | 45 | 3 | 2 | 7 | 3 | 3 | STAP2_HUMAN | N | Signal-transducing adaptor protein 2 | Cytoplasm | N | N |

**Figure S13. List of proteins that could cross-react with antibodies raised against human phosphorylated aSyn fragment** **(related to Figure 9)**

**A**. Summary of the cross-reacting proteins that were previously detected in human plasma and Cerebrospinal fluid (CSF). **B**. For each fragment of 13 residues centred on a phosphorylated residue (S129, S87, Y39, Y125, Y133, Y136 ), the list of candidate hits is provided with local alignment against aSyn,  starting position of hit alignment, length of the protein hit, its molecular weight, the number of exact matches, similar matches, and mismatches, the length of the largest continuous exact matches stretch and the length of the largest continuous exact or similar matches stretch. Uniprot ID, description and subcellular localization retrieved by parsing the "CC -!- SUBCELLULAR LOCATION:" fields of the corresponding Uniprot entries is also provided. Y- detected; N- not detected; Y*- detected by Proximity Extension Assay (PEA) technology; Y# - sequence coverage is reported as 0% in the CSF proteome resource database.

### Figure S14

#### S129 mouse – part 1

| Peptide alignment | Position | Length | MW list [kDa] | Exact | Similar | Mismatch | Max exact stretch | Max exact + similar stretch | ID | Also in human | Description | Subcellular location |
|---|---|---|---|---|---|---|---|---|---|---|---|---|
| EAYEMPSEEGYQD<br>:::::::::::::<br>EAYEMPSEEGYQD | 123 | 140 | 14 | 13 | 0 | 0 | 13 | 13 | SYUA_MOUSE | Y | Alpha-synuclein | Cell junction, Cytoplasm, Membrane, Nucleus and Secreted |
| KARENPSEEAQNL<br> : : ::::<br>EAYEMPSEEGYQD | 115 | 501 | 59 | 6 | 0 | 7 | 4 | 4 | SF3A3_MOUSE | Y | Splicing factor 3A subunit 3 | Nucleus |
| LPEAPPSEEEEQE<br> :::: :.<br>EAYEMPSEEGYQD | 810 | 1128 | 130 | 5 | 1 | 7 | 4 | 4 | TEX2_MOUSE | Y | Testis-expressed protein 2 | Endoplasmic reticulum membrane and Nucleus membrane |
| PPARRPSEEPEPD<br> :::: :<br>EAYEMPSEEGYQD | 373 | 429 | 47 | 5 | 0 | 8 | 4 | 4 | PHAG1_MOUSE | Y | Phosphoprotein associated with glycosphingolipid-enriched microdomains 1 | Cell membrane |
| KIQRMPSESAAQS<br> :::: :<br>EAYEMPSEEGYQD | 113 | 513 | 57 | 5 | 0 | 8 | 4 | 4 | PLRG1_MOUSE | Y | Pleiotropic regulator 1 | Nucleus and Nucleus speckle |
| IITEEPSEEEADM<br> : ::::<br>EAYEMPSEEGYQD | 112 | 851 | 94 | 5 | 0 | 8 | 4 | 4 | DDX21_MOUSE | Y | Nucleolar RNA helicase 2 | Cytoplasm, Mitochondrion and Nucleus |
| SSPEQPSEEW***<br> : ::::<br>EAYEMPSEEGYQD | 917 | 926 | 100 | 5 | 0 | 8 | 4 | 4 | NIBA1_MOUSE | Y | Protein Niban 1 | Cytoplasm and Membrane |
| PRESRPSEEREWD<br> :::: :<br>EAYEMPSEEGYQD | 1153 | 1344 | 160 | 5 | 0 | 8 | 4 | 4 | EIF3A_MOUSE | Y | Eukaryotic translation initiation factor 3 subunit A | Cytoplasm and Nucleus |
| ESGRSPSEEEEDA<br>: ::::<br>EAYEMPSEEGYQD | 308 | 1693 | 190 | 5 | 0 | 8 | 4 | 4 | ARG28_MOUSE | N | Rho guanine nucleotide exchange factor 28 | Cell membrane and Cytoplasm |
| DIIRQPSEEEIIK<br>. ::::<br>EAYEMPSEEGYQD | 110 | 130 | 15 | 4 | 1 | 8 | 4 | 4 | PEA15_MOUSE | Y | Astrocytic phosphoprotein PEA-15 | Cytoplasm |
| PDRTPPSEEDSAE<br> :::: .<br>EAYEMPSEEGYQD | 79 | 315 | 34 | 4 | 1 | 8 | 4 | 4 | SGTA_MOUSE | Y | Small glutamine-rich tetratricopeptide repeat-containing protein alpha | Cytoplasm and Nucleus |
| LVQHSPSEEKMSP<br> ::::<br>EAYEMPSEEGYQD | 140 | 262 | 29 | 4 | 0 | 9 | 4 | 4 | EAF2_MOUSE | N | ELL-associated factor 2 | Nucleus speckle |
| KELPPPSEEIKTG<br> ::::<br>EAYEMPSEEGYQD | 531 | 657 | 75 | 4 | 0 | 9 | 4 | 4 | THOC1_MOUSE | Y | THO complex subunit 1 | Cytoplasm, Nucleus, Nucleus matrix and Nucleus speckle |
| VTPGWPSEEEVGS<br> ::::<br>EAYEMPSEEGYQD | 447 | 733 | 79 | 4 | 0 | 9 | 4 | 4 | CEP68_MOUSE | N | Centrosomal protein of 68 kDa | Cytoplasm |
| NGLSQPSEEEADI<br> ::::<br>EAYEMPSEEGYQD | 149 | 851 | 94 | 4 | 0 | 9 | 4 | 4 | DDX21_MOUSE | Y | Nucleolar RNA helicase 2 | Cytoplasm, Mitochondrion and Nucleus |
| NGLSQPSEEEVDI<br> ::::<br>EAYEMPSEEGYQD | 186 | 851 | 94 | 4 | 0 | 9 | 4 | 4 | DDX21_MOUSE | Y | Nucleolar RNA helicase 2 | Cytoplasm, Mitochondrion and Nucleus |
| AMLHLPSEQGTPE<br> .::::.: .<br>EAYEMPSEEGYQD | 37 | 412 | 48 | 4 | 3 | 6 | 3 | 6 | NEMO_MOUSE | Y | NF-kappa-B essential modulator | Cytoplasm and Nucleus |
| GTSELPSEDDGVE<br> :.:::. .<br>EAYEMPSEEGYQD | 447 | 2323 | 260 | 4 | 3 | 6 | 3 | 6 | C2CD3_MOUSE | N | C2 domain-containing protein 3 | Cytoplasm |

#### Figure S14

##### S129 mouse – part 2

| Peptide alignment | Position | Length | MW list [kDa] | Exact | Similar | Mismatch | Max exact stretch | Max exact + similar stretch | ID | Also in human | Description | Subcellular location |
|---|---|---|---|---|---|---|---|---|---|---|---|---|
| EEAGNPSEDGMQS<br>: :::.: :<br>EAYEMPSEEGYQD | 52 | 501 | 56 | 6 | 1 | 6 | 3 | 5 | PWP1_MOUSE | N | Periodic tryptophan protein 1 homolog | Chromosome and Nucleus |
| ASRTLPSEDEEGE<br> .:::. .<br>EAYEMPSEEGYQD | 8302 | 8799 | 1000 | 3 | 3 | 7 | 3 | 5 | SYNE1_MOUSE | Y | Nesprin-1 | Cytoplasm, Nucleus, Nucleus envelope, Nucleus outer membrane |
| DSSPLPSEKSDED<br>. .::: .:<br>EAYEMPSEEGYQD | 1354 | 1711 | 200 | 4 | 3 | 6 | 3 | 4 | CHD1_MOUSE | Y | Chromodomain-helicase-DNA-binding protein 1 | Cytoplasm and Nucleus |
| LQPGGPSEDECSD<br> :::. :<br>EAYEMPSEEGYQD | 263 | 332 | 37 | 4 | 1 | 8 | 3 | 4 | PINX1_MOUSE | N | PIN2/TERF1-interacting telomerase inhibitor 1 | Chromosome and Nucleus |
| ETQRKPSEDEKGF<br>: :::.<br>EAYEMPSEEGYQD | 1055 | 1231 | 140 | 4 | 1 | 8 | 3 | 4 | EHBP1_MOUSE | Y | EH domain-binding protein 1 | Cytoplasm, Endosome and Membrane |
| YSVQEPSEDSEEE<br> . :::. ..<br>EAYEMPSEEGYQD | 1169 | 1365 | 140 | 3 | 4 | 6 | 3 | 4 | LMTK1_MOUSE | Y | Serine/threonine-protein kinase LMTK1 | Cell projection, Cytoplasm and Membrane |
| QYNQVPSEDFERA<br>. . :::.<br>EAYEMPSEEGYQD | 317 | 365 | 40 | 3 | 3 | 7 | 3 | 4 | CXAR_MOUSE | N | Coxsackievirus and adenovirus receptor homolog | Basolateral cell membrane, Cell junction, Cell membrane and Secreted |
| DSSRLPSEGPQPA<br>. .:::<br>EAYEMPSEEGYQD | 36 | 206 | 22 | 3 | 2 | 8 | 3 | 4 | BLVRB_MOUSE | Y | Flavin reductase (NADPH) | Cytoplasm |
| RHQSDPSEDEDER<br> :::. .<br>EAYEMPSEEGYQD | 123 | 496 | 57 | 3 | 2 | 8 | 3 | 4 | SGK3_MOUSE | Y | Serine/threonine-protein kinase Sgk3 | Cytoplasmic vesicle, Early endosome and Recycling endosome |
| DQQSLPSEPEEAL<br>. .:::<br>EAYEMPSEEGYQD | 336 | 541 | 58 | 3 | 2 | 8 | 3 | 4 | RETR2_MOUSE | N | Reticulophagy regulator 2 | Membrane |
| KSVSTPSEAGSQD<br> ::: : ::<br>EAYEMPSEEGYQD | 85 | 612 | 68 | 6 | 0 | 7 | 3 | 3 | DC1I2_MOUSE | N | Cytoplasmic dynein 1 intermediate chain 2 | Cytoplasm |
| EEEEEPSEGVDEE<br>: : ::: ..<br>EAYEMPSEEGYQD | 337 | 1044 | 120 | 5 | 2 | 6 | 3 | 3 | GGYF1_MOUSE | Y | GRB10-interacting GYF protein 1 | Unspecified |
| DLKESPSEGSLQP<br>. : ::: :<br>EAYEMPSEEGYQD | 2 | 512 | 58 | 5 | 1 | 7 | 3 | 3 | ASIC2_MOUSE | N | Acid-sensing ion channel 2 | Cell membrane |
| PWSKRPSECGCEE<br> ::: : ..<br>EAYEMPSEEGYQD | 57 | 80 | 9 | 4 | 2 | 7 | 3 | 3 | ADIG_MOUSE | N | Adipogenin | Membrane and Nucleus |
| KPGEEPSEYTDEE<br> : ::: ..<br>EAYEMPSEEGYQD | 196 | 217 | 23 | 4 | 2 | 7 | 3 | 3 | PGRC2_MOUSE | Y | Membrane-associated progesterone receptor component 2 | Endoplasmic reticulum, Membrane and Nucleus envelope |
| EGLDTPSENSDEF<br>: . ::: .<br>EAYEMPSEEGYQD | 83 | 326 | 38 | 4 | 2 | 7 | 3 | 3 | BNIP2_MOUSE | Y | BCL2/adenovirus E1B 19 kDa protein-interacting protein 2 | Cytoplasm |
| AKDEPPSEGEAEE<br> : ::: ..<br>EAYEMPSEEGYQD | 467 | 543 | 62 | 4 | 2 | 7 | 3 | 3 | NFL_MOUSE | Y | Neurofilament light polypeptide | Cell projection and Cytoplasm |
| PAERTPSEIQFHQ<br> : ::: ..<br>EAYEMPSEEGYQD | 322 | 821 | 91 | 4 | 2 | 7 | 3 | 3 | M4K2_MOUSE | Y | Mitogen-activated protein kinase kinase kinase kinase 2 | Basolateral cell membrane, Cytoplasm and Golgi apparatus membrane |
| TVRYSPSEAGLHE<br> ::: : ..<br>EAYEMPSEEGYQD | 1829 | 2647 | 280 | 4 | 2 | 7 | 3 | 3 | FLNA_MOUSE | Y | Filamin-A | Cell projection, Cytoplasm and Perikaryon |

### Figure S14

#### S129 mouse – part 3

| Peptide alignment | Position | Length | MW list [kDa] | Exact | Similar | Mismatch | Max exact stretch | Max exact + similar stretch | ID | Also in human | Description | Subcellular location |
|---|---|---|---|---|---|---|---|---|---|---|---|---|
| EVRKKPSEGEEEA<br>: ::: .<br>EAYEMPSEEGYQD | 162 | 192 | 21 | 4 | 1 | 8 | 3 | 3 | CY24A_MOUSE | Y | Cytochrome b-245 light chain | Cell membrane |
| EDEGIPSENEEEK<br>: ::: .<br>EAYEMPSEEGYQD | 2302 | 2776 | 300 | 4 | 1 | 8 | 3 | 3 | AKP13_MOUSE | Y | A-kinase anchor protein 13 | Cytoplasm, Membrane and Nucleus |
| AVCDEPSEPEEEE<br> . ::: ..<br>EAYEMPSEEGYQD | 194 | 1358 | 150 | 3 | 3 | 7 | 3 | 3 | MSH6_MOUSE | N | DNA mismatch repair protein Msh6 | Chromosome and Nucleus |
| RIFQVPSEVMDDV<br> .. :::<br>EAYEMPSEEGYQD | 110 | 154 | 18 | 3 | 2 | 8 | 3 | 3 | SPT19_MOUSE | Y | Spermatogenesis-associated protein 19, mitochondrial | Mitochondrion outer membrane |
| FMKKAPSEIDPEE<br> ::: ..<br>EAYEMPSEEGYQD | 45 | 386 | 44 | 3 | 2 | 8 | 3 | 3 | TTC4_MOUSE | N | Tetratricopeptide repeat protein 4 | Cytoplasm and Nucleus |
| NKFRTPSELHNDN<br> . ::: .<br>EAYEMPSEEGYQD | 461 | 566 | 60 | 3 | 2 | 8 | 3 | 3 | CSPG5_MOUSE | Y | Chondroitin sulfate proteoglycan 5 | Cell junction, Cell membrane, Cell surface, Endoplasmic reticulum membrane and Golgi apparatus membrane |
| LEDKAPSEVAIEE<br> ::: ..<br>EAYEMPSEEGYQD | 506 | 758 | 84 | 3 | 2 | 8 | 3 | 3 | TDIF2_MOUSE | N | Deoxynucleotidyltransferase terminal-interacting protein 2 | Nucleus |
| KRVRRPSESDKEE<br> ::: ..<br>EAYEMPSEEGYQD | 607 | 1684 | 180 | 3 | 2 | 8 | 3 | 3 | AKA12_MOUSE | Y | A-kinase anchor protein 12 | Cytoplasm. Membrane |

### Figure S14

#### S87 mouse– part 1

| Peptide alignment | Position | Length | MW list [kDa] | Exact | Similar | Mismatch | Max exact stretch | Max exact + similar stretch | ID | Also in human | Description | Subcellular location |
|---|---|---|---|---|---|---|---|---|---|---|---|---|
| MDTGAGSIREAGG<br> ::::: : :<br>TVEGAGSIAAATG | 33 | 106 | 12 | 7 | 0 | 6 | 5 | 5 | ATIF1_MOUSE | Y | ATPase inhibitor, mitochondrial | Mitochondrion |
| EPLGVGSIAAGGR<br> : :::::<br>TVEGAGSIAAATG | 350 | 509 | 57 | 6 | 0 | 7 | 5 | 5 | HARS1_MOUSE | N | Histidine--tRNA ligase, cytoplasmic | Cytoplasm |
| LLEEQGSIALRQG<br> : :::: :<br>TVEGAGSIAAATG | 1035 | 2472 | 280 | 6 | 0 | 7 | 4 | 4 | SPTN1_MOUSE | Y | Spectrin alpha chain, non-erythrocytic 1 | Cytoplasm |
| ADESTGSIAKRLQ<br> : ::::<br>TVEGAGSIAAATG | 33 | 364 | 39 | 5 | 0 | 8 | 4 | 4 | ALDOA_MOUSE | Y | Fructose-bisphosphate aldolase A | Cytoplasm |
| FDSLAGSIPATKV<br> :::: :<br>TVEGAGSIAAATG | 13 | 593 | 66 | 5 | 0 | 8 | 4 | 4 | CPNE5_MOUSE | Y | Copine-5 | Cell projection and Perikaryon |
| ATVLSGSIAGLEG<br> :::: :<br>TVEGAGSIAAATG | 977 | 2050 | 230 | 5 | 0 | 8 | 4 | 4 | MY18A_MOUSE | Y | Unconventional myosin-XVIIIa | Cytoplasm, Endoplasmic reticulum-Golgi intermediate compartment, Golgi apparatus and Postsynaptic Golgi apparatus |
| AEDSDGSIASWEL<br> . ::::<br>TVEGAGSIAAATG | 172 | 483 | 55 | 4 | 1 | 8 | 4 | 4 | SMRD3_MOUSE | Y | SWI/SNF-related matrix-associated actin-dependent regulator of chromatin subfamily D member 3 | Nucleus |
| ALKEAGSIVRLYV<br> ::::<br>TVEGAGSIAAATG | 136 | 724 | 80 | 4 | 0 | 9 | 4 | 4 | DLG4_MOUSE | Y | Disks large homolog 4 | Cell junction, Cell membrane, Cell projection and Cytoplasm |
| QLNRAGSISTLDS<br> ::::<br>TVEGAGSIAAATG | 174 | 1216 | 130 | 4 | 0 | 9 | 4 | 4 | RELCH_MOUSE | Y | RAB11-binding protein RELCH | Golgi apparatus and Recycling endosome |
| IQARLGSIAEIDL<br> ::::<br>TVEGAGSIAAATG | 130 | 875 | 100 | 4 | 0 | 9 | 4 | 4 | SRRT_MOUSE | Y | Serrate RNA effector molecule homolog | Cytoplasm and Nucleus |
| RVERGGSISEASG<br> :: ::: :.:<br>TVEGAGSIAAATG | 217 | 332 | 36 | 7 | 1 | 5 | 3 | 3 | JSPR1_MOUSE | N | Junctional sarcoplasmic reticulum protein 1 | Endoplasmic reticulum membrane and Sarcoplasmic reticulum membrane |
| TVTCTGSIRYKTL<br>:: ::: :<br>TVEGAGSIAAATG | 602 | 766 | 83 | 6 | 0 | 7 | 3 | 3 | PLPR4_MOUSE | N | Phospholipid phosphatase-related protein type 4 | Membrane |
| EVIRKGSITEYTA<br> : ::: :<br>TVEGAGSIAAATG | 108 | 326 | 38 | 5 | 0 | 8 | 3 | 3 | BNIP2_MOUSE | Y | BCL2/adenovirus E1B 19 kDa protein-interacting protein 2 | Cytoplasm |
| AEEDNGSIGEETD<br> : ::: :<br>TVEGAGSIAAATG | 654 | 685 | 78 | 5 | 0 | 8 | 3 | 3 | STIM1_MOUSE | Y | Stromal interaction molecule 1 | Cell membrane, Cytoplasm, Endoplasmic reticulum membrane and Sarcoplasmic reticulum |
| LVPHSGSIEKAEI<br> : ::: :<br>TVEGAGSIAAATG | 210 | 363 | 41 | 5 | 0 | 8 | 3 | 3 | ZN830_MOUSE | N | Zinc finger protein 830 | Chromosome, Nucleus and Nucleus speckle |
| SVSRRGSISSVSP<br>.: ::: .<br>TVEGAGSIAAATG | 478 | 729 | 79 | 4 | 2 | 7 | 3 | 3 | DTL_MOUSE | N | Denticleless protein homolog | Chromosome, Cytoplasm, Nucleus and Nucleus membrane |
| TPQRSGSISNYRS<br>: . :::<br>TVEGAGSIAAATG | 490 | 559 | 64 | 4 | 1 | 8 | 3 | 3 | AAPK1_MOUSE | N | 5'-AMP-activated protein kinase catalytic subunit alpha-1 | Cytoplasm and Nucleus |
| SVYFCGSIRGGRE<br>.: :::<br>TVEGAGSIAAATG | 11 | 173 | 19 | 4 | 1 | 8 | 3 | 3 | DNPH1_MOUSE | N | 2'-deoxynucleoside 5'-phosphate N-hydrolase 1 | Cytoplasm and Nucleus |

### Figure S14

#### S87 mouse – part 2

| Peptide alignment | Position | Length | MW list [kDa] | Exact | Similar | Mismatch | Max exact stretch | Max exact + similar stretch | ID | Also in human | Description | Subcellular location |
|---|---|---|---|---|---|---|---|---|---|---|---|---|
| MVLSRGSIFPVSV<br> : ::: .<br>TVEGAGSIAAATG | 931 | 2701 | 310 | 4 | 1 | 8 | 3 | 3 | ITPR2_MOUSE | Y | Inositol 1,4,5-trisphosphate receptor type 2 | Endoplasmic reticulum membrane |
| EEEEEGSIVNGST<br> : ::: .<br>TVEGAGSIAAATG | 489 | 607 | 72 | 4 | 1 | 8 | 3 | 3 | NEXN_MOUSE | Y | Nexilin | Cell junction and Cytoplasm |
| SLSKEGSIGGSGG<br>. ::: :<br>TVEGAGSIAAATG | 1105 | 1340 | 150 | 4 | 1 | 8 | 3 | 3 | SYGP1_MOUSE | Y | Ras/Rap GTPase-activating protein SynGAP | Cell junction and Membrane |
| EVDSLGSIHSIPS<br> :. :::<br>TVEGAGSIAAATG | 340 | 898 | 110 | 4 | 1 | 8 | 3 | 3 | TAOK3_MOUSE | N | Serine/threonine-protein kinase TAO3 | Cytoplasm |
| ELQREGSIETLSN<br> . ::: .<br>TVEGAGSIAAATG | 880 | 928 | 100 | 3 | 2 | 8 | 3 | 3 | BCAS3_MOUSE | N | Breast carcinoma-amplified sequence 3 homolog | Cytoplasm and Nucleus |
| GTDIGGSIRFPSA<br> . ::: .<br>TVEGAGSIAAATG | 235 | 579 | 63 | 3 | 2 | 8 | 3 | 3 | FAAH1_MOUSE | Y | Fatty-acid amide hydrolase 1 | Endoplasmic reticulum membrane and Golgi apparatus membrane |

### Figure S14

#### Y39 mouse

| Peptide alignment | Position | Length | MW list [kDa] | Exact | Similar | Mismatch | Max exact stretch | Max exact + similar stretch | ID | Also in human | Description | Subcellular location |
|---|---|---|---|---|---|---|---|---|---|---|---|---|
| EPDEILYVNMDEG<br> . .:::<br>TKEGVLYVGSKTK | 809 | 888 | 98 | 3 | 2 | 8 | 3 | 4 | UFO_MOUSE | Y | Tyrosine-protein kinase receptor UFO | Cell membrane |
| ASDGKLYVSSESR<br> .: ::: : ..<br>TKEGVLYVGSKTK | 179 | 1123 | 120 | 5 | 3 | 5 | 3 | 3 | ABL1_MOUSE | Y | Tyrosine-protein kinase ABL1 | Cytoplasm, Mitochondrion and Nucleus |
| KEVGGLYVASRAN<br> : ::: :.<br>TKEGVLYVGSKTK | 302 | 1147 | 130 | 5 | 1 | 7 | 3 | 3 | ASAP1_MOUSE | N | Arf-GAP with SH3 domain, ANK repeat and PH domain-containing protein 1 | Cytoplasm and Membrane |
| TRPEPLYVNLALG<br>:. :::<br>TKEGVLYVGSKTK | 1182 | 1305 | 140 | 4 | 1 | 8 | 3 | 3 | RHG33_MOUSE | N | Rho GTPase-activating protein 33 | Cell membrane |
| DYEEHLYVNTQGL<br> : ::: .<br>TKEGVLYVGSKTK | 389 | 573 | 62 | 4 | 1 | 8 | 3 | 3 | SHC2_MOUSE | Y | SHC-transforming protein 2 | Unspecified |
| GGEDPLYVARRLV<br> : ::: .<br>TKEGVLYVGSKTK | 523 | 660 | 72 | 4 | 1 | 8 | 3 | 3 | WRIP1_MOUSE | Y | ATPase WRNIP1 | Cytoplasm and Nucleus |

#### Y125 mouse

| Peptide alignment | Position | Length | MW list [kDa] | Exact | Similar | Mismatch | Max exact stretch | Max exact + similar stretch | ID | Also in human | Description | Subcellular location |
|---|---|---|---|---|---|---|---|---|---|---|---|---|
| DPGSEAYEMPSEE<br>:: :::::::::<br>DPDNEAYEMPSEE | 119 | 140 | 14 | 11 | 0 | 2 | 9 | 9 | SYUA_MOUSE | Y | Alpha-synuclein | Cell junction, Cytoplasm, Membrane, Nucleus and Secreted |
| EEEGEAYEEPDSE<br>. . :::: : :<br>DPDNEAYEMPSEE | 396 | 547 | 60 | 6 | 2 | 5 | 4 | 4 | CD19_MOUSE | N | B-lymphocyte antigen CD19 | Cell membrane and Membrane raft |
| HYNGEAYEDDEHH<br> . ::::<br>DPDNEAYEMPSEE | 375 | 397 | 45 | 4 | 1 | 8 | 4 | 4 | DNJA1_MOUSE | Y | DnaJ homolog subfamily A member 1 | Cytoplasm, Membrane, Microsome, Mitochondrion and Nucleus |
| YVDLQAYEDPAQG<br> : .::: : .<br>DPDNEAYEMPSEE | 600 | 977 | 110 | 5 | 2 | 6 | 3 | 4 | EPHA1_MOUSE | Y | Ephrin type-A receptor 1 | Cell membrane |
| SSGPDAYEEVTDG<br> .::: ..<br>DPDNEAYEMPSEE | 413 | 453 | 50 | 3 | 3 | 7 | 3 | 4 | PTN18_MOUSE | N | Tyrosine-protein phosphatase non-receptor type 18 | Cytoplasm and Nucleus |
| FKVKNAYEESLDQ<br> ::: ..<br>DPDNEAYEMPSEE | 1486 | 1939 | 220 | 3 | 2 | 8 | 3 | 3 | MYH4_MOUSE | N | Myosin-4 | Cytoplasm |

#### Figure S14

##### Y133 mouse

| Peptide alignment | Position | Length | MW list [kDa] | Exact | Similar | Mismatch | Max exact stretch | Max exact + similar stretch | ID | Also in human | Description | Subcellular location |
|---|---|---|---|---|---|---|---|---|---|---|---|---|
| TPSAEGYQDVRDR<br> :: :::::<br>MPSEEGYQDYEPE | 477 | 861 | 97 | 7 | 0 | 6 | 5 | 5 | ABLM1_MOUSE | Y | Actin-binding LIM protein 1 | Cytoplasm |
| PAPAGGYQEFVQA<br> ::::..<br>MPSEEGYQDYEPE | 569 | 810 | 88 | 3 | 2 | 8 | 3 | 5 | IL4RA_MOUSE | Y | Interleukin-4 receptor subunit alpha | Cell membrane and Secreted |
| MVTFQGYQKQTRW<br>: . .:::<br>MPSEEGYQDYEPE | 437 | 575 | 64 | 4 | 2 | 7 | 3 | 4 | I10R1_MOUSE | N | Interleukin-10 receptor subunit alpha | Cell membrane and Cytoplasm |

##### Y136 mouse

| Peptide alignment | Position | Length | MW list [kDa] | Exact | Similar | Mismatch | Max exact stretch | Max exact + similar stretch | ID | Also in human | Description | Subcellular location |
|---|---|---|---|---|---|---|---|---|---|---|---|---|
| GGYVQDYEDFM*<br> :::: :<br>EEGYQDYEPEA* | 253 | 263 | 29 | 5 | 0 | 7 | 4 | 4 | EI3JB_MOUSE | N | Eukaryotic translation initiation factor 3 subunit J-B | Cytoplasm |
| LEKIQDYEKMPD<br> : ::::<br>EEGYQDYEPEA* | 493 | 576 | 67 | 5 | 0 | 7 | 4 | 4 | IL1R1_MOUSE | Y | Interleukin-1 receptor type 1 | Cell membrane, Membrane and Secreted |
| PVPNPDYEPIRK<br> ::::<br>EEGYQDYEPEA* | 164 | 189 | 21 | 4 | 0 | 8 | 4 | 4 | CD3E_MOUSE | Y | T-cell surface glycoprotein CD3 epsilon chain | Cell membrane |
| MLRLQDYEQKTK<br> ::::<br>EEGYQDYEPEA* | 348 | 586 | 69 | 4 | 0 | 8 | 4 | 4 | EZRI_MOUSE | Y | Ezrin | Apical cell membrane, Cell projection and Cytoplasm |
| DEDEEDYESSAK<br>.: .::: :<br>EEGYQDYEPEA* | 64 | 1025 | 120 | 5 | 2 | 5 | 3 | 4 | LCAP_MOUSE | Y | Leucyl-cystinyl aminopeptidase | Cell membrane and Endomembrane system |
| EDSDEDYEKVPL<br>:. .:::<br>EEGYQDYEPEA* | 440 | 559 | 62 | 4 | 2 | 6 | 3 | 4 | 3BP2_MOUSE | Y | SH3 domain-binding protein 2 | Unspecified |
| DMDDHDYEEI**<br>. .::: :<br>EEGYQDYEPEA* | 654 | 663 | 74 | 4 | 2 | 6 | 3 | 4 | THMS2_MOUSE | Y | Protein THEMIS2 | Unspecified |
| PMDNEDYEHEDE<br> .::: :<br>EEGYQDYEPEA* | 168 | 559 | 62 | 4 | 1 | 7 | 3 | 4 | 3BP2_MOUSE | Y | SH3 domain-binding protein 2 | Unspecified |
| DLDDEDYEEDTP<br>. .::: .<br>EEGYQDYEPEA* | 166 | 391 | 44 | 3 | 3 | 6 | 3 | 4 | REQU_MOUSE | Y | Zinc finger protein ubi-d4 | Cytoplasm and Nucleus |
| ERGRSDYESVGS<br>: : :::<br>EEGYQDYEPEA* | 97 | 660 | 73 | 5 | 0 | 7 | 3 | 3 | DDX3L_MOUSE | N | Putative ATP-dependent RNA helicase Pl10 | Unspecified |
| EEPEPDYEAIQT<br>:: :::<br>EEGYQDYEPEA* | 380 | 429 | 47 | 5 | 0 | 7 | 3 | 3 | PHAG1_MOUSE | Y | Phosphoprotein associated with glycosphingolipid-enriched microdomains 1 | Cell membrane |
| EDAEGDYEDVLE<br>:. :::<br>EEGYQDYEPEA* | 399 | 486 | 54 | 4 | 1 | 7 | 3 | 3 | HCLS1_MOUSE | Y | Hematopoietic lineage cell-specific protein | Mitochondrion |
| TSNFDDYEEEEI<br> . ::: :<br>EEGYQDYEPEA* | 325 | 351 | 41 | 4 | 1 | 7 | 3 | 3 | KAPCA_MOUSE | Y | cAMP-dependent protein kinase catalytic subunit alpha | Cell membrane, Cell projection, Cytoplasm, Cytoplasmic vesicle, Membrane, Mitochondrion and Nucleus |
| ESWRNDYEEDDL<br>: ::: .<br>EEGYQDYEPEA* | 138 | 853 | 97 | 4 | 1 | 7 | 3 | 3 | TGS1_MOUSE | N | Trimethylguanosine synthase | Cytoplasm and Nucleus |
| SSKEDDYESDAA<br> ::: .:<br>EEGYQDYEPEA* | 364 | 5180 | 570 | 4 | 1 | 7 | 3 | 3 | UBR4_MOUSE | Y | E3 ubiquitin-protein ligase UBR4 | Cytoplasm, Membrane, Nucleus |
| DDFDSDYENPDE<br>.. :::<br>EEGYQDYEPEA* | 66 | 457 | 51 | 3 | 2 | 7 | 3 | 3 | BLNK_MOUSE | Y | B-cell linker protein | Cell membrane and Cytoplasm |

**Figure S14. List of proteins that could cross-react with antibodies raised against mouse phosphorylated aSyn fragment** (related to Figure 9)

For each fragment of 13 residues centred on a phosphorylated residue (S129, S87, Y39, Y125, Y133, Y136 ), the list of candidate hits is provided with local alignment against aSyn, starting position of hit alignment, length of the protein hit, its molecular weight, the number of exact matches, similar matches, and mismatches, the length of the largest continuous exact matches stretch and the length of the largest continuous exact or similar matches stretch. Uniprot ID, description and subcellular localization retrieved by parsing the "CC -!- SUBCELLULAR LOCATION:" fields of the corresponding Uniprot entries is also provided.

#### Figure S15

### A

| | | | | pSyn #64 | EP 1536Y | MJF-R13 | 81A | GTX | LASH-EGT | Figures |
|---|---|---|---|---|---|---|---|---|---|---|
| KO mice | Primary neurons | ICC | Neurites | + | +/- | + | + | +/- | + | Figure S8 |
| KO mice | Primary neurons | ICC | Cell bodies | + | +/- | + | + | +/- | + | Figure S8 |
| KO mice | Primary neurons | ICC | Nucleus | + | - | + | - | - | + | Figure S8 |
| KO mice | Primary neurons | WB | HMWs > 25 kDa | + | + | + | + | - | - | Figure 5 |
| KO mice | Primary neurons | WB | 17 kDa | + | + | + | + | + | - | Figure 5 |
| KO mice | Primary neurons | WB | 15 kDa | - | - | - | - | - | - | Figure 5 |
| KO mice | Primary neurons | WB | < 15 kDa | - | - | - | - | - | - | Figure 5 |
| KO mice | Mouse tissues | IHC | Neurites | - | - | - | + | +/- | - | Figure 7A |
| KO mice | Mouse tissues | IHC | Cell bodies | +/- | +/- | + | + | - | - | Figure 7A |
| KO mice | Mouse tissues | IHC | Nucleus | + | - | + | - | - | +/- | Figure 7A |
| KO mice | Mouse tissues | WB | HMWs > 25 kDa | + | - | ++ | - | - | +/- | Figure 7B and S12 |
| KO mice | Mouse tissues | WB | 17 kDa | + | +/- | + | - | - | - | Figure 7B and S12 |
| KO mice | Mouse tissues | WB | 15 kDa | - | - | - | - | - | - | Figure 7B and S12 |
| KO mice | Mouse tissues | WB | < 15 kDa | - | - | - | - | - | - | Figure 7B and S12 |

#### Figure S15

### B

| | | | | pSyn #64 | EP 1536Y | MJF-R13 | 81A | GTX | LASH-EGT | Figures |
|---|---|---|---|---|---|---|---|---|---|---|
| WT mice | PFFs-treated Primary neurons | ICC | Neurites | + | + | + | + | + | + | Figures 4A, 6 and S6-S7 |
| | | | Cell bodies | + | + | + | + | + | + | Figures 4A, 6 and S6-S7 |
| | | | Nucleus | + | - | + | - | - | - | Figure 6 |
| | | | Morphological diversity | - | Not tested | + | - | + | Not tested | Figures 6 and S7 |
| | | HTS | Background | ++ | +/- | ++ | +/- | +/- | +/- | Figures 4B-C and S6A |
| | | WB | HMWs > 25 kDa | + | - | ++ | + | - | - | Figures 4D and S6B |
| | | | 17 kDa (unspecific)* | + | + | + | + | + | + | Figures 4D and S6B |
| | | | 15 kDa | + | +/- | ++ | + | - | - | Figures 4A, 6 and S6-S7 |
| | | | < 15 kDa (unspecific)* | + | + | + | + | + | + | Figures 4A, 6 and S6-S7 |
| | PFFs-injected Mouse tissues | IHC | Neurites | + | ++ | + | ++ | +/- | + | Figure 8A and B |
| | | | Cell bodies | ++ | ++ | + | +/- | + | ++ | Figure 8A and B |
| | | | Nucleus | + | ++ | - | +/- | +/- | + | Figure 8A and B |
| | | WB | HMWs > 25 kDa | N.D | N.D | N.D | N.D | N.D | N.D | |
| | | | 17 kDa | N.D | N.D | N.D | N.D | N.D | N.D | |
| | | | 15 kDa | N.D | N.D | N.D | N.D | N.D | N.D | |
| | | | < 15 kDa | N.D | N.D | N.D | N.D | N.D | N.D | |

**Figure S15. pS219 background and specificity in the neuronal and animal seeding models (related to Figures 2-8)**

**A-B.** Tables recapitulating our main observations regarding the background and specificity of the six pS129 antibodies (pSyn#64, MJF-R13, 81A, EP1536Y, GTX and LASH-EGT) for ICC, HTS, IHC and WB applications in aSyn KO or PFFs-treated WT primary neurons or mice. * Detected both in the PBS- and the PFF-treated neurons. N.D for non-determined.
